# Genome duplications, genomic conflict, and rapid phenotypic evolution characterize the Cretaceous radiation of Fagales

**DOI:** 10.1101/2023.06.11.544004

**Authors:** Ying-Ying Yang, Gregory W. Stull, Xiao-Jian Qu, Lei Zhao, Yi Hu, Zhi-Heng Wang, Hong Ma, De-Zhu Li, Stephen A. Smith, Ting-Shuang Yi

**Author notes:** Author for corrspondnce: Ting-Shuang Yi. These authors contributed equally to this work (Ying-Ying Yang, Gregory W. Stull).

## Abstract

- Flowering plant lineages display remarkable diversity in life history and morphological form. However our understanding of how this phenotypic diversity or disparity, arises and what, if any, relation there is to processes such as gene and genome duplication remains unlcear.
- Here we investigate the relationship between phenotypic and genomic evolution in the angiosperm order Fagales, a lineage of woody plants that has been a dominant component of temperate and subtropical forests since the Late Cretaceous. To this end, we examine newly generated trait and transcriptomic datasets spanning the phylogenetic breadth of the order, including most extant genera as well as a rich diversity of Cretaceous fossil representatives.
- Our phylogenomic analyses resolved the location of an ancient polyploidization event in Juglandaceae and identified hotspots of gene-duplication and genomic conflict across the order. Examinations of phenotypic evolution (including morphospace occupancy and rate shifts) show that the morphospace occupied by Fagales was filled by the early Cenozoic and rates of evolution were highest during the early radiation of the Fagales crown and its major families. Many instances of phenotypic rates also correspond to hotspots of gene duplication.
- Our results show that Fagales conform to an “early burst” model of phenotypic diversification, with morphospace being filled early in the group’s evolutionary history. Our study is consistent with others showing a decoupling of species diversification with other biological processes (e.g., phenotypic and genomic evolution), underscoring the complexity of understanding the major drivers of plant evolution over broad timescales.

## Introduction

Flowering plant lineages display remarkable diversity in life history and phenotype, corresponding to broad variation in ecological setting (Soltis *et al*., 2018). Understanding how this phenotypic diversity, or disparity, arises over geologic timescales is a major goal of evolutionary biology. However, despite an abundance of studies on broad-scale patterns of species diversification in angiosperms (e.g., Crane & Lidgard, 1989; Magallón & Sanderson, 2001; Smith *et al*., 2011; Tank *et al*., 2015; Landis *et al*., 2018; Ramírez-Barahona *et al*., 2020) as well as many studies examining patterns of disparity through time in major plant lineages (e.g., Lupia, 1999; Boyce & Knoll, 2002; Crepet & Niklas, 2019), processes underlying disparification have been relatively poorly understood (Oyston *et al*., 2016). Diversification has often been examined as a potential correlate to disparity (e.g., Folk et al., 2019; Nürk et al. 2019; Walden et al. 2020). However, evidence generally suggests that there is no inherent relationship between phenotypic evolution (disparification or rates) and diversification in plants or other lineages (Foote, 1991; Fortey *et al*., 1996; Folk *et al*., 2019), but nevertheless additional studies on these phenomena in plants would be valuable for understanding the circumstances in which they tend to show concordant or dissimilar patterns.

Advances in sequencing over the past couple decades have improved our ability to study genomic correlates to shifts in disparity. Recent examinations of phenotypic evolution alongside phylogenomic analyses have shown that whole-genome duplication (WGD) often correspond to bursts of phenotypic innovation in plants (Smith et al. 2018; Clark & Donoghue, 2018; Guo *et al*., 2020; Sheehan *et al*., 2020; Walden *et al*., 2020; Stull *et al*., 2021), consistent with the long-held idea that gene-duplications (Ohno, 1970), and WGDs more broadly (Levin, 1983), contribute significantly to the emergence of evolutionary novelty. Recent studies have also highlighted that, in plants and other major lineages, major phases of phenotypic evolution often occur along phylogenetic branches also characterized by extensive levels of genomic conflict (McGee *et al*., 2020; Ronco *et al*., 2021; Parins-Fukichi *et al*., 2021), suggesting that coordinated analyses of phenotypic and genomic data can offer important insight on the processes underlying major phases of phenotypic evolution.

Here we examine broad-scale patterns and correlates of phenotypic and genomic evolution in the woody angiosperm clade Fagales, using newly generated trait and transcriptomic datasets collected from nearly all extant genera and a rich diversity of Cretaceous fossil representatives. Comprising seven families and ∼1300 species, Fagales represent one of the most ecologically important plant groups in the Northern Hemisphere, with numerous species of oaks (*Quercus* L.), hickories (*Carya* Nuttall), beeches (*Fagus* L.), and alder (*Alnus* Miller) acting as dominant trees in temperate and subtropical forests (Furlow, 1979; Manos & Stanford, 2001; Manos & Stone, 2001). Because of the exceptionally rich Cretaceous and Cenozoic fossil record of Fagales (Manchester, 1987; Crane, 1989; Friis *et al*., 2011; Xing *et al*., 2016), the clade has been treated as a model system for the integration of fossil evidence into plant evolutionary studies, with particular emphases on placing fossils in phylogenies (Manos *et al*., 2007), divergence times (Sauquet *et al*., 2012), species diversification (Larson-Johnson, 2016; Xiang *et al*., 2014; Xing *et al*., 2014), and biogeographic history (Zhang *et al*., 2022; Siniscalchi *et al*., 2023). Surprisingly less attention has been paid to understanding broad-scale patterns of phenotypic evolution in this group, although the morphology and anatomy of the clade has generally been well documented (e.g., Larson-Johnson, 2016; Wheeler *et al*., 2022). While Fagales are relatively uniform in several life history attributes (e.g., woody growth form; tendency for unisexual flowers and wind-pollination in extant members), they show considerable variation in wood anatomy (Wheeler *et al*., 2022), leaf structure (Hickey & Wolfe, 1975; Doyle, 2007), pollen morphology (Zavada & Dilcher, 1986), and diaspore structure and functional morphology (Larson-Johnson, 2016). Understanding when major phases of disparification in Fagales occurred in geologic history, and how these were related to genomic events (e.g., WGD) and broader diversification patterns, will provide new insights into the evolution of this major plant lineage as well as a window on how disparity arises in plant lineages at macroevolutionary scales.

We present a robust phylogenomic framework for Fagales based on a newly generated transcriptome dataset (including nearly 160 ingroup species), which we use to resolve outstanding relationships, infer and place WGD events, and characterize hotspots of gene duplications and genomic conflict, also use as a backbone for a supertree analysis of Fagales species diversification. To characterize major patterns of phenotypic rates and disparity across Fagales phylogeny, we generated a new phenotypic dataset, matched to the transcriptomic sampling, comprising 152 traits and including 52 fossil taxa (primarily from the mid/late Cretaceous to early Cenozoic). Our phenotypic dataset includes traits from all major organ systems, in contrast to most previous investigations of disparity in plants focusing on a single organ system (e.g., Lupia, 1999; Chartier *et al*., 2017; Kriebel *et al*., 2017; Mander *et al*., 2020; Jardine *et al*., 2022). A phylogenetic framework including extensive fossil diversity was leveraged to examine major patterns of morphological evolution through time. We synthesize these results to address the following questions: Does Fagales display an early-burst model of disparification, reflected by rapid filling of realized morphospace shortly after their origin in the Cretaceous? Do major shifts in phenotypic evolution (in terms of rates or disparity) typically coincide with ancient WGD events? Is phenotypic diversification generally uncoupled with species diversification across major lineages of Fagales?

## Materials and Methods

### Taxon sampling and phenotypic data

We sampled 182 species (including 158 Fagales species and 24 outgroup species representing major lineages of eudicots; see Table S1) for our transcriptomic and phenotypic datasets. All seven Fagales families were represented as well as 32 of the 34 genera of the order; we were unable to obtain fresh material of *Canacomyrica* and *Ceuthostoma* for transcriptome sequencing due to their limited distributions. Of the 183 transcriptomes included, 149 were newly generated for this study; the remainder were obtained from GenBank or previous studies (supplementary Table S1). The phenotypic dataset included all extant species from the transcriptome dataset and 52 fossil taxa. A total of 152 traits were scored for the phenotypic dataset (Text S1); the trait definitions and character states used in this study build on several previous studies examining particular organ types (e.g., Wheeler *et al*., 2022) or aiming to place fossils in Fagales phylogeny (Manos *et al*., 2007; Larson-Johnson, 2016). Trait data for our sampled species were obtained from these earlier studies (Larson-Johnson, 2016; Wheeler *et al*., 2022; Zhang *et al*., 2022), as well as from major floras (Flora of China, Flora of North America, Flora of Australia, Flora Europaea, Flora of Pakistan, Flora of Thailand), the primary literature (The families and genera of vascular plants, Notes on Casuarinaceae III), and websites (http://oaks.of.the.world.free.fr/index.htm, https://www.treesandshrubsonline.org/, https://en.wikipedia.org/).

### Transcriptome sequencing, assembly, and ortholog identification

To sequence the new transcriptomes, total RNAs were extracted from fresh young leaves, leaf buds, flower buds and/or fruit buds of most species, and silica-dried leaf materials of three remaining species (*Alfaroa guanacastensis* D.E.Stone, *Oreomunnea pterocarpa* Oerst., *Ticodendron incognitum* Gómez-Laur. & L.D.Gómez) using the SpectrumTM Plant Total RNA kit (Sigma-Aldrich), quantified using the NanoDrop 1000 Spectrophotometer (Thermo Scientific, Wilmington, DE). About 5ug RNAs were used to construct DNA libraries (NEBNext Ultra RNA Library Prep Kit for Illumina). Paired-end reads (150 bp) were sequenced using the Illumina HiSeq4000 platform by Novogene Corporation, Beijing, China, resulting in 6 Gb of transcriptome sequences per sample. The resulting reads were de-novo assembled using Trinity-v2.4.0 (Haas *et al*., 2013) with default parameters. Nonredundant transcripts were obtained using CD-HIT-EST (-c 0.99 –n 10) (Fu *et al*., 2012). TransDecoder v5.3.067 (Haas *et al*., 2013) was applied to predict amino acid sequences.

We used a hierarchical clustering procedure for homology identification following Walker *et al*. (2018) and Larson *et al*. (2020). We divided Fagales into five subclades according to results of previous phylogenetic studies (Xing *et al*., 2014; Xiang *et al*., 2014; Larson-Johnson, 2016): BTC, Betulaceae + Ticodendraceae + Casuarinaceae; JM, Juglandaceae + Myriaceae; FAGA, Fagaceae; NOTHO, Nothofagaceae; OUT, the outgroups. An all-by-all BLAST search (Altschul *et al*., 1997) with an e-value cutoff of 10 was performed on each group following Yang & Smith (2014) with minor modifications. The tip clusters were then sequentially combined in a post-order fashion (node clustering) using the scripts developed by Walker *et al*. (2018) (https://github.com/jfwalker/Clustering). A total of 9,955 homologue gene trees were obtained from the combined clusters after trimming long branches and filtering low species coverage clusters following scripts of Yang & Smith (2014) (https://bitbucket.org/yangya/ phylogenomic_dataset_construction/src/master/). Based on these homologue gene trees, 643 orthologues containing at least 110 taxa (i.e., ≥ 60% of the total sampling) were extracted using the rooted tree (RT) method of Yang & Smith (2014). For each orthologue, sequences were realigned using PRANK v.140110 (Löytynoja & Goldman, 2010) with default settings, alignment columns with low occupancy (≥ 70% missing data) were trimmed with ‘pxclsq’ function in Phyx (Brown *et al*., 2017), and only sequences with at least 110 species and 300 sites for each ortholog were retained for subsequent analyses.

### Species tree estimation and conflict analysis

Gene trees for all 643 orthologues were inferred using RAxML v8.1.22 (Stamatakis, 2014) with the GTR+Γ model and 200 fast bootstrap replicates. Species trees were inferred using two methods. First, a maximum likelihood tree (hereafter the ML tree) was inferred in RAxML v8.1.22 (Stamatakis, 2014) under the GTR+Γ model, using a concatenated matrix of the 643 ortholog alignments (with a total aligned length of 1,503,397 bp); node support was evaluated by 200 fast bootstrap replicates. Second, a maximum quartet support species tree (MQSST) was estimated in ASTRAL 5.6.3 (Mirarab *et al*., 2014) under default settings. Before the ASTRAL analysis, nodes with BS <10 in gene trees were collapsed using Newick utilities (Junier & Zdobnov, 2010). The ASTRAL analysis was conducted with default support values (local posterior probabilities) as well as support values from multilocus bootstrapping (Seo, 2008). The ML tree was used for most subsequent analyses including conflict analysis, inference of whole-genome duplications, and comparison of phenotypic innovation and rates of phenotypic evolution with levels of gene conflict and duplication.

We used PhyParts (Smith *et al*., 2015) to assess gene-tree conflict by mapping the 643 rooted gene trees onto the ML tree and summarizing the number of gene trees in concordance or conflict with each bipartition. Since the gene trees have different outgroups, trees were then rooted using a hierarchical strategy as listed in supplementary Table S2. We only considered nodes with >70% bootstrap support in the gene trees. The python script ‘phypartspiecharts.py’ (https://github.com/mossmatters/phyloscripts/tree/master/phypartspiecharts) was used to visualize the PhyParts results. The levels of conflict estimated for each bipartition were also plotted against levels of morphological innovation inferred below.

### Gene duplication mapping and WGD inference

A multifaceted approach was used to infer gene duplications and putative WGDs across Fagales, employing phylogenetic gene-duplication mapping, *Ks* plots of within-taxon paralogue pairs vs. between-taxon orthologue pairs, and reconstructions of chromosome number. For gene-tree mapping onto the species tree, only orthogroups with SH-like support > 80 were extracted from the homolog trees, as calculated using the SH-like test (Anisimova *et al*., 2011) in RAxML. In total, 11,555 rooted clusters with at least 60 taxa were extracted. We used PhyParts (Smith *et al*., 2015) to map those rooted clusters onto the inferred ML tree, with gene duplications recorded when sister clades shared two or more taxa (Yang *et al*., 2015).

*Ks* plots based on within-taxon paralogue-pairs were generated using the pipeline available at https://github.com/nstenz/plot-ks. Newly assembled contigs were translated using TRANSDECODER with a minimum protein length of 100 amino acids. Duplicate genes were identified using BLAT (Kent, 2002) based on translated contigs and then duplicate gene pairs were aligned and back translated into their corresponding nucleotide sequence. *Ks* values for each pair of nucleotide alignments were estimated using KAKS CALCULATOR (model GY; Zhang *et al*., 2006). For species showing *Ks* peaks suggestive of WGD, we double checked the results using scripts available at https://github.com/tan ghaibao/bio-pipeline/tree/master/synonymous_ calculation (results can be found in supplementary Fig. S7). Density plots were then generated using R (R core team, 2019) to reveal putative WGD events. We excluded *Ks* values < 0.05 to avoid isoforms from de novo assembled transcriptomes, and *Ks* values > 3 were also excluded to avoid the effects of *Ks* saturation. *Ks* plots for between-taxon orthologue pairs were generated following the method of Wang *et al*. (2019). We then compared orthologue and paralogue *Ks* peaks to determine whether observed peaks are shared between or among lineages (i.e., paralog peaks are older than orthologue peaks/lineage divergence) or instead are species or lineage specific (i.e., paralog peaks are younger than orthologue peaks/lineage divergence).

Chromosome counts were obtained from the Chromosome Counts Database (http://ccdb.tau.ac.il, accessed 13 June 2019). When different counts were reported for a species, we used the lowest recorded number. In general, a clade that has experienced a WGD event is expected to have twice as many chromosomes as their sister group. However, over evolutionary time, changes in chromosome number and diploidization are likely to make such distinctions less clear.

### Fossil placement and divergence-time estimation

We selected 52 fagalean fossil taxa for inclusion in phylogenetic analyses, dating analyses, and subsequent phenotypic reconstructions (Text S2). The fossils included are well-documented Cretaceous and Cenozoic taxa representing all major lineages of the order. We sought to include the oldest (and best-documented) fossil evidence for major Fagales lineages (Friis *et al*., 2011), to generate a robust dated tree for the group and to inform reconstructions of phenotypic evolution during the early diversification of Fagales. The oldest fossil included was *Soepadmoa cupulata* from the Turonian (93.9 to 89.8 million years ago), which represents the oldest reproductive structures of probable Fagales affinity (Gandolfo *et al*., 2018). See Text S2 for further details on the fossils included in this study.

We collected morphological data for 152 characters across 232 species (including the 52 fossils and 180 extant species of Fagales), for analysis in combination with the molecular data described above. Combined molecular-morphological analyses were conducted using maximum likelihood in RAxML to place the sampled fossils in Fagales phylogeny. The tree searches were broadly constrained to match the major relationships from the inferred species tree, and fossil placements were constrained to major clades based on current knowledge of their systematic affinities (see Fig S1 and text S2 for details on constraints used and supporting literature). Performing unconstrained analyses was untenable given the limited number of characters available for multiple fossils, which would have resulted in unstable placements if not constrained to major lineages. ML analyses included a search for the ML tree along with bootstrapping (using the command “-all”), with models partitioned by data type (morphological, MK+ASC; molecular, GTR+G) and branch lengths either linked or scaled across partitions.

Dating analyses were then conducted using an updated version of treePL (Stull *et al*., biorxiv) that allows fossils to be included as tips in the phylogeny. Both the fossil tip dates, and a root age were used as constraints on the ML tree from the combined analysis above (including both modern and fossil representatives), with smoothing parameters inferred via cross-validation analyses. Our outgroup sampling spanned the phylogenetic breadth of eudicots, and we applied a maximum age for the eudicot root (125 Ma) following Magallón *et al*. (2015). The ages derived from this treePL analysis of the combined extant/fossil tree we then used to constrain node ages on the ML supertree (both the best tree and 29 bootstrap trees, the latter to offer confidence intervals on divergence times), also using treePL (Smith & O’Meara, 2012). The ultrametric supertree was then used for subsequent analyses of diversification.

### Disparity and rates of phenotypic evolution

We conducted a suite of analyses to characterize patterns of phenotypic disparity and rate shifts in phenotypic evolution across the Fagales phylogeny. We examined morphospace-occupancy of Fagales (including and excluding fossil representatives) through time to provide a window on the timing of major phases of disparification across the order: was disparity accumulated gradually through time, or rapidly during the early diversification of the order? Specifically, the function ‘plot_chronophylomorphospace’ (Sakamoto & Ruta, 2012) from the R package Claddis (Lloyd, 2016) was used to generate plots showing changes in morphospace occupancy through time in a phylogenetic context. This is accomplished by estimating ancestral states on a provided tree, calculating morphological distances, and then performing a principal coordinates analysis (PCoA), which serves as the basis for plotting phylomorphospace through time (Lloyd, 2016).

Several approaches were used to identify major rate shifts in phenotypic evolution. Following Parins-Fukuchi *et al*. (2021), we used parsimony to determine the number of state changes per branch; we then divided the number of changes per branch by the branch duration (in absolute time, using the dated phylogeny), resulting in per-branch estimates of rates of phenotypic evolution. We also used the Mk model to calculate ML morphological branch lengths, which were similarly divided by their duration in absolute time to provide another estimate of per-branch phenotypic rates. Finally, we used Bayesian Analysis of Macroevolutionary Mixtures (BAMM) v2.5.0 (Rabosky, 2014) to estimate significant shifts in rates of phenotypic evolution across Fagales. Two separate BAMM analyses were conducted using the first two axes from a principal coordinates analysis (PCoA). These BAMM analyses employed the trait model, with priors determined using the function ‘setBAMMpriors’ in BAMMtools (Rabosky *et al*., 2014), which was also used to assess convergence of the run (i.e., whether the effective sample sizes of the number of shift events and the log-likelihoods in each sample were >200 after 10% burn-in).

Finally, we identified ‘jumps’ in phenotypic evolution (i.e., extreme changes in phenotypic values between parent and child nodes) across Fagales phylogeny, employing an approach similar to Smith *et al*. (2018). Two separate analyses were conducted using the first two axes from a PCoA of the morphological matrix. Each of these was used for ancestral state reconstruction; parent and child values were then compared, and differences in the 95th percentile were considered significant shifts in phenotype.

We conducted several analyses to examine the relationship between per-branch levels of phenotypic evolution (both rates and number of state changes) and per-branch levels of gene duplication and genomic conflict. For a visual comparison, we plotted phenotypic values (rate or innovation) against genomic values (duplications or conflict) for each node through time (the x axis). We then explicitly tested the relationships between aspects of phenotypic and genomic evolution using generalized linear regression of each pair of log-transformed variables. Generalized linear regression was performed using the ‘glm’ function in R, treating phenotypic rates or levels of phenotypic innovation as the dependent variables.

### Supermatrix analysis and species diversification

To reconstruct a comprehensive phylogeny of Fagales, we mined 35 publicly available plastid gene sequences of 769 species from NCBI using a cluster analysis in program PyPhlawd under default settings (Smith & Walker, 2018). Plastid gene sequences extracted from 42 plastomes (include two outgroup species) and 36 newly sequenced transcriptomes were also added to the supermatrix. We assessed synonymy among the names of included accessions and only one representative per accepted species name was retained (including in cases where species had several subspecies, varieties, or forms). The plastid genes were aligned individually using MAFFT v7 (Katoh & Standley, 2013), after which alignment columns with under 30% occupancy were removed using the phyx function ‘pxclsq’ (Brown *et al*., 2017). The alignments were then concatenated using the phyx function ‘pxcat’ (Brown *et al*., 2017). The final concatenated matrix included 44,168 sites and 847 species. Phylogenetic reconstruction was carried out using RAxML v8.1.22 (Stamatakis, 2014) with the GTR+Γ model, with 200 bootstraps. Topological constraints were used to ensure consistency with major relationships inferred from the transcriptome dataset (Fig S2), allowing greater continuity in interpretations across all analyses. The tree with the best likelihood score was selected for downstream analyses of diversification and climate occupancy evolution. We conducted another 29 independent tree reconstructions via bootstrapping of the concatenated supermatrix using the ‘-f j’ option in RAxML v8.1.22 (Stamatakis, 2015).

We used BAMM v2.5.0 (Rabosky, 2014) to analyze diversification rate shifts and climatic occupancy evolution across Fagales using the dated supertree. The species diversification analysis used the speciation-extinction model, with the priors determined using the function ‘setBAMMpriors’ in BAMMtools (Rabosky *et al*., 2014). To account for incomplete taxon sampling, we specified sampling fractions for each sampled genus. Convergence of the MCMC chains was confirmed as described previously.

The morphological matrices, molecular data, and scripts to perform morphospace and other downstream analyses used in this study are available at XXX/fagales.

## Results and Discussion

### Phylogenetic relationships and divergence-times

Both the ML and MQSST trees showed well–resolved and highly supported phylogenetic relationships, with nearly identical topologies except a few nodes (see Fig. S3–4). Nothofagaceae and Fagaceae were supported as successively sister to the core Fagales (i.e., the remainder of the order), and the core Fagales were resolved into the Betulaceae + Ticodendraceae + Casuarinaceae (BTC) clade, and the Juglandaceae + Myricaceae (JM) clade. Relationships among Nothofagaceae, Fagaceae, and the core Fagales and relationships within the BTC clade had been largely resolved in previous studies based on analyses of one or a few combined gene loci, largely from the chloroplast and mitochondrial genomes (Manos & Steele, 1997; Cook & Crisp, 2005; Sauquet *et al*., 2012; Xiang *et al*., 2014; Sun *et al*, 2016; Yang *et al*., 2021). However, the placement of Myricaceae has been controversial (Manos & Steele, 1997; Li *et al*., 2002; Xiang *et al*., 2014; Sun *et al*., 2016), although a sister relationship with Juglandaceae is most widely supported (Li *et al*., 2004; Soltis *et al*., 2007; Liu *et al*., 2017; Yang *et al*., 2021; Ding *et al*., 2023). A recent study integrating molecular and morphological data resolved Nothofagaceae and Fagaceae as sister to each other, rather than as successively sister to the rest of Fagales (Siniscalchi *et al*., 2023 biorxiv), but our results are consistent with a wealth of previous phylogenetic and phylogenomic studies (Manos & Steele, 1997; Cook & Crisp, 2005; Sauquet *et al*., 2012; Xiang *et al*., 2014; Sun *et al*., 2016; Yang *et al*., 2021) in showing these two families as successively sister to the rest of the order.

Our analyses present additional results that may help address several previously contentious infrafamilial relationships (Fig. S3–4). In Betulaceae, a sister relationship between *Alnus* and *Betula* L. (Betuloideae) supported by some previous studies (Chen & Li, 2004; Grimm & Renner, 2013) was confirmed here (BSML = 100% / BSMQSST = 1). It is worth noting that the branch subtending this relationship is noticeably short, which might explain why either *Alnus* (Li *et al*., 2004; Xing *et al*., 2014; Li *et al*., 2015) or *Betula* (Yang *et al*., 2021) has previously been recovered as sister to the rest of the family. Subfamily Juglandoideae of Juglandaceae, a clade comprising tribe Platycaryeae and tribe Juglandeae was strongly supported as sister to tribe Hicoreae (*Carya*) (BSML = 100% / BSMQSST = 1), consistent with several previous studies (Zhang *et al*, 2013; Xiang *et al*., 2014; Mu *et al*., 2020). Within tribe Juglandeae, the topology of (*Juglans*, (*Cyclocarya*, *Pterocarya*)) was fully supported (BSML = 100% / BSMQSST = 1), consistent with the results of Dong *et al*. (2017) and Mu *et al*. (2020). Intergeneric relationships within Engelhardtioideae were well resolved as (*Engelhardtia* Blume (*Alfaropsis* Iljinskaya (*Alfaroa* Standley + *Oreomunnea* Oersted))); we follow Iljinskaya (1993) in treating *Alfaropsis* as separate genus from *Engelhardtia*.

In Fagaceae, all genera were inferred to be monophyletic. The large quercoid clade was resolved into two fully supported clades, one comprising *Castanea* Miller and *Castanopsis* (D. Don) Spach (clade I), and another comprising *Lithocarpus* Blume, *Chrysolepis* Hjelmq., *Notholithocarpus* Manos, Cannon & S.H.Oh, and *Quercus* (clade II), consistent with Manos & Stanford (2001) and other more recent studies (Zhou *et al*., 2022; Liu *et al*., 2023). Conflict was observed regarding the relationships in the clade including *Lithocarpus, Chrysolepsis, Notholithocarpus*, and *Quercus*, but this is not surprising in light of recent evidence for ancient hybridization among these lineages (Zhou *et al*., 2022; Liu *et al*., 2023) as well as our results showing a hotspot of gene duplication in the ancestor of this lineage.

Within oaks, the two recognized subgenera (subgenus *Quercus* and subgenus *Cerris*) were each recovered as monophyletic, as were the seven (of eight) sections sampled for this study (we did not sample section *Ponticae*). Within subgenus *Cerris*, section *Cyclobalanopsis* was strongly supported as sister to a clade including section *Ilex* + section *Cerris* (BSML = 100% / BSMQSST = 99%). Within subgenus *Quercus*, section *Lobatae*, section *Protobalanus*, and section *Virentes* were supported as successive sister to section *Quercus*, consistent with Hipp *et al*. (2020).

With 52 extinct Fagales species included in divergence time estimation, our dated phylogeny comprising 32 extant and 36 extinct genera represents an excellent framework for examining evolutionary questions in Fagales. Our dating results showed that Fagales originated in the Early Cretaceous, with a stem age of 108.5 Ma and a crown age of 105 Ma (Fig. 1, Fig. S5). Crown ages for extant families range from 93 to 67 Ma: Juglandaceae (92.4 Ma), Fagaceae (89.8 Ma), Casuarinaceae (81.1 Ma), Betulaceae (78.2 Ma), Myricaceae (76.4 Ma), Ticodendraceae (73.7 Ma) and Nothofagaceae (67.0 Ma). Our inferred age for crown Fagales (103.4 Ma) is similar to that of Xiang *et al*. (105.5 Ma, 2014), Xing *et al*. (105.7 Ma, 2014), and Magallón *et al*. (102.8 Ma, 2015). We also found that most genera (28/32 sampled) had originated and begun to diversify by the Eocene-Oligocene boundry (33.9 Ma), consistent with the Late Eocene peak in generic diversity found by Xing *et al*. (2014). More details on divergence times of major lineages can be found in Table S3.

**Figure 1.**
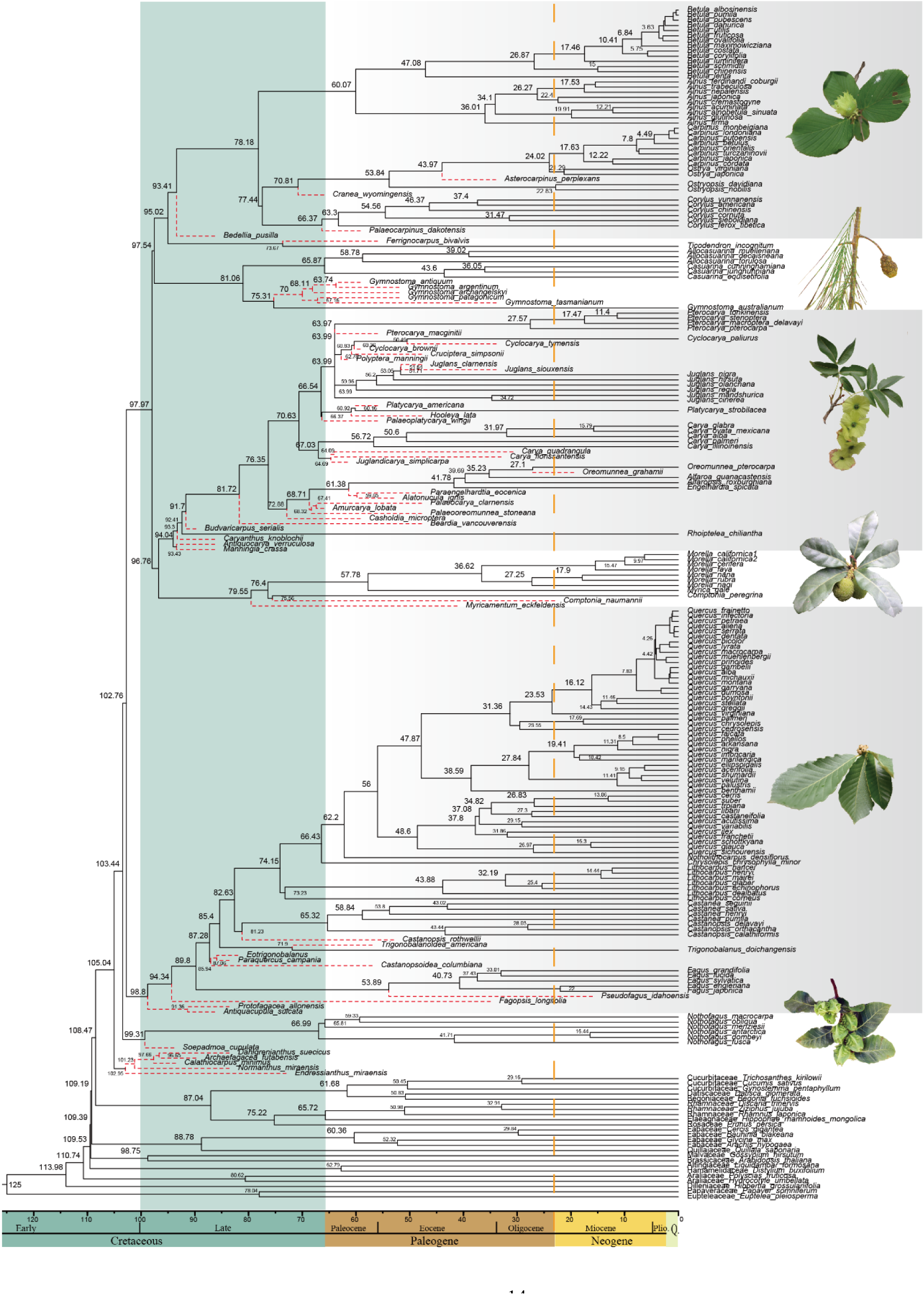
Dated phylogeny of Fagales including fossil evidence inferred using treePL. The input phylogeny, including 232 species (180 extant species; 52 fossil species shown by red dashed lines), was generated from combined molecular-morphological analyses. Results showed that Fagales ariose and began to diversify before the Late Cretaceous, while all seven extant families arose in the Late Cretaceous.

### Gene duplications and genomic conflict across Fagales

Our multifaceted approach for WGD inference showed one clear WGD event in Juglandaceae and multiple gene-duplication hotspots across Fagales (Fig. 2, Fig. S7). After examining 11,038 gene trees, a total of 636 duplicated genes (5.8%) were detected in the branch subtending the Juglandaceae crown, and 2348 duplicated genes (21.3%) were detected in the branch following the divergence of *Rhoiptelea chiliantha*. All the species of Juglandaceae shared a very distinct peak around *Ks* = 0.3 based on two methods of calculation (except for *Rhoiptelea* in one *Ks* analysis, Fig. S8-9), indicating a shared WGD in the family. Although the *Ks* peak at 0.3 in *Rhoiptelea* was less clear, between-taxon *Ks* plots showed that duplication events were earlier than the separation of *Rhoiptelea chiliantha* from Juglandaceae (Fig. S10). So, it is likely that this WGD event happened before the diversification of crown Juglandaceae, with greater losses of duplicate genes in *Rhoiptelea*. This conclusion is also supported by changes in base chromosome numbers: counts of Juglandaceae (n = 16) are twice that of its sister group Myricaceae (n = 8) (Fig. S11). Previously, it was unclear if this WGD occurred in the most recent common ancestor of the entire family (Luo *et al*., 2015; Griesmann *et al*., 2018; One Thousand Plant Transcriptomes Initiative, 2019), or after the divergence of *Rhoiptelea*. Our study complements the recent study by Ding *et al*. (2023), which used genome-synteny analyses, in showing that this ancient WGD is shared across all extant Juglandaceae.

**Figure 2.**
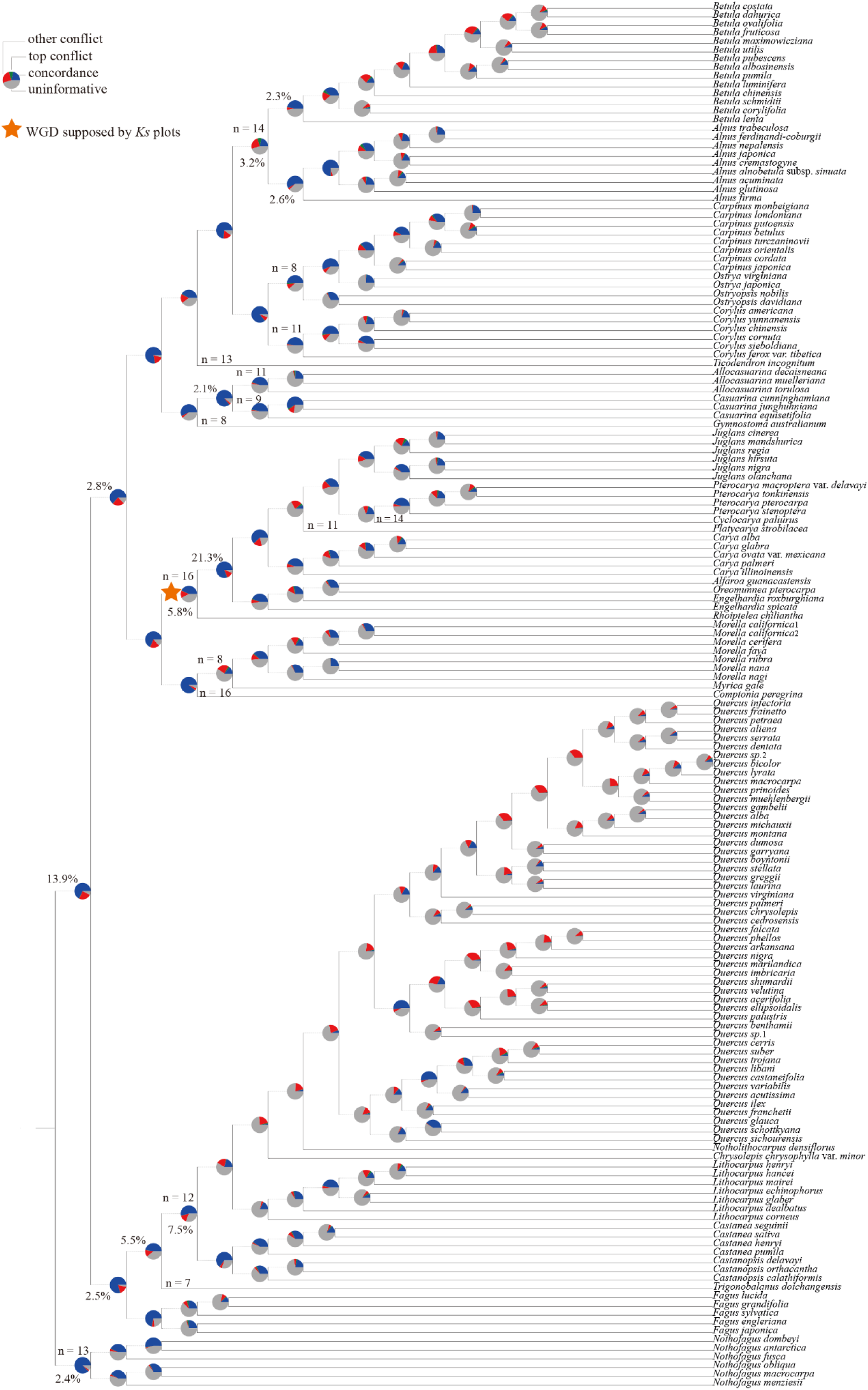
Phylogenetic tree showing gene conflicts and duplications. Species tree inferred from the 643 orthologues using RAxML. Pie charts at each node show the proportion of gene trees in concordance (blue), in conflict (green, main alternative; red, other conflicting topologies), or without information for that particular bipartition (grey). Nodes with percentages of duplicate genes are highlighted in the inset plot in the top/bottom left corner; Orange stars indicate WGD events supported by Ks plots; Changes in the haploid chromosome number are indicated on the phylogeny.

In addition to this WGD event, at least 10 gene-duplication hotspots (GDs) were detected across Fagales (Fig. 2, Fig. S7). The branch subtending Fagaceae + core Fagales included 1534 duplicated genes (13.9%), and a small percentage of gene duplications were mapped to the core Fagales (309, 2.8%). In Fagaceae, there were 604 (5.5%) duplicated genes on the branch subtending Quercoideae, with 828 (7.5%) on the branch subtending the large quercoid clade (*Quercus*, *Notholithocarpus*, *Chrysolepis*, *Lithocarpus*, *Castanopsis*, and *Castanea*). In Betulaceae, 351 duplicated genes (3.2%) were detected at the crown of the family, with 290 duplicated genes (2.6%) in the branch subtending Coryloideae, and 250 duplicated genes (2.3%) in the branch subtending *Betula*. In Casuarinaceae, 234 duplicated genes (2.1%) were detected in the branch subtending *Allocasuarina* L.A.S. Johnson and *Casuarina* L.. We also observed changes in chromosome number for several major clades, including Betulaceae (n = 8 to n = 14), Casuarinaceae (n = 8 to n = 11), and Fagaceae (n = 11 to n = 12) (Fig. 2, Fig. S11).

Significant gene tree conflict was observed across the phylogeny of Quercoideae (Fagaceae), especially in the quercoid clade (*Quercus*, *Notholithocarpus*, *Chrysolepis*, *Lithocarpus*), which has experienced rapid diversification and widespread gene flow (see Fig. 2, Fig. S6 and Zhou *et al*., 2022; Liu *et al*., 2023). Many conflicting gene trees were also identified concerning intergeneric relationships of Betuloideae (involving *Alnus* and *Betula*) and Juglandoideae (involving *Platycarya*, *Cyclocarya*, *Pterocarya* and *Juglans*). Conflict can be the result of biological processes such as ILS, hybridization, and horizontal gene transfer, and a growing body of literature suggests that hybridization has been common in Fagaceae, Betulaceae and Juglandaceae, even among the ancestors of modern genera during their initial radiation (Wang *et al*., 2016; Ding *et al*., 2023; Liu *et al*., 2023).

### Patterns of phenotypic evolution across Fagales

Phylogenetic reconstructions of morphospace occupancy through time show that Fagales and major families demonstrate an ‘early-burst’ model of disparification, given that the totality of morphospace occupied by the order was more or less filled early on in its evolution history, coinciding with family-level divergences (Fig. 3, Fig. S13-15). This is particularly clear when examining the morphospace occupied by early fossil representatives of Fagales, which are found at the extremities of morphospace. Although the first two principal coordinates account for < 5% of the variance in the dataset, this is a common property of morphospace analyses of discrete characters given the substantial number of principal coordinates (Lloyd, 2016). Major families (e.g., Betulaceae, Fagaceae, and Juglandaceae) show a similar pattern to Fagales as a whole where their morphospace is filled relatively early in their evolutionary history (Fig. 3).

**Figure 3.**
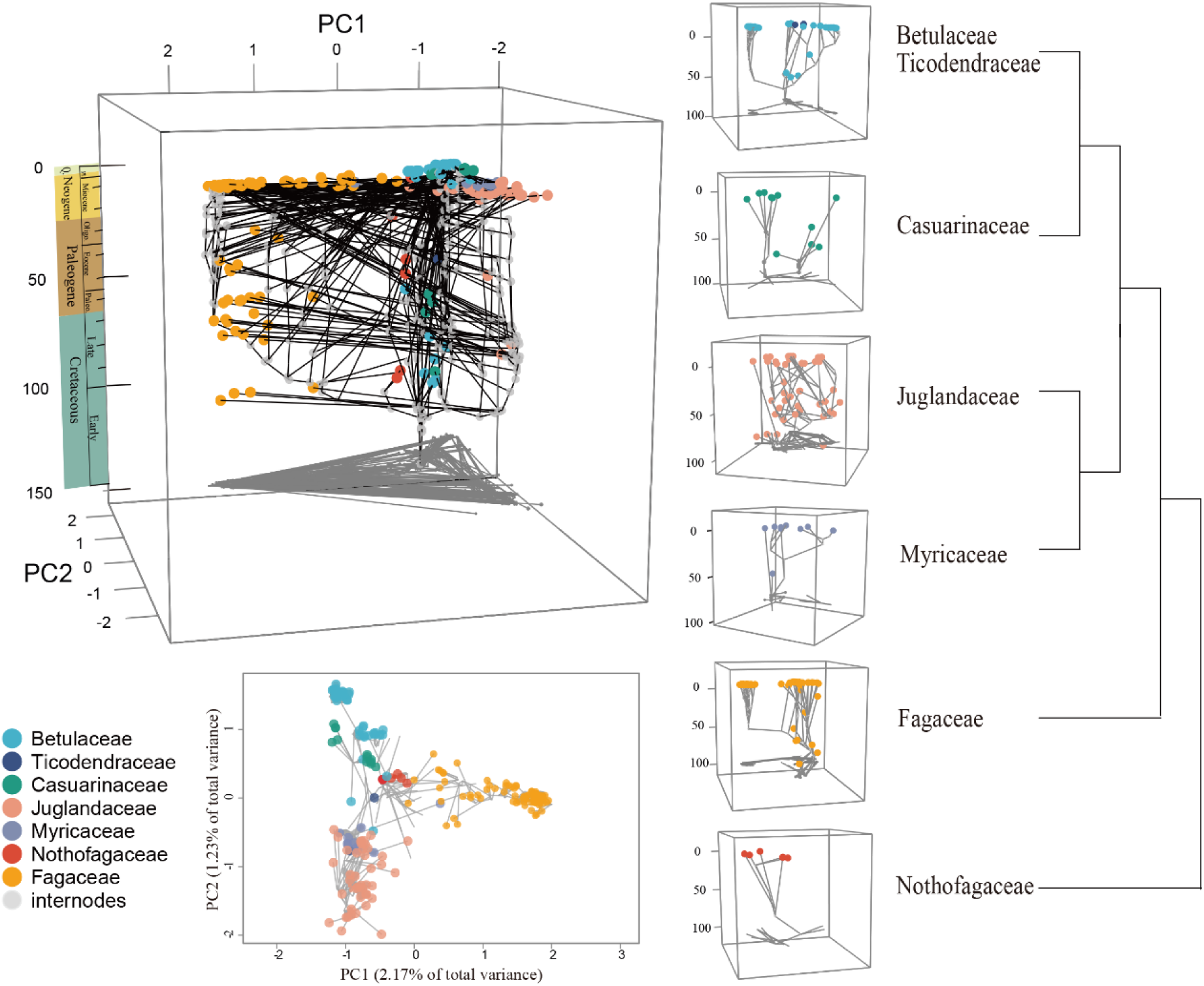
Chronophylomorphospace of Fagales (left) and all families (right) with both extant and extinct species included. (n=232). The top left plot supports an ‘early-burst’ model of disparification in Fagales.

The stem and crown ages of Fagales date to 108 and 103 mya, respectively. The subsequent lineage diversification of crown Fagales, from ∼103 to 90 mya, was accompanied by major expansions in morphospace. A further expansion in morphospace (in major extant lineages) is observed around the late Cretaceous to early Cenozoic, which represents the maximal extent of realized morphospace in the order (i.e., extant lineages fall within morphospace realized by the early Cenozoic).

The distribution of phenotypic rates and innovation (based on inferred phenotypic branch lengths) showed the greatest morphological change along many of the deepest branches of Fagales, which are confined to a narrow temporal window, indicating rapid phenotypic change (Fig. 4, Fig. S16). The BAMM analysis showed that phenotypic rates were fastest during the initial radiation of the clade, with major Fagales lineages showing uniform rates following the Cretaceous radiation (Fig. S17). Estimates of phenotypic rates based on the ML and parsimony approaches (following Parins-Fukuchi *et al*., 2021) similarly showed the greatest rates along the backbone of Fagales. Major evolutionary ‘jumps’ (i.e., significant parent-to-child changes in phenotype, which provides a window on phylogenetic locations of major phenotypic changes) are also generally found on deeper branches, including at the crown of core Fagales and at or near the crowns of most families (Fagaceae, Myricaceae, Jugladaceae, Betulaceae); major clades within Fagales also are characterized by evolutionary jumps in phenotype (Fig. S16).

**Figure 4.**
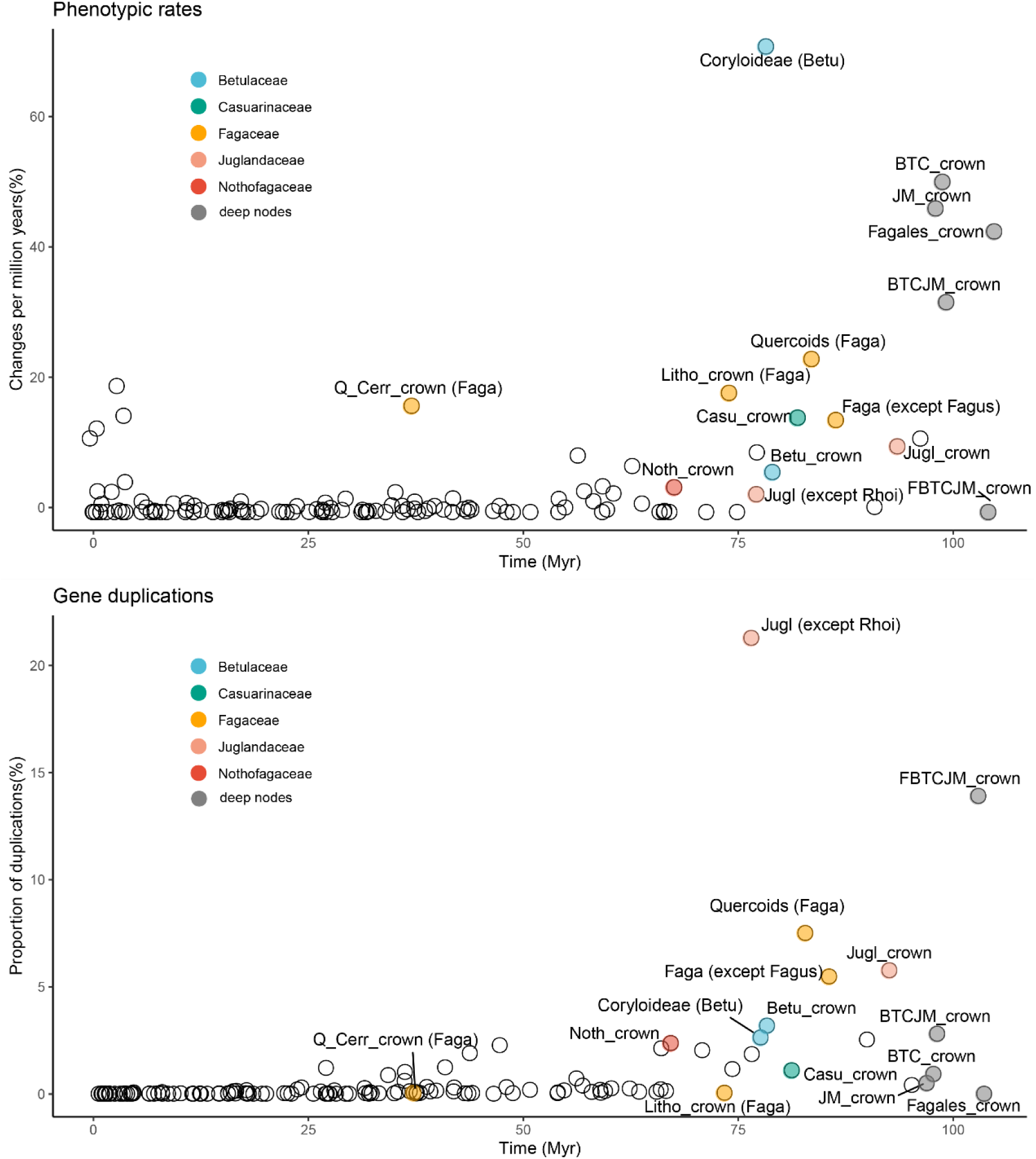
Rates of phenotypic innovation versus gene duplications. Plots juxtaposing branch-specific phenotypic rates (the number of state changes per branch divided by the branch duration in millions of years) and gene duplication at corresponding nodes across Fagales phylogeny (n = 143). The x axis represents time in millions of years. Each circle represents a branch in the phylogeny. Outlier branches are highlighted with colored circles. Betu, Betulaceae; Casu, Casuarinaceae; Faga, Fagaceae; Jugl, Juglandaceae; Noth, Nothofagaceae; BTC, the Betulaceae + Ticodendraceae + Casuarinaceae clade; JM, the Juglandaceae + Myricaceae clade; BTCJM, the core Fagales; FBTCJM, Fagales except Nothofagaceae; Litho, *Lithocarpus* of Fagaceae; Q_Cerr (Faga), *Quercus* section *Cerris* of Fagaceae; Faga (except Fagus), the Fagaceae except Fagus; Jugl (except Rhoi), Juglandaceae except *Rhoiptelea*; crown, the crown node of the lineage.

### Relationship of phenotypic and genomic evolution and diversification

Overall, we find a positive relationship between rates of phenotypic evolution and gene duplication (linear regression: r^2^ = 0.1786, P ≈ 0.000, Fig. S18), suggesting that gene duplications may have played a vital role in the phenotypic evolution of Fagales, consistent with other studies examining similar relationships (e.g., Stull *et al*., 2021, Jardine *et al*., 2022) and the general observation that neofunctionalization of duplicate genes derived fron GD and WGD events can help precipitate radical or/or rapid phenotypic innovations (Rensing, 2014). The branches with the most dramatic phenotypic changes are concentrated in Quercoideae (Fagaceae), Juglandaceae, and Betulaceae, where we also see particularly elevated levels of gene duplication. It has been observed that elevated rates of phenotypic innovation often coincide with the emergence of major lineages (Parins-Fukuchi *et al*., 2021). In our analyses, we observed that elevated phenotypic rates and gene duplication hotspots tended to both occur deeper in Fagales phylogeny, corresponding to the origin of major clades and named lineages (such as families or genera). While the relationship between these phenomena was not perfect, it is noteworthy that several of the observed phenotypic ‘jumps’ also coincide (or closely follow) with the inferred WGD and GD events—for example, at the crown/near of Betulaceaee, near the crown of core Fagales, at the branch subtending Quercoideae, and near the crown of Juglandaceae (Fig. S16).

Our diversification analysis of Fagales detected 11 major shifts, including shifts on the branches leading to *Alnus* + *Betula* (Betuloideae), *Carpinus* L. + *Ostrya* Scopoli, *Quercus*, and lineages within *Allocasuarina*, *Carya*, *Castanopsis*, *Morella* and *Lithocarpus* (Fig. S12). None of these shifts show a direct corresponding to branches with elevated levels of gene duplication. Our results are consistent with recent studies showing an decoupled relationship between WGD/GD and species diversification (Smith *et al*., 2018; Stull *et al*., 2021), suggesting that other biological attributes or ecological processes play a more direct role in shaping diversification dynamics in Fagales. Families with higher diversification rates did tend to show greater levels of gene tree conflicts (Fig. S20), but this is to be expected given that successive rapid speciation events will tend to result in greater ILS and, by extension, phylogenomic conflict.

## Acknowledgements

We thank the Germplasm Bank of Wild Species at the Kunming Institute of Botany (KIB) and Dr. Hong Ma’s lab in Department of Biology at University of Pennsylvania for facilitating this study; the curators and staff of Kunming Botany Garden (BG), Arnold Arboretum, Brisbane BG, Royal BG Edinburgh, Missouri BG, UC Berkeley BG, Xishuangbanna Tropical BG, and Dr. Min Deng (Yunnan University), Dr. Xun Gong (KIB), Dr. Jie Cai (KIB) for samples. This work was funded by the National Natural Science Foundation of China [key international (regional) cooperative research project No. 31720103903], the Strategic Priority Research Program of the Chinese Academy of Sciences (CAS) [grant No. XDB31000000], and the Science and Technology Basic Resources Investigation Program of China [2019FY100900]. G.W.S. acknowledges support from the CAS President’s International Fellowship Initiative (no. 2020PB0009) and the China Postdoctoral Science Foundation (CPSF) International Postdoctoral Exchange Program. S. A. S. acknowledges support from NSF DEB 1917146 and NSF DBI 1930030.

## Author contributions

TS Yi, DZ Li, SA Smith, YY Yang and GW Stull designed the study. YY Yang, XJ Qu, TS Yi, Y Hu collected and prepared samples for transcriptome sequencing; YY Yang, GW Stull, L Zhao generated the trait dataset and molecular datasets, and conducted analyses; GW Stull and YY Yang rote the manuscript, with contributions from TS Yi, SA Smith, ZH Wang, H Ma, and DZ Li. YY Yang and GW Stull contributed equally to this work.

## References

1. Altschul SF, Madden TL, Schäffer AA, Zhang J, Zhang Z, Miller W, Lipman DJ. 1997. Gapped BLAST and PSI-BLAST: a new generation of protein database search programs. Nucleic Acids Research 25: 3389–3402.

2. Anisimova M, Gil M, Dufayard JF, Dessimoz C, Gascuel O. 2011. Survey of branch support methods demonstrates accuracy, power, and robustness of fast likelihood-based approximation schemes. Systematic Biology 60: 685–699.

3. Boyce CK, Knoll AH. 2002. Evolution of developmental potential and the mul-tiple independent origins of leaves in Paleozoic vascular plants. Paleobiology 28: 70–100.

4. Brown JW, Walker JF, Smith SA. 2017. Phyx: phylogenetic tools for unix. Bioinformatics 33: 1886–1888.

5. Chen ZD, Li JH. 2004. Phylogenetics and biogeography of Alnus (Betulaceae) inferred from sequences of nuclear ribosomal DNA ITS region. International Journal of Plant Sciences 165: 325–335.

6. Chartier M, Lofstrand S, von Balthazar M, Gerber S, Jabbour F, Sauquet H, Schonenberger J. 2017. How (much) do flowers vary? Unbalanced disparity among flower functional modules and a mosaic pattern of morphospace occupation in the order Ericales. Proceedings of the Royal Society B: Biological Sciences 284: 20170066.

7. Clark JW, Donoghue PCJ. 2018. Whole-genome duplication and plant macroevolution. Trends in Plant Science 23: 933–945.

8. Cook LG, Crisp MD. 2005. Not so ancient: the extant crown group of Nothofagus represents a post-Gondwanan radiation. Proceedings of the Royal Society B: Biological Sciences 272: 2535–2544.

9. Crane PR. 1989. Early fossil history and evolution of the Betulaceae. Evolution, Systematics, and Fossil History of the Hamamelidae. Volume 2.“Higher Hamamelidae.” Systematic Association 40: 87–116.

10. Crane PR, Lidgard S. 1989. Angiosperm diversification and paleolatitudinal gradients in Cretaceous floristic diversity. Science 246: 675–678.

11. Crepet WL, Niklas KJ. 2019. The evolution of early vascular plant complexity. International Journal of Plant Sciences 180: 800–810.

12. Ding YM, Pang XX, Cao Y, Zhang WP, Renner SS, Zhang DY, Bai WN. 2023. Genome structure-based Juglandaceae phylogenies contradict alignment-based phylogenies and substitution rates vary with DNA repair genes. Nature communications 14, 617.

13. Dong W, Xu C, Li W, Xie X, Lu Y, Liu Y, Jin X, Suo Z. 2017. Phylogenetic resolution in Juglans based on complete chloroplast genomes and nuclear DNA sequences. Frontiers in Plant Science 8: 1148.

14. Doyle JA. 2007. Systematic value and evolution of leaf architecture across the angiosperms in light of molecular phylogenetic analyses. CFS Courier Forschungsinstitut Senckenberg 258: 21–37.

15. Folk RA, Stubbs RL, Mort ME, Cellinese N, Allen JM, Soltis PS, Soltis DE, Guralnick RP. 2019. Rates of niche and phenotype evolution lag behind diversification in a temperate radiation. Proceedings of the National Academy of Sciences USA 116: 10874–10882.

16. Foote M. 1991. Morphological and taxonomic diversity in clade’s history: the blastoid record and stochastic simulations. Contributions from the Museum of Paleontology 28: 101–140.

17. Foote M. 1992. Paleozoic record of morphological diversity in blastozoan echinoderms. Proceedings of the National Academy of Sciences USA 89: 7325–7329.

18. Foote M. 1994. Morphological disparity in Ordovician–Devonian crinoids and the early saturation of morphological space. Paleobiology 20: 320–344.

19. Fortey RA, Briggs DEG, Wills MA. 1996. The Cambrian evolutionary ‘explosion’: decoupling cladogenesis from morphological disparity. Biological Journal of the Linnean Society 57: 13–33.

20. Friis EM, Crane PR, Pedersen KR. 2011. Early flowers and angiosperm evolution. Cambridge, UK & New York, NY, USA: Cambridge University Press.

21. Fu L, Niu B, Zhu Z, Wu S, Li W. 2012. CD-HIT: accelerated for clustering the next-generation sequencing data. Bioinformatic 28: 3150–3152.

22. Furlow JJ. 1979. The systematics of the America species of Alnus (Betulaceae). Rhodora 81: 1– 121.

23. Gandolfo MA, Nixon KC, Crepet WL, Grimaldi DA. 2018. A late Cretaceous fagalean inflorescence preserved in amber from New Jersey. American Journal of Botany 105:1424– 1435.

24. Guo J, Xu W, Hu Y, Huang J, Zhao Y, Zhang L, Huang CH, Ma H. 2020. Phylotranscriptomics in Cucurbitaceae reveal multiple whole-genome duplications and key morphological and molecular innovations. Molecular Plant 13: 1117–1133.

25. Griesmann M, Chang Y, Liu X, Song Y, Haberer G, Crook MB, Billault-Penneteau B, Lauressergues D, Keller J, Imanishi L, et al. 2018. Phylogenomics reveals multiple losses of nitrogen-fixing root nodule symbiosis. Science 361: eaat1743.

26. Grimm GW, Renner SS. 2013. Harvesting Betulaceae sequences from GenBank to generate a new chronogram for the family. Botanical Journal of the Linnean Society 172: 465–477.

27. Haas BJ, Papanicolaou A, Yassour M, Grabherr M, Blood PD, Bowden J, Couger MB, Eccles D, Li B, Lieber M, et al. 2013. De novo transcript sequence reconstruction from RNA-seq using the Trinity platform for reference generation and analysis. Nature Protocols 8: 1494–512.

28. Hickey LJ, Wolfe JA. 1975. The bases of angiosperm phylogeny: vegetative morphology. Annals of the Missouri Botanical Garden: 538–589.

29. Hipp AL, Manos PS, Hahn M, Avishai M, Bodénès C, Cavender-Bares J, Crowl AA, Deng M, Denk T, Fitz-Gibbon S, et al. 2019. Genomic landscape of the global oak phylogeny. New Phytologist 226: 1198–1212.

30. Hughes CE, Atchison GW. 2015. The ubiquity of alpine plant radiations: from the Andes to the Hengduan Mountains. New Phytologist 207: 275–282.

31. Hughes M, Gerber S, Wills MA. 2013. Clades reach highest morphological disparity early in their evolution. Proceedings of the National Academy of Sciences USA 110: 13875–13879.

32. Iljinskaya, IA. 1993. Alfaropsis, a new genus of the Juglandaceae. Botanicheskii Zhurnal 78: 79–83.

33. Jardine PE, Palazzesi L, Telleria MC, Barreda VD. 2022. Why does pollen morphology vary? Evolutionary dynamics and morphospace occupation in the largest angiosperm order (Asterales). New Phytologist 234: 1075–1087.

34. Junier T, Zdobnov EM. 2010. The Newick utilities: high-throughput phylogenetic tree processing in the UNIX shell. Bioinformatics 26, 1669–1670.

35. Katoh K, Standley DM. 2013. MAFFT multiple sequence alignment software version 7: improvements in performance and usability. Molecular Biology and Evolution 30: 772–780.

36. Kent WJ. 2002. BLAT--the BLAST-like alignment tool. Genome Research 12: 656–664.

37. Kriebel R, Khabbazian M, Sytsma KJ. 2017. A continuous morphological approach to study the evolution of pollen in a phylogenetic context: an example with the order Myrtales. PLoS ONE 12: e0187228.

38. Landis JB, Soltis DE, Li Z, Marx HE, Barker MS, Tank DC, Soltis PS. 2018. Impact of whole-genome duplication events on diversification rates in angiosperms. American Journal of Botany 105: 348–363.

39. Larson-Johnson K. 2016. Phylogenetic investigation of the complex evolutionary history of dispersal mode and diversification rates across living and fossil Fagales. New Phytologist 209: 418–435.

40. Larson DA, Walker JF, Vargas OM, Smith SA. 2020. A consensus phylogenomic approach highlights paleopolyploid and rapid radiation in the history of Ericales. American Journal of Botany 107: 773–789.

41. Levin DA. 1983. Polyploidy and novelty in flowering plants. American Naturalist 122: 1–25.

42. Li HL, Wang W, Mortimer PE, Li RQ, Li DZ, Hyde KD, Xu JC, Soltis DE, Chen ZD. 2015. Large-scale phylogenetic analyses reveal multiple gains of actinorhizal nitrogen-fixing symbioses in angiosperms associated with climate change. Scientific Reports 5: 14023.

43. Li RQ, Chen ZD, Hong YP, Lu AM. 2002. Phylogenetic relationships of the “higher” hamamelids based on chloroplast trnL-F sequences. Acta Botanica Sinica 44: 1462–1468.

44. Li RQ, Chen ZD, Lu AM, Soltis DE, Soltis PS, Manos PS. 2004. Phylogenetic relationships in Fagales based on DNA sequences from three genomes. International Journal of Plant Sciences 165: 311–324.

45. Liu LX, Li R, Worth JRP, Li X, Li P, Cameron KM, Fu CX. 2017. The complete chloroplast genome of Chinese bayberry (Morella rubra, Myricaceae): Implications for understanding the evolution of Fagales. Frontiers in Plant Science 8: 968.

46. Liu SY, Yang YY, Tian Q, Yang ZY, Li SF, Valdes PJ, Farnsworth A, Kates HR, Siniscalchi CM, Guralnick RP, Soltis DE, Soltis PS, Stull GW, Folk RA, Yi TS. 2023. Phylogenomic analyses reveal widespread gene flow during the early radiation of oaks and relatives (Fagaceae: Quercoideae). bioRxiv doi: https://doi.org/10.1101/2023.04.25.538215.

47. Lloyd GT. 2016. Estimating morphological diversity and tempo with discrete character-taxon matrices: implementation, challenges, progress, and future directions. Biological Journal of the Linnean Society 118: 131–151.

48. Löytynoja A., Goldman N. 2010. webPRANK: a phylogeny-aware multiple sequence aligner with interactive alignment browser. BMC Bioinformatics 11, 579.

49. Luo MC, You FM, Li P, Wang JR, Zhu T, Dandekar AM, Leslie CA, Aradhya M, McGuire PE, Dvorak J. 2015. Synteny analysis in Rosids with a walnut physical map reveals slow genome evolution in long-lived woody perennials. BMC Genomics 16: 707.

50. Lupia R. 1999. Discordant morphological disparity and taxonomic diversity during the Cretaceous angiosperm radiation: north American pollen record. Paleobiology 25: 1–28.

51. Magallón S, Gómez-Acevedo S, Sánchez-Reyes LL, Hernández-Hernández T. 2015. A metacalibrated time-tree documents the early rise of flowering plant phylogenetic diversity. New Phytologist 207: 437–453.

52. Magallón S, Sanderson MJ. 2001. Absolute diversification rates in angiosperm clades. Evolution 55: 1762–1780.

53. Manchester SR. 1987. The Fossil History of the Juglandaceae. Monographs in Systematic Botany, Volume 21 (Missouri Botanical Garden). Lawrence, KS: Allen Press.

54. Mander L, Parins-Fukuchi C, Dick CW, Punyasena SW, Jaramillo C. 2020. Phylogenetic and ecological correlates of pollen morphological diversity in a neotropical rainforest. Biotropica 53: 74–85.

55. Manos PS, Stanford AM. 2001. The historical biogeography of Fagaceae: tracking the tertiary history of temperate and subtropical forests of the Northern Hemisphere. International Journal of Plant Sciences 162: S77–S93.

56. Manos PS, Steele KP. 1997. Phylogenetic analyses of ’’higher’’ Hamamelididae based on plastid sequence data. American Journal of Botany 84: 1407–1419.

57. Manos PS, Stone DE. 2001. Evolution, phylogeny, and systematics of the Juglandaceae. Annals of the Missouri Botanical Garden 88: 231–269.

58. Manos PS, Soltis PS, Soltis DE, Manchester SR, Oh SH, Bell CD, Dilcher DL, Stone DE. 2007. Phylogeny of extant and fossil Juglandaceae inferred from the integration of molecular and morphological data sets. Systematic Biology 56: 412–430.

59. McGee MD, Borstein SR, Meier JI, Marques DA, Mwaiko S, Taabu A, Kishe MA, O’Meara B, Bruggmann R, Excoffier L, et al. 2020. The ecological and genomic basis of explosive adaptive radiation. Nature 586: 75–79.

60. Mirarab S, Reaz R, Bayzid MdS, Zimmermann T, Swenson MS, Warnow T. 2014. ASTRAL: genome-scale coalescent-based species tree estimation, Bioinformatics 30: i541– i548.

61. Mu XY, Tong L, Sun M, Zhu YX, Wen J, Lin QW, Liu B. 2020. Phylogeny and divergence time estimation of the walnut family (Juglandaceae) based on nuclear RAD-Seq and chloroplast genome data. Molecular Phylogenetics and Evolution 147: 106802.

62. Nürk NM, Atchison GW, Hughes CE. Island woodiness underpins accelerated disparification in plant radiations. New Phytologist 224: 518–531.

63. Ohno S. 1970. Evolution by Gene Duplication. Springer Verlag. One Thousand Plant Transcriptomes Initiative. 2019. One thousand plant transcriptomes and the phylogenomics of green plants. Nature 574: 679–685.

64. Oyston JW, Hughes M, Gerber S, Wills MA. 2016. Why should we investigate the morphological disparity of plant clades? Annals of Botany 117: 859–879.

65. Parins-Fukuchi C., Stull GW, Smith SA. 2021. Phylogenomic conflict coincides with rapid morphological innovation. Proceedings of the National Academy of Sciences USA 118: e2023058118.

66. R Core Team. 2019. R: A language and environment for statistical computing. R Foundation for Statistical Computing, Vienna, Austria. Website: https://www.R-project.org/.

67. Rabosky DL. 2014. Automatic detection of key innovations, rate shifts, and diversity-dependence on phylogenetic trees. PLoS One 9: e89543.

68. Rabosky DL, Donnellan SC, Grundler M, Lovette IJ. 2014. Analysis and visualization of complex macroevolutionary dynamics: an example from Australian scincid lizards. Systematic Biology 63: 610–627.

69. Ramírez-Barahona S, Sauquet H, Magallón S. 2020. The delayed and geographically heterogeneous diversification of flowering plant families. Nature Ecology and Evolution 4: 1232–1238.

70. Rensing SA. 2014. Gene duplication as a driver of plant morphogenetic evolution. Current Opinion in Plant Biology 17: 43–48

71. Ronco F, Matschiner M, Böhne A, Boila A, Büscher HH, El Taher A, Indermaur A, Malinsky M, Ricci V, Kahmen A., Jentoft S. 2021. Drivers and dynamics of a massive adaptive radiation in cichlid fishes. Nature 589: 76–81.

72. Sakamoto M, Ruta M. 2012. Convergence and divergence in the evolution of cat skulls: temporal and spatial patterns of morphological diversity. PLoS ONE 7: e39752.

73. Sauquet H, Ho SYW, Gandolfo MA, Jordan GJ, Wilf P, Cantrill DJ, Bayly MJ, Bromham L, Brown GK, Carpenter RJ, et al. 2012. Testing the Impact of Calibration on Molecular Divergence Times Using a Fossil-Rich Group: The Case of Nothofagus (Fagales). Systematic Biology 61: 289–313.

74. Seo TK. 2008. Calculating bootstrap probabilities of phylogeny using multilocus sequence data. Molecular Biology and Evolution 25: 960–971.

75. Sheehan H, Feng T, Walker-Hale N, Lopez-Nieves S, Pucker B, Guo R, Yim WC, Badgami R, Timoneda A, Zhao L, et al. 2020. Evolution of L-DOPA 4,5-dioxygenase activity allows for recurrent specialization to betalain pigmentation in Caryophyllales. New Phytologist 227: 914–929.

76. Siniscalchi CM, Correa-Narvaez J, Kates HR, Soltis DE, Soltis PS, Guralnick RP, Manchester SR, Folk RA. 2023. Fagalean phylogeny in a nutshell: Chronicling the diversification history of Fagales. bioRxiv doi: https://doi.org/10.1101/2023.03.06.531381.

77. Smith SA, Beaulieu JM, Stamatakis A, Donoghue MJ. 2011. Understanding angiosperm diversification using small and large phylogenetic trees. American Journal of Botany 98: 404–414.

78. Smith SA, Brown JW, Yang Y, Bruenn R, Drummond CP, Brockington SF, Walker JF, Last N, Douglas NA, Moore MJ. 2018. Disparity, diversity, and duplications in the Caryophyllales. New Phytologist 217: 836–854.

79. Smith SA, O’Meara BC. 2012. treePL: divergence time estimation using penalized likelihood for large phylogenies. Bioinformatics 28: 2689–2690.

80. Smith SA, Moore MJ, Brown JW, Yang Y. 2015. Analysis of phylogenomic datasets reveals conflict, concordance, and gene duplications with examples from animals and plants. BMC Evolutionary Biology 15, 150.

81. Smith SA, Brown JW, Yang Y, Bruenn R, Drummond CP, Brockington SF, et al. Disparity, diversity, and duplications in the Caryophyllales. New Phytologist 217: 836–854.

82. Smith SA, Walker JF. 2019. PyPHLAWD: a python tool for phylogenetic dataset construction. Methods in Ecology and Evolution 10: 104–108.

83. Soltis DE, Gitzendanner MA, Soltis PS. 2007. A 567-taxon data set for angiosperms: The challenges posed by Bayesian analyses of large data sets. International Journal of Plant Sciences 168: 137–157.

84. Soltis DE, Soltis PS, Endress P, Chase MW, Manchester SR, Judd W, Majure L, Mavrodiev E. 2018. Phylogeny and evolution of the angiosperms: revised and updated edition. University of Chicago Press.

85. Stamatakis A. 2014. RAxML version 8: a tool for phylogenetic analysis and post-analysis of large phylogenies. Bioinformatics 30: 1312–1313.

86. Stroud JT, Losos JB. 2016. Ecological opportunity and adaptive radiation. Annual Review of Ecology, Evolution, and Systematics 47: 507–532.

87. Stull GW, Qu XJ, Parins-Fukuchi C, Yang YY, Yang JB, Yang ZY, Hu Y, Ma H, Soltis PS, Soltis DE, et al. 2021. Gene duplications and phylogenomic conflict underlie major pulses of phenotypic evolution in gymnosperms. Nature Plants 7: 1015–1025.

88. Sun M, Naeem R, Su JX, Cao ZY, Burleigh JG, Soltis PS, Soltis DE, Chen ZD. 2016. Phylogeny of the Rosidae: a dense taxon sampling analysis. Journal of Systematics and Evolution 54: 363–391.

89. Tank DC, Eastman JM, Pennell MW, Soltis PS, Soltis DE, Hinchliff CE, Brown JW, Sessa EB, Harmon LJ. 2015. Nested radiations and the pulse of angiosperm diversification: increased diversification rates often follow whole genome duplications. New Phytologis 207 :454–467.

90. Walden N, German DA, Wolf EM, Kiefer M, Rigault P, Huang XC, Kiefer C, Schmickl R, Franzke A, Neuffer B, et al. 2020. Nested whole-genome duplications coincide with diversification and high morphological disparity in Brassicaceae. Nature Communications 11: 1–12.

91. Walker JF, Yang Y, Feng T, Timoneda A, Mikenas J, Hutchison V, Edwards C, Wang N, Ahluwalia S, Olivieri J, et al. 2018. From cacti to carnivores: improved phylotranscriptomic sampling and hierarchical homology inference provide further insight into the evolution of caryophyllales. American Journal of Botany 105: 446–462.

92. Wang N, Mcallister HA, Bartlett PR, Buggs RJA. 2016. Molecular phylogeny and genome size evolution of the genus Betula (Betulaceae). Annals of Botany 117: 1023–1035.

93. Wang N, Yang Y, Moore MJ, Brockington SF, Walker JF, Brown JW, Liang B, Feng T, Edwards C, Mikenas J, et al. 2019. Evolution of Portulacineae marked by gene tree conflict and gene family expansion associated with adaptation to harsh environments, Molecular Biology and Evolution 36: 112–126.

94. Wheeler EA, Baas P, Manchester SR. 2022. Wood anatomy of modern and fossil Fagales in relation to phylogenetic hypotheses, familial classification, and patterns of character evolution. International Journal of Plant Sciences 183: 61–86.

95. Wills MA. 1998. Crustacean disparity through the Phanerozoic: comparing morphological and stratigraphic data. Biological Journal of the Linnean Society 65: 455–500.

96. Xiang XG, Wang W, Li RQ, Lin L, Liu Y, Zhou ZK, Li ZY, Chen ZD. 2014. Large-scale phylogenetic analyses reveal fagalean diversification promoted by the interplay of diaspores and environments in the Paleogene. Perspectives in Plant Ecology, Evolution and Systematics 16: 101–110.

97. Xing Y. Onstein RE, Carter RJ, Stadler T, Peter Linder H. 2014. Fossils and a large molecular phylogeny show that the evolution of species richness, generic diversity, and turnover rates are disconnected. Evolution 68: 2821–2832.

98. Xing Y, Gandolfo MA, Onstein RE, Cantrill DJ, Jacobs BF, Jordan GJ, Lee DE, Popova S, Srivastava R, Su T, Vikulin SV. 2016. Testing the biases in the rich Cenozoic angiosperm macrofossil record. International Journal of Plant Sciences 177: 371–388.

99. Yang Y, Moore MJ, Brockington SF, Soltis DE, Wong GKS, Carpenter EJ, Zhang Y, Chen L, Yan Z, Xie Y, et al. 2015. Dissecting molecular evolution in the highly diverse plant clade Caryophyllales using transcriptome sequencing. Molecular Biology and Evolution 32: 2001–2014.

100. Yang Y, Smith SA. 2014. Orthology inference in nonmodel organisms using transcriptomes and low-coverage genomes: improving accuracy and matrix occupancy for phylogenomics. Molecular Biology and Evolution 31: 3081–3092.

101. Yang YY, Qu XJ, Zhang R, Stull GW, Yi TS. 2021. Plastid phylogenomic analyses of Fagales reveal signatures of conflict and ancient chloroplast capture. Molecular Phylogenetics and Evolution 163, 107232.

102. Zavada MS, Dilcher DL. 1986. Comparative pollen morphology and its relationship to phylogeny of pollen in the Hamamelidae. Annals of the Missouri Botanical Garden 73: 348– 381.

103. Zhang JB, Li RQ, Xiang XG, Manchester SR, Lin L, Wang W, Wen J, Chen ZD. 2013. Integrated fossil and molecular data reveal the biogeographic diversification of the eastern Asian-eastern North American disjunct hickory genus (Carya Nutt.). Plos One 8: e70449.

104. Zhang Z, Li J, Zhao XQ, Wang J, Wong GK, Yu J. 2006. KaKs_Calculator: calculating Ka and Ks through model selection and model averaging. Genomics Proteomics Bioinformatics 4: 259–263.

105. Zhang Q, Ree RH, Salamin N, Xing Y, Silvestro D. 2022. Fossil-informed models reveal a boreotropical origin and divergent evolutionary trajectories in the walnut family (Juglandaceae). Systematic Biology 71: 242–258.

106. Zhou BF, Yuan S, Crowl AA, Liang YY, Shi Y, Chen XY, An QQ, Kang M, Manos PS, Wang BS. 2022. Phylogenomic analyses highlight innovation and introgression in the continental radiations of Fagaceae across the Northern Hemisphere. Nature Communications 13, 1320.

