## Supplemental materials for "Genome duplications, genomic conflict, and rapid phenotypic evolution characterize the Cretaceous radiation of Fagales"

### 1. Supplementary information

#### Text S1. List of traits scored for the phenotypic reconstructions

The primary sources of trait data were floras (e.g., Flora of China, Flora of North America, Flora of Australia, Flora of Europe), the general literature (Larson-Johnson, 2016; Zhang et al., 2020; Zhang et al., 2022; Wheeler et al., 2022; etc.), and websites (http://oaksoftheworld.fr). Traits related to wood anatomy, secondary metabolites, and other conserved features (e.g., ovary position, presence of endosperms, germination type, etc.) were collected at the generic level and scored across all included species except in cases were variation was noted at the generic level.

1 Growth form: tree (0), shrub (1), herb (2)
2 Branchlet pith: solid (0), chambered (1)
3 Porosity: diffuse-porous (0), semi-ring-porous (1), ring-porous (2)
4 Vessels arranged in a distinct pattern: radial (0), diagonal (1), dendritic (2), not present (3)
5 Vessels grouping: radial multiples present (0), radial multiples of four or more common (1), more than 90% of vessels solitary (2)
6 Perforation plate type: scalariform (0), simple (1)
7 Intervessel pitting: opposite (0), alternate (1), scalariform (2)
8 Vessel-ray parenchyma pits: bordered and similar to intervessel pits (0), pits with reduced to simple borders (1)
9 Helical thickenings present in vessel elements: not present (0), present (1)
10 Mean vessel tangential diameter:≤50 μm (0),≤100 μm (1), 100–200 μm (2), ≥200 μm (3)
11 Vasicentric tracheids present: not present (0), present (1)
12 Fiber pits: indistinct (minutely bordered to simple) (0), distinctly bordered (1)
13 Axial parenchyma occurrence: diffuse (0), diffuse-in-aggregates (1), marginal (2), banded (3), scanty paratracheal (4), vasicentric (5), rare (6)
14 Exclusively uniseriate rays: not present (0), present (1)
15 Aggregate rays: not present (0), present (1)
16 Rays of two distinct sizes: not present (0), present (1)
17 Cellular composition: homocellular and composed of procumbent cells (0), heterocellular and composed of procumbent and square/upright cells (1)
18 Crystal presence and location: in ray parenchyma (0), in idioblasts (1), in axial parenchyma (2), not present (3)
19 Plant nitrogen-fixation capacity: Yes (0), No (1)
20 Terminal buds: present (0), absent (1)
21 Terminal buds 2: strictly naked (0), with bud scales (1)
22 Stipules: present (0), absent (1)
23 Leaves: evergreen (0), winter-deciduous (1)
24 Leaves: simple (0), compound, odd-pinnate (1), compound, even-pinnate (2)
25 Phyllotaxy: alternate (0), opposite (1), whorled (2)
26 Leaf petiole length (mm): <10 (0), 10–20 (1), 21–30 (2), >30 (3)
27 Leaf length(cm): <0.1 (0), 0.1–0.9 (1), 1–10 (2), 11–20 (3), >20 (4)
28 Leaf width(cm): <1 (0), 1–5 (1), 6–10 (2), >10 (3)

29 Leaflet petiole length(mm): 0–5 (0), ≥5 (1)
30 Leaflet length(cm): 0–5 (0), 5–10 (1), >10 (2)
31 Leaflet width(cm): 0–5 (0), >5 (1)
32 Leaf/leaflet margin: entire (0), toothed-simple (1), toothed-compound (2)
33 Leaf/leaflet lobing: absent (0), shallowly lobed (1), deeply lobed (2)
34 Serration spacing: one or fewer teeth per pair of secondary veins (0), two or more per pair of secondary veins (1)
35 Leaf/leaflet venation of primary veins: pinnate (0), palmate (1)
36 Leaf/leaflet venation of secondary veins: craspedodromous (0), camptodromous (1)
37 Number of secondary veins pairs on opposite sides of the leaf: ≤5 (0), 6–10 (1), 11–20 (2), >20 (3)
38 Leaf/leaflet pubescence: glabrous (0), tomentose/villous/pilose/pubescent/hirsute (1), pubescent only along veins (2)
39 Embedded glands on leaf/leaflet: present (0), absent (1)
40 Flowers: unisexual (0), bisexual (1)
41 Pollination mode: wind pollination/anemophilous (0), insect pollination/entomophilous (1)
42 Inflorescences: androgynous (0), monoecism (1), dioecism (2), andromonoecious, or androdioecious (3), gynomonocious (4), polygamomonoecious (5)
43 Arrangement of bisexual flowers: solitary (0), catkin/spike/racemose (1), headlike/fasciculate (2), panicle (3), cyme (4), umbel (5)
44 Inflorescences: unbranched (0), branched (1)
45 Flower bracts: absent (0), present (1)
46 Bracts: entire (0), 3-lobed (1)
47 Flower tepal number: none (0), one (1), two (2), three (3), four (4), five (5), six (6), seven (7), >7 (8)
48 Staminode in pistillate flowers: absent (0), present (1)
49 Pistillode in staminode floret: absent (0), present (1)
50 Arrangement of male flowers: solitary (0), catkin/spike/racemose (1), headlike/fascicle (2), panicle (3), cyme (4), umbel (5)
51 Staminate inflorescence stances: erect (0), pendulous (1)
52 Staminate inflorescence lengths (mm): ≤20(0), 20-50(1), 50–100(2), >100(3)
53 Staminate inflorescences: unbranched (0), branched (1)
54 Number of male flowers subtended by a bract: one (0), two (1), three (2), four (3), five (4), six-seven (5)
55 Staminate flower bracts: absent (0), present (1)
56 Staminate bracts: entire (0), 3-lobed (1)
57 Staminate flower bracteoles number: none (0), one (1), two (2), three (3), four (4), five (5)
58 Staminate flower tepal number: none (0), one (1), two (2), three (3), four (4), five (5), six (6), seven (7), >7 (8)
59 Stamen number in staminate/bisexual flowers: one (0), two (1), three (2), four (3), five (4), six (5), seven (6), eight (7), nine (8), ten and more (9)
60 Anther attachment: basifixed (0), dorsifixed (1)
61 Number of locules present in anther: 2-loculed, separate (0), 2-loculed, connate (1), four, separate (2), one (3)

62 Anther length (mm): <0.5 (0), 0.5–1 (1), >1(2)
63 Pollen size: <17 micrometers (0), 17–26 micrometers (1), >26 micrometers (2)
64 Pollen aperture number: three (0), four (1), five (2), more (3)
65 Pollen aperture shape: colpate (0), porate (1), colporate (2)
66 Pollen shape: prolate (0), spheroidal (1), oblate (2)
67 Pollen shape of equatorial outline: triangular (0), circular (1), polygonal (2)
68 Pollen microspinule ornamentation: in even geometric pattern (0), in rows on ridges (1), irregularly distributed (2), no microspinules (3)
69 Pollen plica: absent (0), present (1)
70 Pollen arcus: absent (0), present (1)
71 Pollen endexine thickening around pores: absent (0), present (1)
72 Pollen exine: granular (0), collumellate (1), microfoveolate (2), perforate (3), smooth (4), reticulate (5)
73 Pollen operculum: absent (0), present (1)
74 Pollen vestibulum: absent (0), present (1)
75 Arrangement of female inflorescence: solitary (0), catkin/spike/racemose (1), headlike/fasciculate (2), panicle (3), cyme (4), umbel (5)
76 Pistillate flower/catkin/spike stance: pendulous (0), erect (1)
77 Female inflorescence length(mm): < 10 (0), 11–30 (1), 31–100 (2), >100 (3)
78 Female inflorescence width(mm): ≤10 (0), 11–20 (1), >20 (2)
79 Female inflorescence peduncle (mm): ≤ 10 (0), 11–30 (1), >30 (2)
80 Pistillate catkin/spike: lax (0), condensed (1)
81 Female flowers number in inflorescences: one (0), two (1), three (2), >3 (3)
82 Central flower of basic pistillate/bisexual cymule/dichasium: absent (0), reduced (1), present (2)
83 Lateral flowers of basic bisexual/female cymule/dichasium: absent (0), reduced (1), one absent (2), both present (3)
84 Pistillate flower bract number: one (0), one and half or two (1), three (2), numerous (3)
85 Pistillate flower bract length(mm): < 5 (0), 6–10 (1), >10 (2)
86 Pistillate flower bract: entire (0), 2-lobed (1), 3-lobed (2), 4-lobed (3), 5-lobed or more (4)
87 Pistillate flower bracteoles number: zero (0), two (1), three (2), four to six (3), six to eight (4), numerous (5)
88 Pistillate flower tepal number: zero/absent (0), one (1), two (2), three (3), four (4), five (5), ≥6(6)
89 Tepal whorls in pistillate number: one (0), two (1)
90 Ovary position: superior (0), inferior (1), semi-inferior (2)
91 Carpel number of (central) flower: one (0), two (1), three (2), four (3), five (4), six (5), >6 (6)
92 Carpel alignment: median (0), transverse (1), oblique (2)
93 Ovary locule number: one (0), two (1), three (2), four (3), five (4), six (5)
94 Number of ovules per ovary: one (0), two (1), three (2), four (3), six (4), twelve (5), more (6)
95 Ovules: unitegmic (0), bitegmic (1)
96 Placentation: basal (0), axile (1), parietal (2), marginal (3)

97 Ovule states: orthotropous (0), hemitropous (1), anatropous (2)
98 Style number: zero (0), one (1), two (2), three (3), four (4), five (5), six or more (6)
99 Stigmas: capitate (0), decurrent (1), punctate (2), spiralled (3), ligulate (4), linear (5)
100 Stigmas: bifurcate (0), not bifurcate (1)
101 Stigmatic area: outer-facing part of the style arms (0), inner-facing part of the style arms (1), all surfaces of the stylar arms (2)
102 Stigma orientation: carinal (0), split-carinal (1), commisural (2)
103 Endosperms: absent (0), present (1)
104 State of infructescence/fruit: erect (0), pendant (1)
105 Infructescence length(mm): ≤ 50 (0), 51–100 (1), 101–200 (2), >200 (3)
106 Fruits/involucres number of an infructescence: < 5 (0), 5–10 (1), > 10 (2)
107 Maturation: annual (0), biennial (1)
108 Fruit type: nuts (0), pods (1), capsules (2), drupe (3), berry (4), pepo (5), silique (6)
109 Leaf-like fused bract/bracteoles adnate to nutlet: unpaired (0), paired, equal (1), paired, unequal (2)
110 Leaf-like fused bract/bracteoles adnate to nutlet: open (0), closed (1)
111 Mature bracts form closed as: tubular (0), saccate (1), cupule (2), husk (3), conelike (4)
112 Fused bract/bracteoles persistent on axis: scale-like (0), sac-like (1), cone-like (2), cupule-like (3)
113 Bracts/involucre and nuts: enclosing half or less (0), enclosing more than half (1), enclosing all of nut (2)
114 Bracteoles: membranous (0) leathery (1), spiny (2), fleshy (3), spongy (4), woody (5)
115 Pistillate bract in fruit: free (0), attached (1)
116 Cupule valves stastes from the earliest stage: free (0), fused and dehisce upon maturity (1), form cups (2)
117 Cupule organization at maturity: valved (0), non-valved (1)
118 Cupule shape: subglobose (0), cup-shaped (1), irregular (2), pyramidal (3)
119 Cupule depth(mm): ≤ 10 (0), 10–20 (1), 20–50 (2), >50 (3)
120 Cupule width(mm): < 10 (0), 11–20 (1), 21–50 (2), >50 (3)
121 Appendages of cupules: spine-like (0), scale-like (1), leaf-like (2), glandular filaments (3), lamellae (4)
122 Spine on basal of cupules: solitary (0), clusters (1)
123 Cupule scales: slender (0), triangular (1)
124 Bracteoles of cupule: not united, arranged spirally (0); united, arranged in concentric rings (1); not united, arranged in concentric rings (2)
125 Number of nuts per fruit: one (0), two (1), three (2), more than three (3)
126 Fruit length (mm): ≤ 5 (0), 5–10 (1), 10–20 (2), 20–30 (3), >30 (4)
127 Fruit width (mm): ≤ 5 (0), 5–10 (1), 10–20 (2), 20–30 (3), >30 (4)
128 Fruit: wingless (0), winged (1)
129 Fruit wing orientation relative to the long axis of the nutlet: parallel (0), oblique (1), perpendicular (2)
130 Fruit wing venation: absent (0), subparallel (1), pinnate (2), basinerved (triveined/pentaveined) (3)
131 Epicarp: Leathery (0), fleshy (1)

132 Nut shape in transverse cross-section: trigonous (0), circular (1), elliptical (2), small and strongly dorsiventrally compressed (3), square/polygon (4), lenticular (5)
133 Nutshell surface: smooth (0), striate or wrinkled (1), grooved (2), ribbed (3), grooved and ribbed (4), alveolate (5), with papillae (6)
134 Nut length (mm): ≤ 5 (0), 5–10 (1), 10–20 (2), 20–30 (3), >30 (4)
135 Nut width (mm): ≤ 5 (0), 5–10 (1), 10–20 (2), 20–30 (3), >30 (4)
136 Nut mass (g/1000 seeds): ≤ 1 (0), 1–5 (1), 5–10 (2), 10–50 (3), 50–100 (4), 100–500 (5), 500–1000 (6), 1000–5000 (7), >5001 (8)
137 Wing length (mm): ≤ 5 (0), 5–10 (1), 10–20 (2), 20–30 (3), >30 (4)
138 Wing width (mm): ≤ 5 (0), 5–10 (1), 10–20 (2), 20–30 (3), >30 (4)
139 Wing shape: circular (0), petaline (1), foliaceous (2), trilobed (3), 2-winged or linear configuration (4)
140 Nutshell sclerenchyma: fibers (0), sclereids (1)
141 Nutshell lacunae: absent (0), present (1)
142 Fruit chambers at base: 1 (0), 2 (1), 3 (2), 4 (3), 8 (4)
143 Germination type: epigeal (0), hypogeal (1)
144 Alkaloids: absent (0), present (1)
145 Flavo noids: absent (0), present (1)
146 Myricetin: absent (0), present (1)
147 Phenolic acids: absent (0), present (1)
148 Phenylpropanoids: absent (0), present (1)
149 Quinones: absent (0), present (1)
150 Tannins: absent (0), present (1)
151 Terpenoids: absent (0), present (1)
152 Triterpenes of the taraxeran, ursan and lupan types: absent (0), present (1)
153 Steroids: absent (0), present (1)
154 Cinnamic acid spermidine amides in the exine of pollen grains: absent (0), present (1)
155 Biphenylheptanoids: absent (0), present (1)
156 Biphenyletherheptanoids: absent (0), present (1)

#### Text S2. Fossil sampling and constraints

The 52 fossil taxa included in this study are listed below, along with information on their age, geographic origin, and both primary and secondary references concerning the description and/or phylogenetic placement of the fossil material. We used the literature to determine the currently accepted taxonomic placement of each fossil; this information was used (along with the results from the transcriptome analyses) to generate a constraint tree for the combined molecular-mophological phylogenetic analysis (see Suppl. Fig. S1). The transcriptome tree was used as a framework to constrain major relationships, but we left many relationships in the constraint tree unresolved to allow for the combined analyses to determine more precise fossil placements. For example, several fossil taxa of *Juglans* were included in the analyses;

in the constraint tree we included all species of *Juglans* (extant and extinct) in an unresolved clade, allowing for the fossils to be placed as stem or crown members according to the phylogenetic analysis. This general approach allowed us to levereage the wealth of taxonomic and phylogenetic information from the literature to produce a better resolved phylogenetic tree, including fossils, for Fagales. Without this type of approach, the tree would be largely unresolved, due to the limited number of characters for each fossil and the frequent lack of overlap among the characters preserved, in part because the fossils represent different organ types.

| Family | Species | Age (Ma) | Locality | Reference |
|---|---|---|---|---|
| Betulaceae | *Asterocarpinus perplexans* | 33–34 | Western North America: the Florissant flora of Colorado, the lower John Day flora of Oregon, and the Grant flora of Montan | Manchester & Crane, 1987 |
| Betulaceae | *Cranea wyomingensis* | 56–66 | Northern Wyoming, USA | Manchester & Chen, 1998 |
| Betulaceae | *Palaeocarpinus dakotensis* | 56–59.2 | North Dakota, USA | Manchester *et al.*, 2004 |
| Betulaceae | *Bedellia pusilla* | 83.6–86.3 | The Allon fossil flora, Georgia, USA | Sims *et al.*, 1999 |
| Betulaceae | *Endressianthus miraensis* | 71–73 | Clay pit south of Mira, Portugal | Friis *et al.*, 2003 |
| Casuarinaceae | *Gymnostoma antiquum* | 56–59.2 | New South Wales, Australia | Scriven & Hill, 1995 |
| Casuarinaceae | *Gymnostoma argentinum* | 51–52 | NW Chubut Province, Argentina | Zamaloa *et al.*, 2006 |
| Casuarinaceae | *Gymnostoma archangelskyi* | 51–52 | NW Chubut Province, Argentina | Zamaloa *et al.*, 2006 |
| Casuarinaceae | *Gymnostoma patagonicum* | 51–52 | NW Chubut Province, Argentina | Zamaloa *et al.*, 2006 |
| Casuarinaceae | *Gymnostoma tasmanianum* | 28.1–33.9 | Little Rapid River, Tasmania | Guerin & Hill, 2003 |
| Ticodendraceae | *Ferrignocarpus bivalvis* | 47.8–56 | London Clay flora of southern England | Manchester, 2011 |
| Fagaceae | *Castanopsoidea columbiana* | 55–57 | Western Tennessee, USA | Crepet & Nixon, 1989 |
| Fagaceae | *Fagopsis longifolia* | 33–35 | Oligocene Florissant Flora of Colorado, USA | Manchester & Crane, 1983 |
| Fagaceae | *Pseudofagus idahoensis* | 14–16 | Miocene Clarkia flora of northern Idaho, USA | Smiley and Huggins, 1981 |
| Fagaceae | *Trigonobalanoidea americana* | 55–57 | Bovay locality in northern Mississippi, and the Warman locality in western Tennessee, USA | Crepet & Nixon, 1989; Blanchard *et al.*, 2016 |

| | | | | |
|---|---|---|---|---|
| Fagaceae | *Castanopsis rothwellii* | 51–53 | Laguna del Hunco flora of Chubut, southern Argentina | Wilf **et al**., 2019 |
| Fagaceae | *Protofagacea allonensis* | 72.1–83.6 | Central Georgia, U.S.A | Herendeen **et al**., 1995 |
| Fagaceae | *Eotrigonobalanus* | 81–82 | Elk Basin, Wyoming;western Greenland;Princeton Chert, British Columbia, USA | Grimsson **et al**. 2016 |
| Fagaceae | *Paraquercus campania* | 72.1–83.6 | Elk Basin, Wyoming, boundary to Montana, USA | Grimsson **et al**. 2016 |
| Fagales | *Soepadmoa cupulata* | 89.8–93.9 | Old Crossman Clay Pit, Sayreville, New Jersey, USA | Gandolfo **et al**., 2018 |
| Juglandaceae | *Beardia vancouverensis* | 41.2–47.8 | Vancouver Island, British Columbia, Canada | Elliott **et al**., 2006 |
| Juglandaceae | *Casholdia microptera* | 56–59.2 | Newbury, Berkshire, England | Crane & Manchester, 1982 |
| Juglandaceae | *Cruciptera simpsonii* | 41.2–47.8 | Washington and Wyoming, USA | Manchester, 1991 |
| Juglandaceae | *Hooleya lata* | 41.2–47.8 | Lower part of the Clarno Formation near Mitchell, Orego | Wing and Hickey,1984; Manchester, 1987 |
| Juglandaceae | *Palaeooreomunnea stoneana* | 41.2–47.8 | Middle Eocene Claiborne Formation of Kentucky and Tennessee, USA | Manchester, 1987; Wang **et al**., 2013 |
| Juglandaceae | *Paraengelhardtia eocenica* | 41.2–47.8 | Middle Eocene Claiborne Formation of Henry County, Tennessee, USA | Manchester, 1987 |
| Juglandaceae | *Palaeoplatycarya wingii* | 47.8–56 | early Eocene Wind River Formation, Wyoming, USA | Manchester, 1987 |
| Juglandaceae | *Polyptera manningii* | 59.2–61.6 | Paleocene of Wyoming and Montana, USA | Manchester & Dilcher, 1997 |
| Juglandaceae | *Alatonucula ignis* | 51–53 | Laguna del Hunco flora of Chubut, Patagonia, Argentina | Hermsen and Gandolfo, 2016; Zhang **et al**., 2021 |
| Juglandaceae | *Amurcarya lobata* | 61.6–66 | Arkhara, the Amur Province, Russian Far East | Kodrul and Krassilov, 2005; Zhang **et al**., 2021 |
| Juglandaceae | *Carya florissantensis* | 27.82–33 | Florissant beds of Colorado, USA | Manchester, 1987; Zhang **et al**., 2021 |
| Juglandaceae | *Carya quadrangula* | 24.7–30.8 | Germany | Manchester, 1987; Zhang **et al**., 2021 |
| Juglandaceae | *Cyclocarya brownii* | 56–59.2 | Rocky Mountain Region, USA | Manchester & Dilcher, 1982; Zhang **et al**., 2021 |
| Juglandaceae | *Cyclocarya tymensis* | 23–33.9 | Tym River in western Siberia, Russia | Manchester, 1987; Zhang **et al**., 2021 |
| Juglandaceae | *Juglandicarya simplicarpa* | 56–61.6 | Wyoming, USA | Manchester, 1987; Zhang **et al**., 2021 |

| | | | | |
|---|---|---|---|---|
| Juglandaceae | *Juglans clarnensis* | 41.2–47.8 | Middle Eocene; Oregon | Manchester, 1987; Zhang ***et al***., 2021 |
| Juglandaceae | *Juglans siouxensis* | 28.1–33.9 | Sioux County, Nebraska | Manchester, 1987; Zhang ***et al***., 2021 |
| Juglandaceae | *Oreomunnea grahamii* | 14–19.5 | Panama Canal | Herrera ***et al***., 2014; Zhang ***et al***., 2021; |
| Juglandaceae | *Palaeocarya clarnensis* | 41.2–47.8 | Clarno Formation of Oregon, USA | Manchester, 1987; Zhang ***et al***., 2021 |
| Juglandaceae | *Platycarya americana* | 47.8–56 | Early Eocene; North Dakota, USA | Manchester, 1987; Zhang ***et al***., 2021 |
| Juglandaceae | *Pterocarya macginitii* | 47.8–56 | Wyoming, USA | Manchester and Dilcher, 1982; Zhang ***et al***., 2021 |
| Myricaceae | *Comptonia naumannii* | 20.44–23.03 | Miocene of Weichang, China | Liang ***et al***., 2010 |
| Myricaceae | *Myricamentum eckfeldensis* | 37.7–41.2 | Rhineland-Palatinate, western Germany | Wilde ***et al***., 2021 |
| Nothofagaceae /Fagaceae | *Antiquacupula sulcata* | 83.6–86.3 | Buffalo Creek Member of the Gaillard Formation in central Georgia, USA | Sims ***et al***., 1998 |
| Normapolles | *Archaefagacea futabensis* | 86.3–89.8 | Futaba Group in northeastern Honshu, Japan | Takahashi ***et al***., 2008 |
| Normapolles | *Normanthus miraensis* | 66–83.6 | Late Cretaceous of Portugal | Scho¨ nenberger ***et al***. 2001 |
| Normapolles | *Caryanthus knoblochii* | 72.1–86.3 | Scania, Sweden | Friis ***et al***., 1983, 2006 |
| Normapolles | *Manningia crassa* | 72.1–86.3 | Scania, Sweden | Friis ***et al***., 1983 |
| Normapolles | *Budvaricarpus serialis* | 83.6–89.8 | Late Cretaceous of the Czech Republic | Heřmanová ***et al***., 2011; Zhang ***et al***., 2021 |
| Normapolles | *Calathiocarpus (C. minimus)* | 88.6–93.6 | Central Europe (Austria; the Czech Republic; Germany; the Netherlands; Poland) | Friis ***et al***., 2006 |
| Normapolles | *Dahlgrenianthus suecicus* | 72.1–83.6 | Czech Republic, Germany; the Netherlands and southern Sweden (Scania) | Friis ***et al***., 2006 |
| Normapolles | *Antiquocarya verruculosa* | 72.1–86.3 | Scania, Sweden | Friis ***et al***., 1983, 2006 |

#### 2. Supplementary Tables

**Table S1**. **Sampling information, newly sequenced transcriptomes, and previously sequenced genomes are listed here.**

Botanical garden acronyms are as follows: Arnold: The Arnold Arboretum of Harvard University; KIB: Kunming Botanical Garden, Chinese Academy of Sciences; MBG: Missouri Botanical Garden; UCBG: University of California Botanical Garden at Berkeley; XTBG: Xishuangbanna Tropical Botanical Garden, Chinese Academy of Sciences

| Order | Family | Species | Locality | Specimen No. | Molecualr data Type |
|---|---|---|---|---|---|
| Fagales | Nothofagaceae | *Nothofagus fusca* (Hook.f.) Oerst. | UCBG | Yi17112 | Transcriptome |
| Fagales | Nothofagaceae | *Nothofagus menziesii* (Hook.f.) Oerst. | UCBG | Yi17110 | Transcriptome |
| Fagales | Nothofagaceae | *Nothofagus obliqua* (Mirb.) Oerst. | UCBG | Yi17081 | Transcriptome |
| Fagales | Nothofagaceae | *Nothofagus macrocarpa* (A.DC.) F.M.Vázquez & R.A.Rodr. | UCBG | Yi17084 | Transcriptome |
| Fagales | Nothofagaceae | *Nothofagus antarctica* (G.Forst.) Oerst. | Seattle-Tacoma Area | HM2390 | Transcriptome |
| Fagales | Nothofagaceae | *Nothofagus dombeyi* (Mirb.) Oerst. | UCBG | Yi17079 | Transcriptome |
| Fagales | Fagaceae | *Fagus engleriana* Seemen ex Diels | Arnold | Yi17145 | Transcriptome |
| Fagales | Fagaceae | *Fagus japonica* Maxim. | Arnold | Yi17147 | Transcriptome |
| Fagales | Fagaceae | *Fagus grandifolia* Ehrh. | Arnold | Yi17159 | Transcriptome |
| Fagales | Fagaceae | *Fagus sylvatica* L. | UCBG | Yi17033 | Transcriptome |
| Fagales | Fagaceae | *Fagus lucida* Rehder & E.H.Wilson | Arnold | Yi17148 | Transcriptome |
| Fagales | Fagaceae | *Castanea sativa* Mill. | Arnold | Yi17135 | Transcriptome |
| Fagales | Fagaceae | *Castanea henryi* (Skan) Rehder & E.H.Wilson | KIB | Yi17379 | Transcriptome |
| Fagales | Fagaceae | *Castanea seguinii* Dode | KIB | Yi17353 | Transcriptome |
| Fagales | Fagaceae | *Castanea pumila* (L.) Mill. | Arnold | Yi17133 | Transcriptome |
| Fagales | Fagaceae | *Castanopsis orthacantha* Franch. | KIB | Yi17380 | Transcriptome |
| Fagales | Fagaceae | *Castanopsis calathiformis* (Skan) Rehder & E.H.Wilson | KIB | Yi17360 | Transcriptome |
| Fagales | Fagaceae | *Castanopsis delavayi* Franch. | KIB | Yi17381 | Transcriptome |
| Fagales | Fagaceae | *Chrysolepis chrysophylla* var. *minor* (Benth.) Munz | UCBG | Yi17115 | Transcriptome |
| Fagales | Fagaceae | *Lithocarpus mairei* (Schottky) Rehder | KIB | Yi17355 | Transcriptome |
| Fagales | Fagaceae | *Lithocarpus dealbatus* (Hook.f. & Thomson ex Miq.) Rehder | KIB | Yi17359 | Transcriptome |

| | | | | | |
|---|---|---|---|---|---|
| Fagales | Fagaceae | *Lithocarpus glaber* (Thunb.) Nakai | UCBG | Yi17021 | Transcriptome |
| Fagales | Fagaceae | *Lithocarpus echinophorus* (Hickel & A.Camus) A.Camus | KIB | Yi17354 | Transcriptome |
| Fagales | Fagaceae | *Lithocarpus henryi* (Seemen) Rehder & E.H.Wilson | KIB | HG088 | Transcriptome |
| Fagales | Fagaceae | *Lithocarpus hancei* (Benth.) Rehder | KIB | Yi17356 | Transcriptome |
| Fagales | Fagaceae | *Lithocarpus corneus* (Lour.) Rehder | XTBG | Living collection | Transcriptome |
| Fagales | Fagaceae | *Quercus glauca* Thunb. | | ZLP | Transcriptome |
| Fagales | Fagaceae | *Quercus sichourensis* (Y.C.Hsu) C.C.Huang & Y.T.Chang | KIB | Yi17358 | Transcriptome |
| Fagales | Fagaceae | *Quercus schottkyana* Rehder & E.H.Wilson | KIB | Yi17361 | Transcriptome |
| Fagales | Fagaceae | *Quercus trojana* Webb | UCBG | Yi17036 | Transcriptome |
| Fagales | Fagaceae | *Quercus cerris* L. | MBG | Yi17247 | Transcriptome |
| Fagales | Fagaceae | *Quercus libani* G.Olivier | Arnold | Yi17152 | Transcriptome |
| Fagales | Fagaceae | *Quercus suber* L. | UCBG | Yi17076 | Transcriptome |
| Fagales | Fagaceae | *Quercus castaneifolia* C.A.Mey. | Arnold | Yi17129 | Transcriptome |
| Fagales | Fagaceae | *Quercus variabilis* Blume | KIB | Yi17377 | Transcriptome |
| Fagales | Fagaceae | *Quercus acutissima* Carruth. | KIB | Yi17351 | Transcriptome |
| Fagales | Fagaceae | *Quercus franchetii* Skan | KIB | HG054 | Transcriptome |
| Fagales | Fagaceae | *Quercus ilex* L. | MBG | Yi17243 | Transcriptome |
| Fagales | Fagaceae | *Quercus nigra* | MBG | Yi17235 | Transcriptome |
| Fagales | Fagaceae | *Quercus laurina* Bonpl. | UCBG | Yi17074 | Transcriptome |
| Fagales | Fagaceae | *Quercus palustris* Münchh. | Arnold | Yi17158 | Transcriptome |
| Fagales | Fagaceae | *Quercus shumardii* Buckley | Arnold | Yi17161 | Transcriptome |
| Fagales | Fagaceae | *Quercus acerifolia* (E.J.Palmer) Stoynoff & Hess | Arnold | Yi17131 | Transcriptome |
| Fagales | Fagaceae | *Quercus phellos* L. | MBG | Yi17246 | Transcriptome |
| Fagales | Fagaceae | *Quercus benthamii* A.DC. | UCBG | Yi17073 | Transcriptome |
| Fagales | Fagaceae | *Quercus imbricaria* Michx. | Arnold | Yi17146 | Transcriptome |
| Fagales | Fagaceae | *Quercus ellipsoidalis* E.J.Hill | Arnold | Yi17128 | Transcriptome |
| Fagales | Fagaceae | *Quercus falcata* Michx. | MBG | Yi17234 | Transcriptome |
| Fagales | Fagaceae | *Quercus velutina* Lam. | Arnold | Yi17126 | Transcriptome |
| Fagales | Fagaceae | *Quercus marilandica* (L.) Münchh. | Arnold | Yi17155 | Transcriptome |
| Fagales | Fagaceae | *Quercus dumosa* Nutt. | UCBG | Yi17057 | Transcriptome |
| Fagales | Fagaceae | *Quercus stellata* Wangenh. | Arnold | Yi17132 | Transcriptome |
| Fagales | Fagaceae | *Quercus montana* Willd. | UCBG | Yi17055 | Transcriptome |
| Fagales | Fagaceae | *Quercus greggii (A.DC.)* Trel. | UCBG | Yi17071 | Transcriptome |
| Fagales | Fagaceae | *Quercus* sect. *Lobatae* | KIB | Yi17352 | Transcriptome |
| Fagales | Fagaceae | *Quercus gambelii* Nutt. | Arnold | Yi17140 | Transcriptome |

| | | | | | |
|---|---|---|---|---|---|
| Fagales | Fagaceae | *Quercus garryana* Douglas ex Hook. | UCBG | Yi17016 | Transcriptome |
| Fagales | Fagaceae | *Quercus macrocarpa* Michx. | MBG | Yi17228 | Transcriptome |
| Fagales | Fagaceae | *Quercus muehlenbergii* Engelm. | MBG | Yi17231 | Transcriptome |
| Fagales | Fagaceae | *Quercus prinoides* | Arnold | Yi17134 | Transcriptome |
| Fagales | Fagaceae | *Quercus alba* L. | Arnold | Yi17162 | Transcriptome |
| Fagales | Fagaceae | *Quercus arkansana* Sarg. | Arnold | Yi17130 | Transcriptome |
| Fagales | Fagaceae | *Quercus bicolor* Willd. | MBG | Yi17229 | Transcriptome |
| Fagales | Fagaceae | *Quercus dentata* Thunb. | MBG | Yi17233 | Transcriptome |
| Fagales | Fagaceae | *Quercus petraea* (Matt.) Liebl. | Arnold | Yi17143 | Transcriptome |
| Fagales | Fagaceae | *Quercus lyrata* Walter | Arnold | Yi17151 | Transcriptome |
| Fagales | Fagaceae | *Quercus frainetto* Ten. | Arnold | Yi17141 | Transcriptome |
| Fagales | Fagaceae | *Quercus serrata* Murray | KIB | Yi17357 | Transcriptome |
| Fagales | Fagaceae | *Quercus pontica* K.Koch | MBG | Yi17232 | Transcriptome |
| Fagales | Fagaceae | *Quercus infectoria* G.Olivier | MBG | Yi17237 | Transcriptome |
| Fagales | Fagaceae | *Quercus michauxii* Nutt. | Arnold | Yi17156 | Transcriptome |
| Fagales | Fagaceae | *Quercus boyntonii* Beadle | MBG | Yi17239 | Transcriptome |
| Fagales | Fagaceae | *Quercus prinus* L.(synonym of *Quercus michauxii* Nutt.) | Arnold | Yi17150 | Transcriptome |
| Fagales | Fagaceae | *Quercus aliena* Blume | Arnold | Yi17160 | Transcriptome |
| Fagales | Fagaceae | *Quercus chrysolepis* Liebm. | UCBG | Yi17044 | Transcriptome |
| Fagales | Fagaceae | *Quercus cedrosensis* C.H.Mull. | UCBG | Yi17018 | Transcriptome |
| Fagales | Fagaceae | *Quercus palmeri* Engelm. | UCBG | Yi17019 | Transcriptome |
| Fagales | Fagaceae | *Quercus virginiana* Mill. | UCBG | Yi17056 | Transcriptome |
| Fagales | Fagaceae | *Trigonobalanus doichangensis* (A.Camus) Forman | KIB | Yi17376 | Transcriptome |
| Fagales | Fagaceae | *Notholithocarpus densiflorus* (Hook. & Arn.) Manos, Cannon & S.H.Oh | UCBG | Yi17042 | Transcriptome |
| Fagales | Myricaceae | *Comptonia peregrine* (L.) Coult. | | HM2020 | Transcriptome |
| Fagales | Myricaceae | *Myrica gale* L. | Arnold | Yi17157 | Transcriptome |
| Fagales | Myricaceae | *Morella rubra* Lour. | KIB | Yi17368 | Transcriptome |
| Fagales | Myricaceae | *Morella nagi* | UCBG | Yi17030 | Transcriptome |
| Fagales | Myricaceae | *Morella nana* (A. Chev.) J.Herb. | KIB | Yi17367 | Transcriptome |
| Fagales | Myricaceae | *Morella faya* (Aiton) Wilbur | UCBG | Yi17034 | Transcriptome |
| Fagales | Myricaceae | *Morella californica* (Cham.) Wilbur | UCBG | Yi17058 | Transcriptome |
| Fagales | Myricaceae | *Morella californica* (Cham.) Wilbur | UCBG | Yi17114 | Transcriptome |
| Fagales | Myricaceae | *Morella cerifera* (L.) Small | Costa Rica | NZ9348 | Transcriptome |
| Fagales | Juglandaceae | *Rhoiptelea chiliantha* Diels & Hand.-Mazz. | KIB | Yi17382 | Transcriptome |

| | | | | | |
|---|---|---|---|---|---|
| Fagales | Juglandaceae | *Platycarya strobilacea* Siebold & Zucc. | KIB | Yi17371 | Transcriptome |
| Fagales | Juglandaceae | *Carya illinoinensis* (Wangenh.) K.Koch | KIB | Yi17370 | Transcriptome |
| Fagales | Juglandaceae | *Carya ovata* var. *mexicana* (Engelm. ex Hemsl.) W.E.Manning | UCBG | Yi17070 | Transcriptome |
| Fagales | Juglandaceae | *Carya glabra* (Mill.) Sweet | Arnold | Yi17123 | Transcriptome |
| Fagales | Juglandaceae | *Carya tomentosa* (synonym of *Carya alba* (L.) Nutt. ex Elliott) | Arnold | Yi17125 | Transcriptome |
| Fagales | Juglandaceae | *Carya palmeri* W.E.Manning | UCBG | Yi17072 | Transcriptome |
| Fagales | Juglandaceae | *Juglans regia* L. | NCBI | SRR1767236 | Transcriptome |
| Fagales | Juglandaceae | *Juglans nigra* L. | NCBI | NCBI: txid16719 | Transcriptome |
| Fagales | Juglandaceae | *Juglans olanchana* Standl. & L.O.Williams | UCBG | Yi17069 | Transcriptome |
| Fagales | Juglandaceae | *Juglans mandshurica* Maxim. | KIB | Yi17383 | Transcriptome |
| Fagales | Juglandaceae | *Juglans cinerea* L. | Arnold | Yi17127 | Transcriptome |
| Fagales | Juglandaceae | *Juglans hirsuta* W.E. Manning | UCBG | Yi17068 | Transcriptome |
| Fagales | Juglandaceae | *Pterocarya macroptera* var. *delavayi* (Franch.) W.E. Manning | KIB | Yi17384 | Transcriptome |
| Fagales | Juglandaceae | *Pterocarya stenoptera* C. DC. | UCBG | Yi17039 | Transcriptome |
| Fagales | Juglandaceae | *Pterocarya tonkinensis* (Franch.) Dode | KIB | Yi17372 | Transcriptome |
| Fagales | Juglandaceae | *Pterocarya pterocarpa* Kunth ex I. Iljinsk. | KIB | Yi17373 | Transcriptome |
| Fagales | Juglandaceae | *Cyclocarya paliurus* (Batalin) Iljinsk. | KIB | Yi17374 | Transcriptome |
| Fagales | Juglandaceae | *Engelhardia spicata* | KIB | Yi17385 | Transcriptome |
| Fagales | Juglandaceae | *Engelhardia roxburghiana* Wall. | Leshan, Sichuan | Yi17375 | Transcriptome |
| Fagales | Juglandaceae | *Oreomunnea pterocarpa* Oerst. | Costa Rica | NZ8894 | Transcriptome |
| Fagales | Juglandaceae | *Alfaroa guanacastensis* D.E.Stone | Costa Rica | NZ9187 | Transcriptome |
| Fagales | Casuarinaceae | *Allocasuarina muelleriana* (Miq.) L.A.S.Johnson | UCBG | Yi17111 | Transcriptome |
| Fagales | Casuarinaceae | *Allocasuarina torulosa* (Aiton) L.A.S.Johnson | UCBG | Yi17100 | Transcriptome |
| Fagales | Casuarinaceae | *Allocasuarina decaisneana* (F.Muell.) L.A.S.Johnson | Brisbane Botanic Garden | Yi14632 | Transcriptome |
| Fagales | Casuarinaceae | *Casuarina cunninghamiana* Miq. | | ZC68 | Transcriptome |

| | | | | | |
|---|---|---|---|---|---|
| Fagales | Casuarinaceae | *Casuarina junghuhniana* Miq. | NCBI | SRR2239112 | Transcriptome |
| Fagales | Casuarinaceae | *Casuarina equisetifolia* L. | KIB | Yi17369 | Transcriptome |
| Fagales | Casuarinaceae | *Gymnostoma australianum* L.A.S.Johnson | UCBG | Yi17108 | Transcriptome |
| Fagales | Betulaceae | *Alnus firma* Siebold & Zucc. | Arnold | Yi17153 | Transcriptome |
| Fagales | Betulaceae | *Alnus viridis var.sinuata* (sn of *Alnus alnobetula* subsp. *sinuata* (Regel) Raus.) | UCBG | Yi17015 | Transcriptome |
| Fagales | Betulaceae | *Alnus japonica* (Thunb.) Steud. | KIB Germplasm Bank of Wild Species | lilan531 | Transcriptome |
| Fagales | Betulaceae | *Alnus acuminata* Kunth | UCBG | Yi17067 | Transcriptome |
| Fagales | Betulaceae | *Alnus glutinosa* (L.) Gaertn. | UCBG | Yi17031 | Transcriptome |
| Fagales | Betulaceae | *Alnus nepalensis* D.Don | KIB | Yi17386 | Transcriptome |
| Fagales | Betulaceae | *Alnus ferdinandi-coburgii* C.K.Schneid. | KIB | Yi17378 | Transcriptome |
| Fagales | Betulaceae | *Alnus cremastogyne* Burkill | KIB Germplasm Bank of Wild Species | TanCM1351 | Transcriptome |
| Fagales | Betulaceae | *Alnus trabeculosa* Hand.-Mazz. | KIB Germplasm Bank of Wild Species | TanCM421 | Transcriptome |
| Fagales | Betulaceae | *Betula luminifera* H.J.P.Winkl. | KIB | Yi17363 | Transcriptome |
| Fagales | Betulaceae | *Betula maximowicziana* Regel | Arnold | Yi17120 | Transcriptome |
| Fagales | Betulaceae | *Betula schmidtii* Regel | UCBG | Yi17026 | Transcriptome |
| Fagales | Betulaceae | *Betula chinensis* Maxim. | Arnold | Yi17121 | Transcriptome |
| Fagales | Betulaceae | *Betula lenta* L. | Arnold | Yi17119 | Transcriptome |
| Fagales | Betulaceae | *Betula ovalifolia* Rupr. | KIB Germplasm Bank of Wild Species | LiuB0006 | Transcriptome |
| Fagales | Betulaceae | *Betula pumila* L. | UCBG | Yi17050 | Transcriptome |
| Fagales | Betulaceae | *Betula fruticosa* Pall. | Arnold | Yi17124 | Transcriptome |
| Fagales | Betulaceae | *Betula pubescens* Ehrh. | Arnold | Yi17118 | Transcriptome |
| Fagales | Betulaceae | *Betula costata* Trautv. | KIB Germplas | NiuYL473 | Transcriptome |

| | | | | | |
|---|---|---|---|---|---|
| | | | m Bank of Wild Species | | |
| Fagales | Betulaceae | *Betula utilis* D. Don | KIB Germplas m Bank of Wild Species | 09CS1574 | Transcriptome |
| Fagales | Betulaceae | *Betula albosinensis* Burkill | KIB Germplas m Bank of Wild Species | SCU-11-123 | Transcriptome |
| Fagales | Betulaceae | *Betula dahurica* Pall. | Arnold | Yi17122 | Transcriptome |
| Fagales | Betulaceae | *Betula corylifolia* Regel & Maxim. | Royal Botanic Garden Edinburgh | Living collection | Transcriptome |
| Fagales | Betulaceae | *Carpinus cordata* Blume | Arnold | Yi17139 | Transcriptome |
| Fagales | Betulaceae | *Carpinus japonica* Blume | Arnold | Yi17138 | Transcriptome |
| Fagales | Betulaceae | *Carpinus betulus* L. | Arnold | Yi17136 | Transcriptome |
| Fagales | Betulaceae | *Carpinus londoniana* H.J.P.Winkl. | KIB Germplas m Bank of Wild Species | YNS0231 | Transcriptome |
| Fagales | Betulaceae | *Carpinus monbeigiana* Hand.-Mazz. | KIB | Yi17387 | Transcriptome |
| Fagales | Betulaceae | *Carpinus orientalis* Mill. | Arnold | Yi17137 | Transcriptome |
| Fagales | Betulaceae | *Carpinus putoensis* W.C.Cheng | KIB | Yi17388 | Transcriptome |
| Fagales | Betulaceae | *Carpinus turzaninowii* Hance | KIB | Yi17365 | Transcriptome |
| Fagales | Betulaceae | *Corylus avellana* L. | NCBI | SRR527087 | Transcriptome |
| Fagales | Betulaceae | *Corylus americana* Walter | UCBG | Yi17053 | Transcriptome |
| Fagales | Betulaceae | *Corylus yunnanensis* (Franch.) A.Camus | KIB | Yi17366 | Transcriptome |
| Fagales | Betulaceae | *Corylus chinensis* Franch. | KIB | Yi17362 | Transcriptome |
| Fagales | Betulaceae | *Corylus cornuta* Marshall | UCBG | Yi17051 | Transcriptome |
| Fagales | Betulaceae | *Corylus sieboldiana* Blume | | ZLP | Transcriptome |
| Fagales | Betulaceae | *Corylus ferox* var. *tibetica* (Batalin) Franch. (*C. tibetica* ) | UCBG | Yi17025 | Transcriptome |
| Fagales | Betulaceae | *Ostrya japonica* Sarg. | KIB | Yi17364 | Transcriptome |
| Fagales | Betulaceae | *Ostrya virginiana* (Mill.) K.Koch | UCBG | Yi17040 | Transcriptome |
| Fagales | Betulaceae | *Ostryopsis davidiana* Decne. | Arnold | Yi17163 | Transcriptome |

| | | | | | |
|---|---|---|---|---|---|
| Fagales | Betulaceae | *Ostryopsis nobilis* Balf.f. & W.W.Sm. | UCBG | Yi17028 | Transcriptome |
| Fagales | Ticodendraceae | *Ticodendron incognitum* Gómez-Laur. & L.D.Gómez | Costa Rica | Yi17389 | Transcriptome |
| Ranunculales | Eupteleaceae | *Euptelea pleiosperma* Hook.f. & Thomson | NCBI | ERR2040156 | Transcriptome |
| Ranunculales | Papaveraceae | *Papaver somniferum* L. | NCBI | SRR6378470 | Transcriptome |
| Dilleniales | Dilleniaceae | *Hibbertia grossulariifolia* Salisb. | NCBI | ERR2040191 | Transcriptome |
| Apiales | Araliaceae | *Hydrocotyle umbellata* L. | NCBI | ERR2040643 | Transcriptome |
| Apiales | Araliaceae | *Polyscias fruticosa* (L.) Harms | NCBI | ERR2040642 | Transcriptome |
| Saxifragales | Altingiaceae | *Liquidambar formosana* Hance | NCBI | SRR1514949 | Transcriptome |
| Saxifragales | Hamamelidaceae | *Distylium buxifolium* (Hance) Merr. | NCBI | SRR6374699 | Transcriptome |
| Brassicales | Brassicaceae | *Arabidopsis thaliana* (L.) Heynh. | PhytozomeV13 | TAIR10 | Genome |
| Malvales | Malvaceae | *Gossypium hirsutum* L. | NCBI | ASM98774v1 | Transcriptome |
| Fabales | Quillajaceae | *Quillaja saponaria* Molina | NCBI | GCA_003338715.1 | Genome |
| Fabales | Fabaceae | *Arachis hypogaea* L. | NCBI | SRR6387670 | Transcriptome |
| Fabales | Fabaceae | *Bauhinia blakeana* Dunn | NCBI | ERR2021955 | Transcriptome |
| Fabales | Fabaceae | *Cercis gigantea* | NCBI | SRR957672 | Transcriptome |
| Fabales | Fabaceae | *Glycine max* (L.) Merr. | PhytozomeV13 | Wm82.a2.v1 | Genome |
| Cucurbitales | Datiscaceae | *Datisca glomerata* (C.Presl) Baill. | GigaDB | ftp://parrot.genomics.cn/gigadb/pub/10.5524/101001_102000/101046 | Genome |
| Cucurbitales | Begoniaceae | *Begonia fuchsioides* Hook. | GigaDB | ftp://parrot.genomics.cn/gigadb/pub/10.5524/101001_102000/101043/ | Genome |

| | | | | | |
|---|---|---|---|---|---|
| Cucurbitales | Cucurbitaceae | *Trichosanthes kirilowii* Maxim. | NCBI | SRR5259241 | Transcriptome |
| Cucurbitales | Cucurbitaceae | *Gynostemma pentaphyllum* (Thunb.) Makino | NCBI | SRR3886577 | Transcriptome |
| Cucurbitales | Cucurbitaceae | *Cucumis sativus* L. | PhytozomeV13 | v1.0 | Genome |
| Rosales | Rosaceae | *Prunus persica* (L.) Batsch | PhytozomeV13 | v2.1 | Genome |
| Rosales | Rhamnaceae | *Ochetophila trinervis* (Gillies ex Hook.) Poepp. ex Endl. (*Discaria trinervis*) | GigaDB | ftp://parrot.genomics.cn/gigadb/pub/10.5524/101001_102000/101048/ | Genome |
| Rosales | Rhamnaceae | *Rhamnus japonica* Maxim. | NCBI | ERR2040416 | Transcriptome |
| Rosales | Rhamnaceae | *Ziziphus jujuba* Mill. | NCBI | SRR3348541 | Transcriptome |
| Rosales | Elaeagnaceae | *Hippophae rhamnoides* subsp. *mongolica* | NCBI | ERR1294014 | Transcriptome |

**Table S2. Rooting strategy of gene trees**

| Root hierarchical strategies | Species |
|---|---|
| 1 | *Euptelea pleiosperma, Papaver somniferum, Hibbertia grossulariifolia* |
| 2 | *Hydrocotyle umbellate, Polyscias fruticosa* |
| 3 | *Liquidambar formosana, Distylium buxifolium* |
| 4 | *Arabidopsis thaliana, Gossypium hirsutum* |
| 5 | *Prunus persica, Hippophae rhamnoides* subsp. *Mongolica, Rhamnus japonica, Ziziphus jujuba, Discaria trinervis* |
| 6 | *Datisca glomerata, Begonia fuchsioides, Gynostemma pentaphyllum, Cucumis sativus, Trichosanthes kirilowii* |
| 7 | *Quillaja saponaria, Cercis gigantea, Bauhinia blakeana, Glycine max, Arachis hypogaea* |

**Table S3. Divergence time of major lineages**

| Lineages | Stem (Ma) | Crown (Ma) |
|---|---|---|
| **Fagales_fossil** | **108.47** | **105.04** |
| **Fagales** | **105.04** | **103.44** |
| **Nothofaceae_fossil_*Nothofagus*** | **103.44** | **99.31** |
| **Nothofaceae_*Nothofagus*** | **99.31** | **66.99** |
| **Fagaceae_fossil** | **102.76** | **98.8** |
| **Fagaceae** | **94.3** | **89.8** |
| Quercoids | 85.4 | 82.63 |
| Quercoids I | 82.63 | 81.23 |
| Quercoids II | 82.63 | 74.15 |
| Fagaceae_*Quercus* | 62.2 | 56 |
| Fagaceae_*Notholithocarpus* | 62.2 | 62.2 |
| Fagaceae_*Chrysolepis* | 66.4 | 66.4 |
| Fagaceae_*Lithocarpus* | 74.15 | 73.23 |
| Fagaceae_*Castanea* | 65.32 | 58.84 |
| Fagaceae_*Castanopsis* | 65.32 | 43.44 |
| Fagaceae_*Fagus* | 53.89 | 40.73 |
| Fagaceae_*Trigonobalanus* | 85.4 | 71.9 |
| **Juglandaceae_fossil** | **96.76** | **94** |
| **Juglandaceae** | **93.3** | **92.41** |
| Juglandaceae_*Alfaroa* | 35.23 | 35.23 |
| Juglandaceae_*Oreomunnea* | 35.23 | 27.1 |
| Juglandaceae_*Engelhardia* | 41.7 | 41.7 |
| Juglandaceae_*Alfaropsis* | 39.7 | 39.7 |
| Juglandaceae_*Juglans* | 63.99 | 63.99 |
| Juglandaceae_*Cyclocarya* | 60.83 | 60.36 |
| Juglandaceae_*Platycarya* | 60.92 | 60.16 |
| Juglandaceae_*Carya* | 70.63 | 67.03 |
| Juglandaceae_*Pterocarya* | 63.99 | 63.97 |
| Juglandaceae_*Rhoiptelea* | 92.41 | 92.41 |
| **Myricaceae_fossil** | **96.76** | **79.55** |
| **Myricaceae** | **79.55** | **76.4** |
| Myricaceae_*Morella* | 57.78 | 36.62 |
| Myricaceae_*Myrica* | 57.78 | 57.78 |
| Myricaceae_*Comptonia* | 75.56 | 75.56 |
| Myricaceae_*Canacomyrica* | - | - |
| **Casuarinaceae** | **97.54** | **81.06** |
| Casuarinaceae_*Allocasuarina* | 65.87 | 58.78 |
| Casuarinaceae_*Casuarina* | 65.87 | 43.6 |
| Casuarinaceae_*Ceuthostoma* | - | - |
| Casuarinaceae_*Gymnostoma* | 81.06 | 75.31 |
| **Ticodendraceae_Ticodendron** | **95.02** | **73.67** |

| | | |
|---|---|---|
| **Betulaceae_fossil** | **95.02** | **93.41** |
| **Betulaceae** | **93.41** | **78.18** |
| Betulaceae_*Alnus* | 60.07 | 36.01 |
| Betulaceae_*Betula* | 60.07 | 47.08 |
| Betulaceae_*Carpinus* | 24.02 | 17.63 |
| Betulaceae_*Ostrya* | 24.02 | 21.29 |
| Betulaceae_*Ostryopsis* | 53.84 | 22.83 |
| Betulaceae_*Corylus* | 66.37 | 63.3 |

### 3. Supplementary Figures

**Fig. S1 Topological constraint used for combined phylogenetic analyses.** Major relationships based on the transcriptome analyses, with fossil placements based on the literature.

83

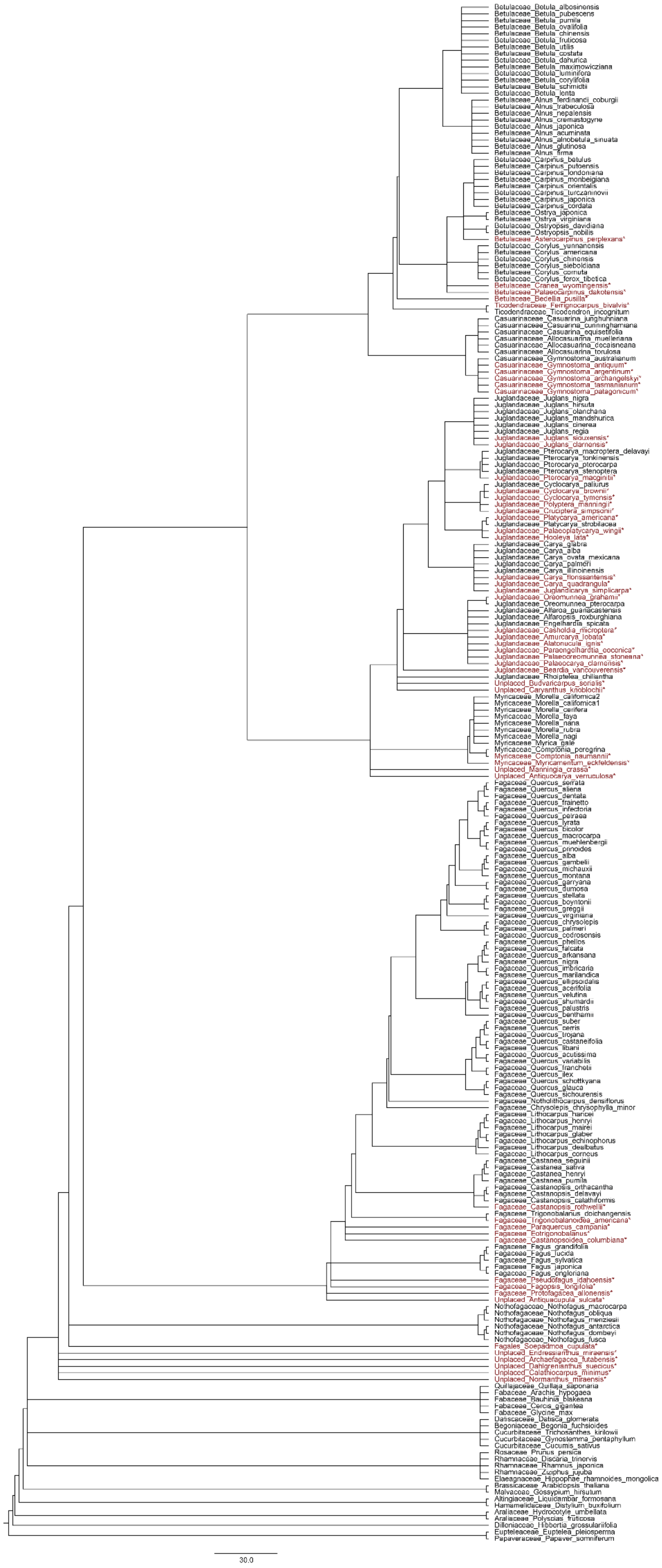

Betulaceae_Betula_albosinensis
Betulaceae_Betula_pubescens
Betulaceae_Betula_pumila
Betulaceae_Betula_ovalifolia
Betulaceae_Betula_chinensis
Betulaceae_Betula_fruticosa
Betulaceae_Betula_utilis
Betulaceae_Betula_costata
Betulaceae_Betula_dahurica
Betulaceae_Betula_maximowicziana
Betulaceae_Betula_luminifera
Betulaceae_Betula_corylifolia
Betulaceae_Betula_schmidtii
Betulaceae_Betula_lenta
Betulaceae_Alnus_ferdinandi_coburgii
Betulaceae_Alnus_trabeculosa
Betulaceae_Alnus_nepalensis
Betulaceae_Alnus_cremastogyne
Betulaceae_Alnus_japonica
Betulaceae_Alnus_acuminata
Betulaceae_Alnus_alnobetula_sinuata
Betulaceae_Alnus_glutinosa
Betulaceae_Alnus_firma
Betulaceae_Carpinus_betulus
Betulaceae_Carpinus_putoensis
Betulaceae_Carpinus_londoniana
Betulaceae_Carpinus_monbeigiana
Betulaceae_Carpinus_orientalis
Betulaceae_Carpinus_turczaninovii
Betulaceae_Carpinus_japonica
Betulaceae_Carpinus_cordata
Betulaceae_Ostrya_japonica
Betulaceae_Ostrya_virginiana
Betulaceae_Ostryopsis_davidiana
Betulaceae_Ostryopsis_nobilis
Betulaceae_Asterocarpinus_perplexans*
Betulaceae_Corylus_yunnanensis
Betulaceae_Corylus_americana
Betulaceae_Corylus_chinensis
Betulaceae_Corylus_sieboldiana
Betulaceae_Corylus_cornuta
Betulaceae_Corylus_ferox_tibetica
Betulaceae_Cranea_wyomingensis*
Betulaceae_Palaeocarpinus_dakotensis*
Betulaceae_Bedellia_pusilla*
Ticodendraceae_Ferrignocarpus_bivalvis*
Ticodendraceae_Ticodendron_incognitum
Casuarinaceae_Casuarina_junghuhniana
Casuarinaceae_Casuarina_cunninghamiana
Casuarinaceae_Casuarina_equisetifolia
Casuarinaceae_Allocasuarina_muelleriana
Casuarinaceae_Allocasuarina_decaisneana
Casuarinaceae_Allocasuarina_torulosa
Casuarinaceae_Gymnostoma_australianum
Casuarinaceae_Gymnostoma_antiquum*
Casuarinaceae_Gymnostoma_argentinum*
Casuarinaceae_Gymnostoma_archangelskyi*
Casuarinaceae_Gymnostoma_tasmanianum*
Casuarinaceae_Gymnostoma_patagonicum*
Juglandaceae_Juglans_nigra
Juglandaceae_Juglans_hirsuta
Juglandaceae_Juglans_olanchana
Juglandaceae_Juglans_mandshurica
Juglandaceae_Juglans_cinerea
Juglandaceae_Juglans_regia
Juglandaceae_Juglans_siouxensis*
Juglandaceae_Juglans_clarnensis*
Juglandaceae_Pterocarya_macroptera_delavayi
Juglandaceae_Pterocarya_tonkinensis
Juglandaceae_Pterocarya_pterocarpa
Juglandaceae_Pterocarya_stenoptera
Juglandaceae_Pterocarya_macginitii*
Juglandaceae_Cyclocarya_paliurus
Juglandaceae_Cyclocarya_brownii*
Juglandaceae_Cyclocarya_tymensis*
Juglandaceae_Polyptera_manningii*
Juglandaceae_Cruciptera_simpsonii*
Juglandaceae_Platycarya_americana*
Juglandaceae_Platycarya_strobilacea
Juglandaceae_Palaeoplatycarya_wingii*
Juglandaceae_Hooleya_lata*
Juglandaceae_Carya_glabra
Juglandaceae_Carya_alba
Juglandaceae_Carya_ovata_mexicana
Juglandaceae_Carya_palmeri
Juglandaceae_Carya_illinoinensis
Juglandaceae_Carya_florissantensis*
Juglandaceae_Carya_quadrangula*
Juglandaceae_Juglandicarya_simplicarpa*
Juglandaceae_Oreomunnea_grahamii*
Juglandaceae_Oreomunnea_pterocarpa
Juglandaceae_Alfaroa_guanacastensis
Juglandaceae_Alfaropsis_roxburghiana
Juglandaceae_Engelhardia_spicata
Juglandaceae_Casholdia_microptera*
Juglandaceae_Amurcarya_lobata*
Juglandaceae_Alatonucula_ignis*
Juglandaceae_Paraengelhardtia_eocenica*
Juglandaceae_Palaeooreomunnea_stoneana*
Juglandaceae_Palaeocarya_clarnensis*
Juglandaceae_Beardia_vancouverensis*
Juglandaceae_Rhoiptelea_chiliantha
Unplaced_Budvaricarpus_serialis*
Unplaced_Caryanthus_knoblochii*
Myricaceae_Morella_californica2
Myricaceae_Morella_californica1
Myricaceae_Morella_cerifera
Myricaceae_Morella_faya
Myricaceae_Morella_nana
Myricaceae_Morella_rubra
Myricaceae_Morella_nagi
Myricaceae_Myrica_gale
Myricaceae_Comptonia_peregrina
Myricaceae_Comptonia_naumannii*
Myricaceae_Myricamentum_eckfeldensis*
Unplaced_Manningia_crassa*
Unplaced_Antiquocarya_verruculosa*
Fagaceae_Quercus_serrata
Fagaceae_Quercus_aliena
Fagaceae_Quercus_dentata
Fagaceae_Quercus_frainetto
Fagaceae_Quercus_infectoria
Fagaceae_Quercus_petraea
Fagaceae_Quercus_lyrata
Fagaceae_Quercus_bicolor
Fagaceae_Quercus_macrocarpa
Fagaceae_Quercus_muehlenbergii
Fagaceae_Quercus_prinoides
Fagaceae_Quercus_alba
Fagaceae_Quercus_gambelii
Fagaceae_Quercus_michauxii
Fagaceae_Quercus_montana
Fagaceae_Quercus_garryana
Fagaceae_Quercus_dumosa
Fagaceae_Quercus_stellata
Fagaceae_Quercus_boyntonii
Fagaceae_Quercus_greggii
Fagaceae_Quercus_virginiana
Fagaceae_Quercus_chrysolepis
Fagaceae_Quercus_palmeri
Fagaceae_Quercus_cedrosensis
Fagaceae_Quercus_phellos
Fagaceae_Quercus_falcata
Fagaceae_Quercus_arkansana
Fagaceae_Quercus_nigra
Fagaceae_Quercus_imbricaria
Fagaceae_Quercus_marilandica
Fagaceae_Quercus_ellipsoidalis
Fagaceae_Quercus_acerifolia
Fagaceae_Quercus_velutina
Fagaceae_Quercus_shumardii
Fagaceae_Quercus_palustris
Fagaceae_Quercus_benthamii
Fagaceae_Quercus_suber
Fagaceae_Quercus_cerris
Fagaceae_Quercus_trojana
Fagaceae_Quercus_castaneifolia
Fagaceae_Quercus_libani
Fagaceae_Quercus_acutissima
Fagaceae_Quercus_variabilis
Fagaceae_Quercus_franchetii
Fagaceae_Quercus_ilex
Fagaceae_Quercus_schottkyana
Fagaceae_Quercus_glauca
Fagaceae_Quercus_sichourensis
Fagaceae_Notholithocarpus_densiflorus
Fagaceae_Chrysolepis_chrysophylla_minor
Fagaceae_Lithocarpus_hancei
Fagaceae_Lithocarpus_henryi
Fagaceae_Lithocarpus_mairei
Fagaceae_Lithocarpus_glaber
Fagaceae_Lithocarpus_echinophorus
Fagaceae_Lithocarpus_dealbatus
Fagaceae_Lithocarpus_corneus
Fagaceae_Castanea_seguinii
Fagaceae_Castanea_sativa
Fagaceae_Castanea_henryi
Fagaceae_Castanea_pumila
Fagaceae_Castanopsis_orthacantha
Fagaceae_Castanopsis_delavayi
Fagaceae_Castanopsis_calathiformis
Fagaceae_Castanopsis_rothwellii*
Fagaceae_Trigonobalanus_doichangensis
Fagaceae_Trigonobalanoidea_americana*
Fagaceae_Paraquercus_campania*
Fagaceae_Eotrigonobalanus*
Fagaceae_Castanopsoidea_columbiana*
Fagaceae_Fagus_grandifolia
Fagaceae_Fagus_lucida
Fagaceae_Fagus_sylvatica
Fagaceae_Fagus_japonica
Fagaceae_Fagus_engleriana
Fagaceae_Pseudofagus_idahoensis*
Fagaceae_Fagopsis_longifolia*
Fagaceae_Protofagacea_allonensis*
Unplaced_Antiquacupula_sulcata*
Nothofagaceae_Nothofagus_macrocarpa
Nothofagaceae_Nothofagus_obliqua
Nothofagaceae_Nothofagus_menziesii
Nothofagaceae_Nothofagus_antarctica
Nothofagaceae_Nothofagus_dombeyi
Nothofagaceae_Nothofagus_fusca
Fagales_Soepadmoa_cupulata*
Unplaced_Endressianthus_miraensis*
Unplaced_Archaefagacea_futabensis*
Unplaced_Dahlgrenianthus_suecicus*
Unplaced_Calathiocarpus_minimus*
Unplaced_Normanthus_miraensis*
Quillajaceae_Quillaja_saponaria
Fabaceae_Arachis_hypogaea
Fabaceae_Bauhinia_blakeana
Fabaceae_Cercis_gigantea
Fabaceae_Glycine_max
Datiscaceae_Datisca_glomerata
Begoniaceae_Begonia_fuchsioides
Cucurbitaceae_Trichosanthes_kirilowii
Cucurbitaceae_Gynostemma_pentaphyllum
Cucurbitaceae_Cucumis_sativus
Rosaceae_Prunus_persica
Rhamnaceae_Discaria_trinervis
Rhamnaceae_Rhamnus_japonica
Rhamnaceae_Ziziphus_jujuba
Elaeagnaceae_Hippophae_rhamnoides_mongolica
Brassicaceae_Arabidopsis_thaliana
Malvaceae_Gossypium_hirsutum
Altingiaceae_Liquidambar_formosana
Hamamelidaceae_Distylium_buxifolium
Araliaceae_Hydrocotyle_umbellata
Araliaceae_Polyscias_fruticosa
Dilleniaceae_Hibbertia_grossulariifolia
Eupteleaceae_Euptelea_pleiosperma
Papaveraceae_Papaver_somniferum
30.0

**Fig. S2 Topological constraint used for supertree reconstruction.** We constrained relationships among all the genera (and sections within *Quercus*) based on the relationships inferred from the transcriptomic data.

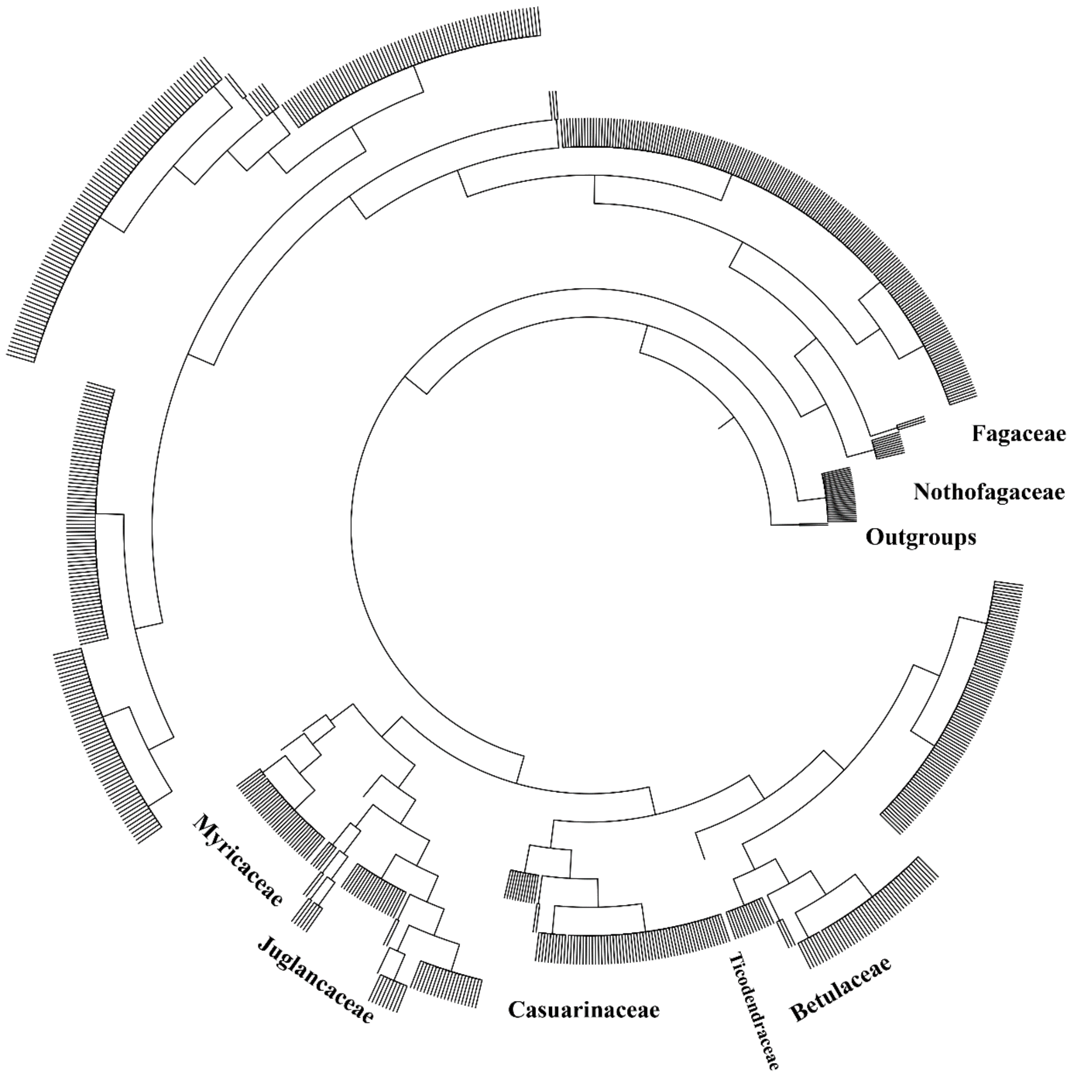

**Fig. S3 The maximum-likelihood tree of Fagales inferred from 643 orthologous genes by RAxML (CML tree).**

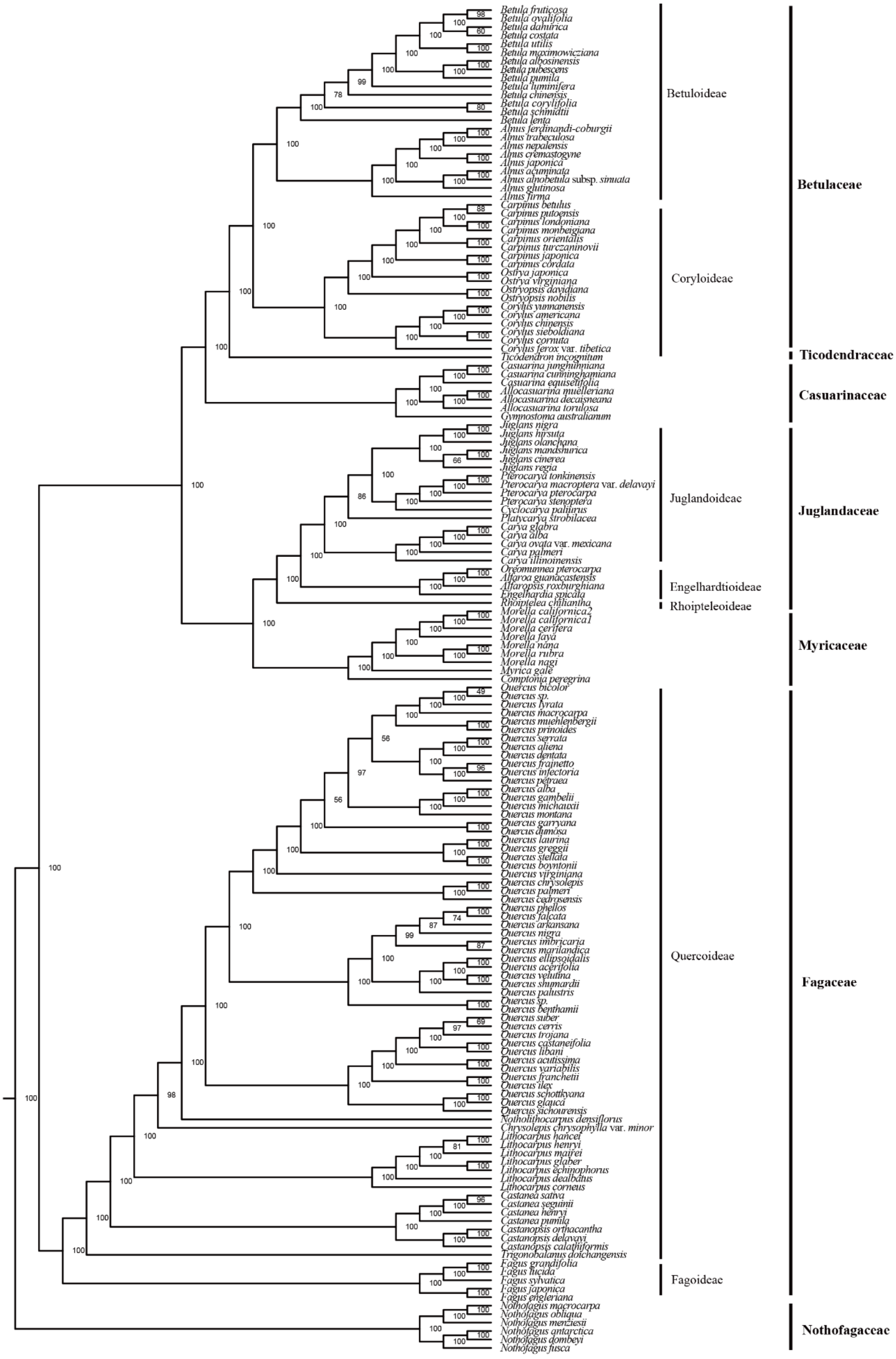

**Fig. S4 The MQSST inferred from 643 gene trees.**

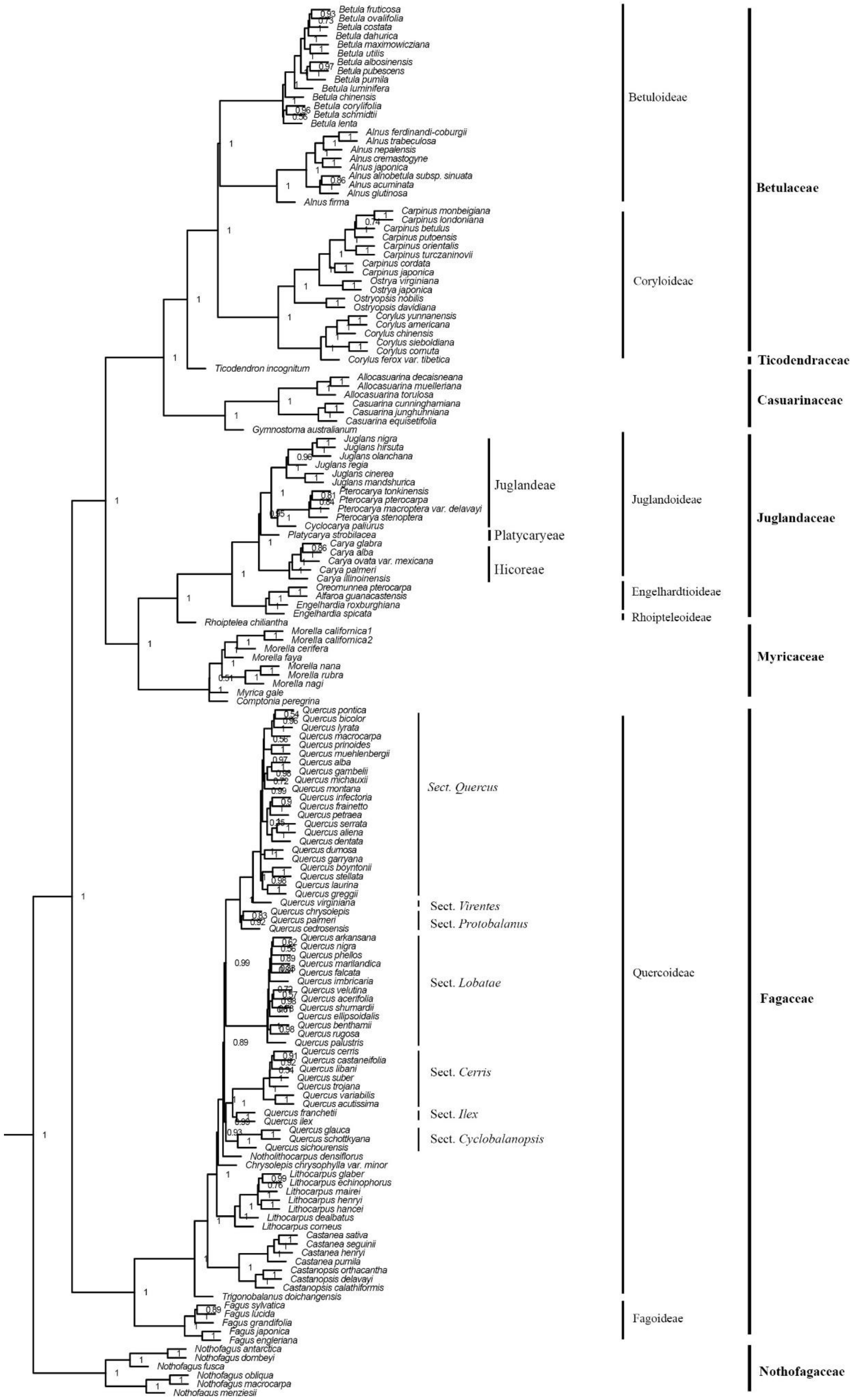

**Fig. S5 Variation in divergence-time estimates.** The trees shown are the 30 independently dated ML trees, plotted using the 'densiTree' function in phangorn. The individual ML trees were dated using treePL.

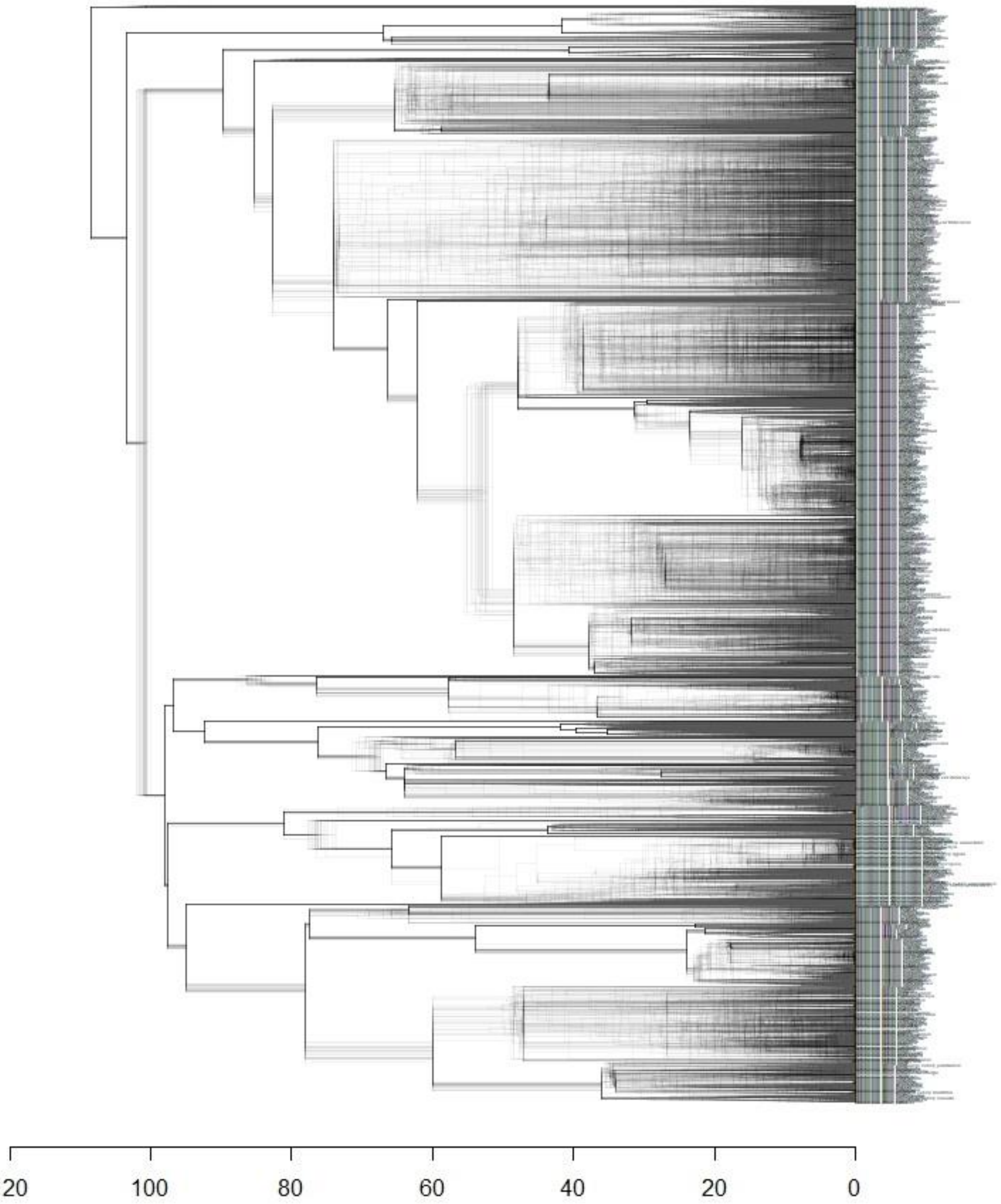

**Fig. S6 Concordance analysis using the 643 gene trees and the CML tree.** Pie charts depict conflict among the input trees, with the blue, green, red, and gray slices representing, respectively, the proportion of bipartitions that are concordant, conflicting (supporting a single main alternative topology), conflicting (supporting various alternative topologies), or uninformative (BS < 70% or missing taxon) at each node in the CML tree. The numbers above and below each branch are, respectively, the number of bipartitions concordant and conflicting with that particular node.

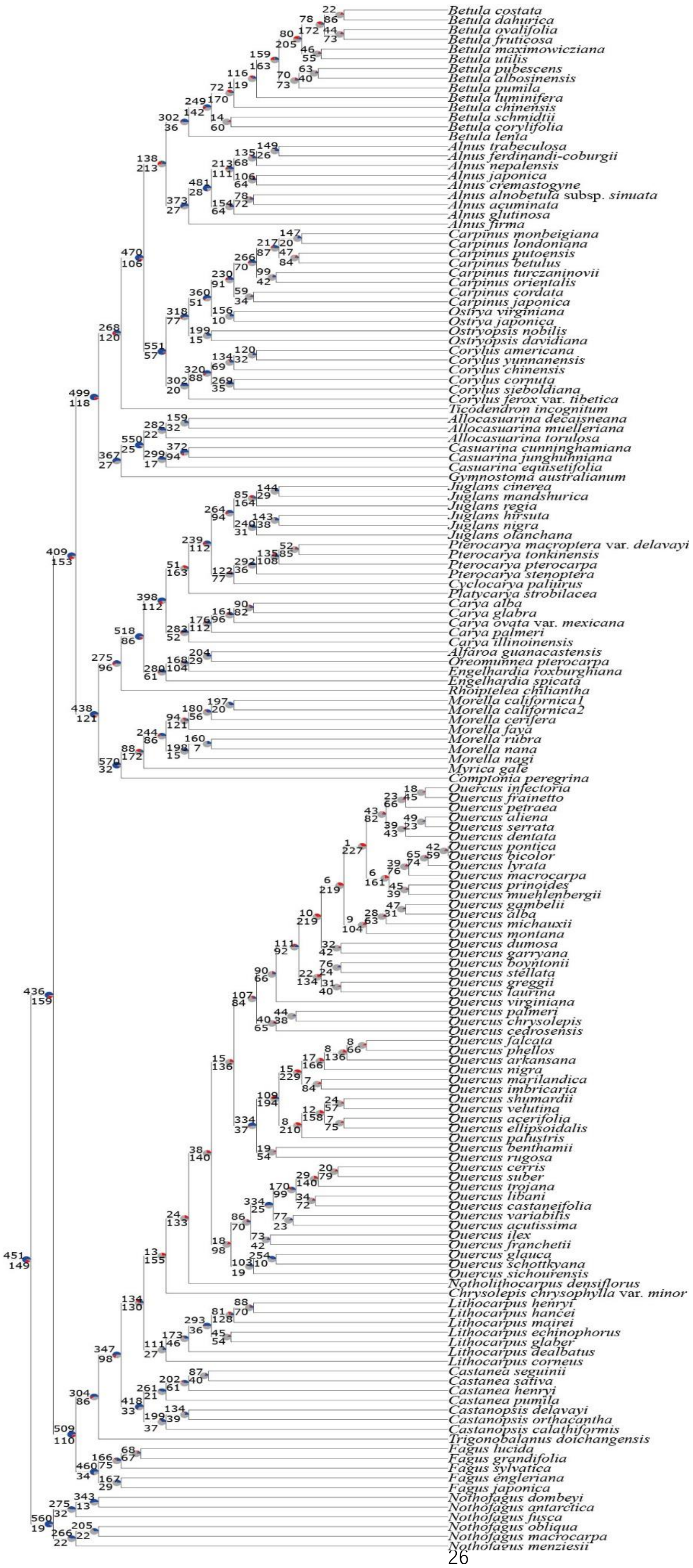
Betula costata
Betula dahurica
Betula ovalifolia
Betula fruticosa
Betula maximowicziana
Betula utilis
Betula pubescens
Betula albosinensis
Betula pumila
Betula luminifera
Betula chinensis
Betula schmidtii
Betula corylifolia
Betula lenta
Alnus trabeculosa
Alnus ferdinandi-coburgii
Alnus nepalensis
Alnus japonica
Alnus cremastogyne
Alnus alnobetula subsp. sinuata
Alnus acuminata
Alnus glutinosa
Alnus firma
Carpinus monbeigiana
Carpinus londoniana
Carpinus putoensis
Carpinus betulus
Carpinus turczaninovii
Carpinus orientalis
Carpinus cordata
Carpinus japonica
Ostrya virginiana
Ostrya japonica
Ostryopsis nobilis
Ostryopsis davidiana
Corylus americana
Corylus yunnanensis
Corylus chinensis
Corylus cornuta
Corylus sieboldiana
Corylus ferox var. tibetica
Ticodendron incognitum
Allocasuarina decaisneana
Allocasuarina muelleriana
Allocasuarina torulosa
Casuarina cunninghamiana
Casuarina junghuhniana
Casuarina equisetifolia
Gymnostoma australianum
Juglans cinerea
Juglans mandshurica
Juglans regia
Juglans hirsuta
Juglans nigra
Juglans olanchana
Pterocarya macroptera var. delavayi
Pterocarya tonkinensis
Pterocarya pterocarpa
Pterocarya stenoptera
Cyclocarya paliurus
Platycarya strobilacea
Carya alba
Carya glabra
Carya ovata var. mexicana
Carya palmeri
Carya illinoinensis
Alfaroa guanacastensis
Oreomunnea pterocarpa
Engelhardia roxburghiana
Engelhardia spicata
Rhoiptelea chiliantha
Morella californica1
Morella californica2
Morella cerifera
Morella faya
Morella rubra
Morella nana
Morella nagi
Myrica gale
Comptonia peregrina
Quercus infectoria
Quercus frainetto
Quercus petraea
Quercus aliena
Quercus serrata
Quercus dentata
Quercus pontica
Quercus bicolor
Quercus lyrata
Quercus macrocarpa
Quercus prinoides
Quercus muehlenbergii
Quercus gambelii
Quercus alba
Quercus michauxii
Quercus montana
Quercus dumosa
Quercus garryana
Quercus boyntonii
Quercus stellata
Quercus greggii
Quercus laurina
Quercus virginiana
Quercus palmeri
Quercus chrysolepis
Quercus cedrosensis
Quercus falcata
Quercus phellos
Quercus arkansana
Quercus nigra
Quercus marilandica
Quercus imbricaria
Quercus shumardii
Quercus velutina
Quercus acerifolia
Quercus ellipsoidalis
Quercus palustris
Quercus benthamii
Quercus rugosa
Quercus cerris
Quercus suber
Quercus trojana
Quercus libani
Quercus castaneifolia
Quercus variabilis
Quercus acutissima
Quercus ilex
Quercus franchetii
Quercus glauca
Quercus schottkyana
Quercus sichourensis
Notholithocarpus densiflorus
Chrysolepis chrysophylla var. minor
Lithocarpus henryi
Lithocarpus hancei
Lithocarpus mairei
Lithocarpus echinophorus
Lithocarpus glaber
Lithocarpus dealbatus
Lithocarpus corneus
Castanea seguinii
Castanea sativa
Castanea henryi
Castanea pumila
Castanopsis delavayi
Castanopsis orthacantha
Castanopsis calathiformis
Trigonobalanus doichangensis
Fagus lucida
Fagus grandifolia
Fagus sylvatica
Fagus engleriana
Fagus japonica
Nothofagus dombeyi
Nothofagus antarctica
Nothofagus fusca
Nothofagus obliqua
Nothofagus macrocarpa
Nothofagus menziesii

**Fig. S7 Gene duplication events detected from 11,038 homologous trees.** Duplication events are labeled for branches with proportions of duplicated genes greater than 2%. The diameter of the solid circles is proportional to the number of duplicated genes. The red circle indicates a potential WGD event.

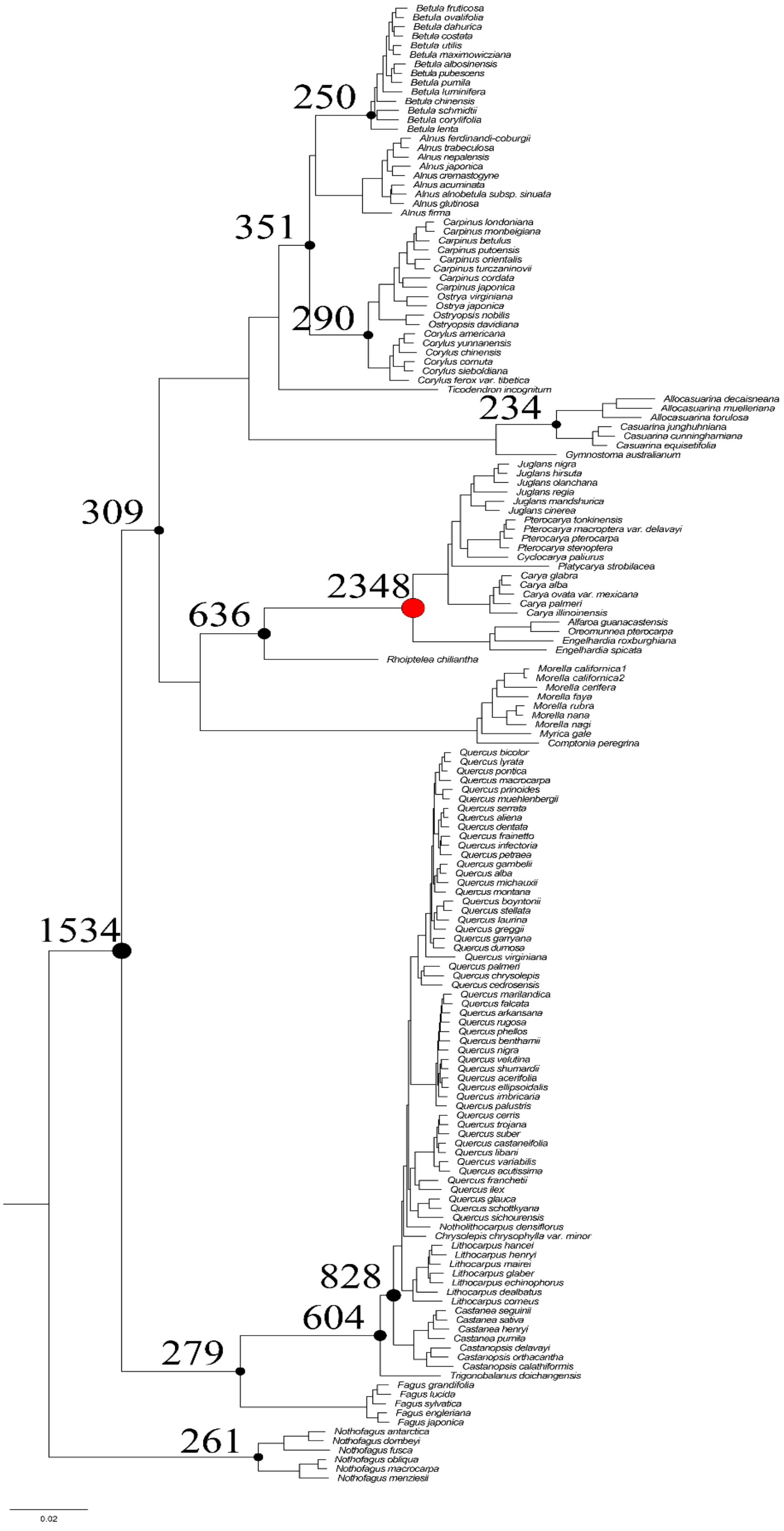

**Fig. S8-1 Distribution of synonymous substitutions (*Ks*) among paralogous gene pairs of Betulaceae.** Peaks around *Ks* = 1.5 represents the early eudicot paleohexaploidy event. *Ks* plots were generated using Perl scripts available at https://github.com/nstenz/plot-ks.

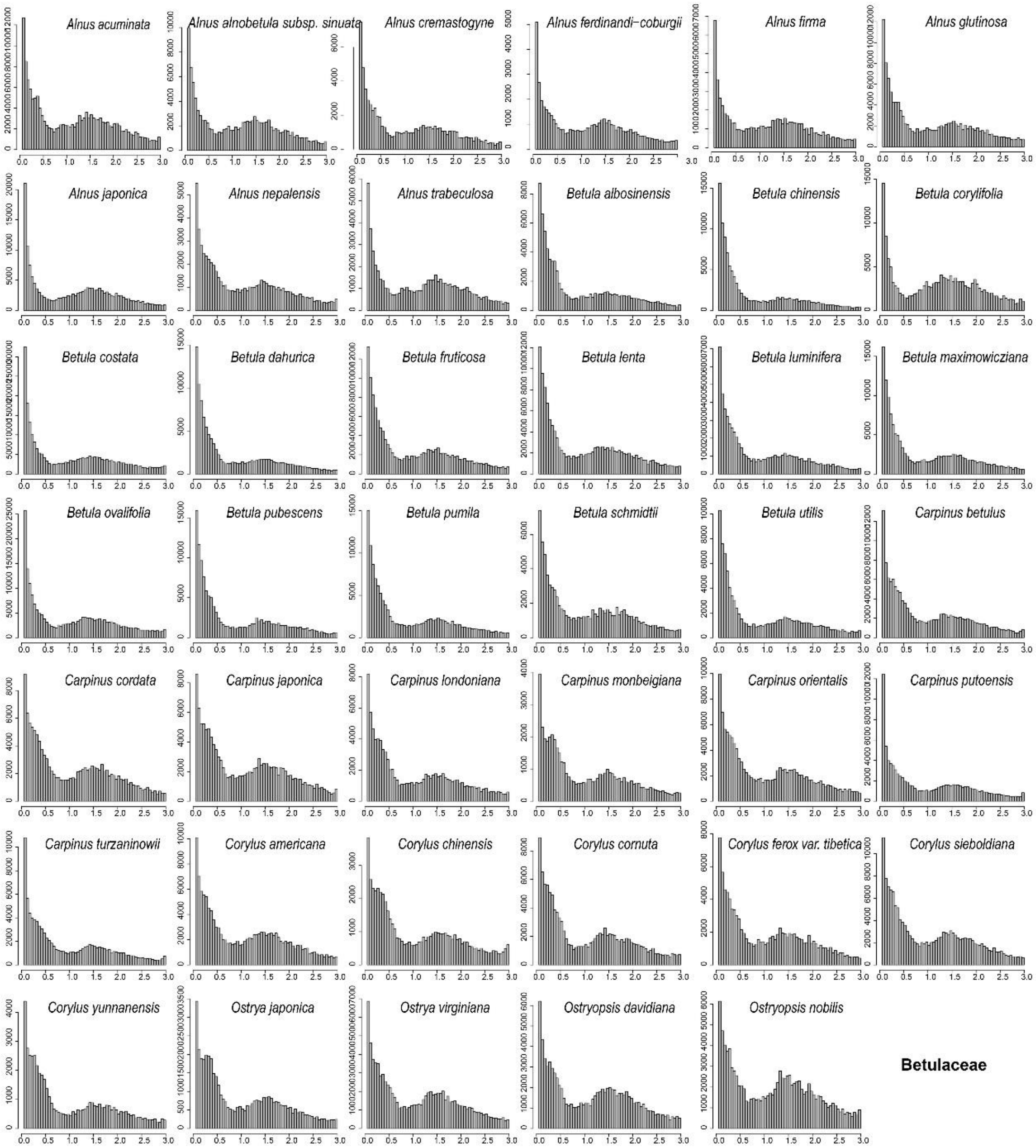

**Fig. S8-2 Distribution of synonymous substitutions (*Ks*) among paralogous gene pairs of Casuarinaceae, Juglandaceae, Myricaceae, and Ticodendraceae.** The Juglandaceae-specific WGD is marked by blue arrows. Peaks around *Ks* = 1.5 represents the early eudicot paleohexaploidy event. *Ks* plots were generated using Perl scripts available at https://github.com/nstenz/plot-ks.

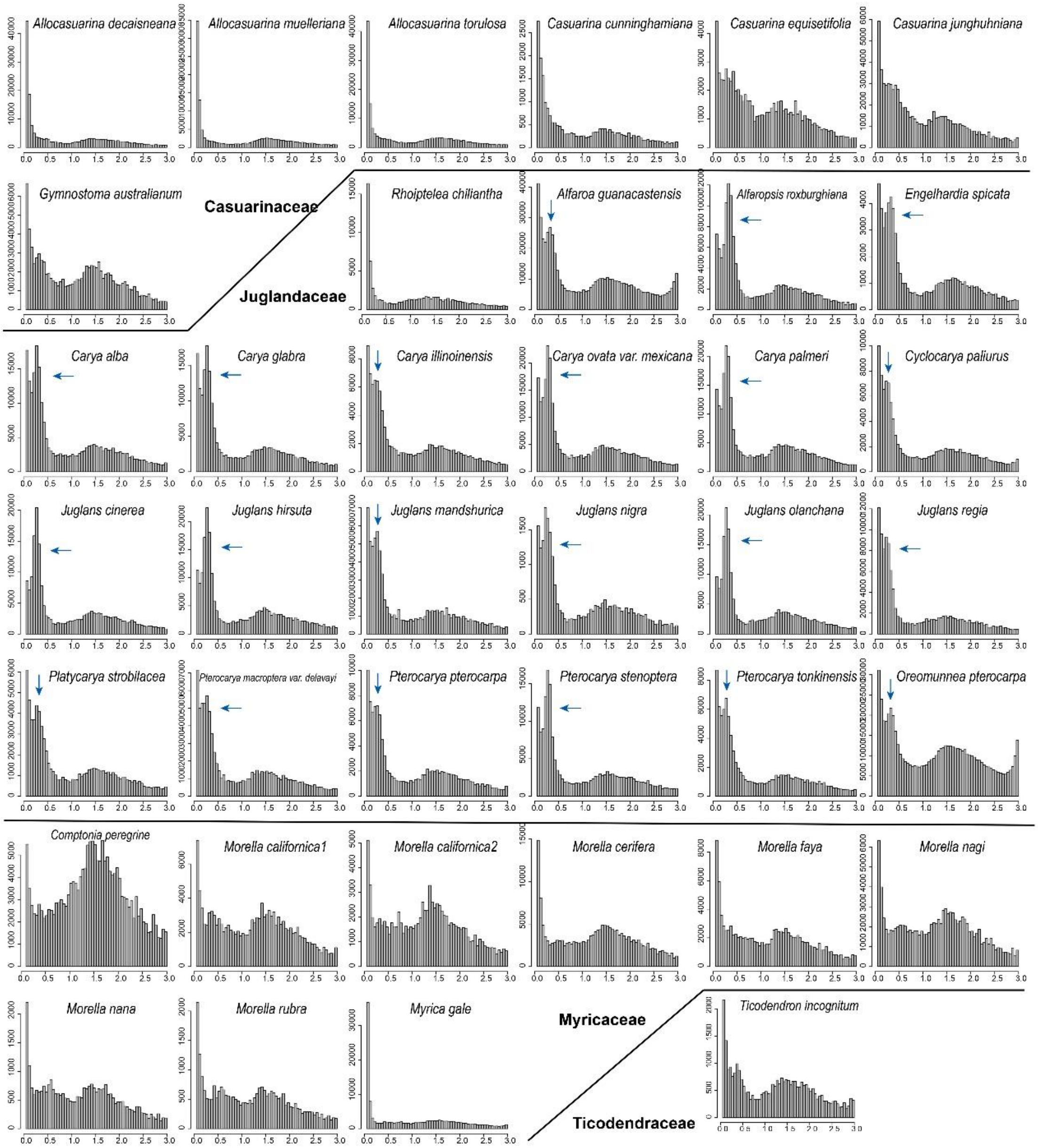

**Fig. S8-3 Distribution of synonymous substitutions (*Ks*) among paralogous gene pairs of Fagaceae.** Peaks around *Ks* = 1.5 represents the early eudicot paleohexaploidy event. *Ks* plots were generated using Perl scripts available at https://github.com/nstenz/plot-ks.

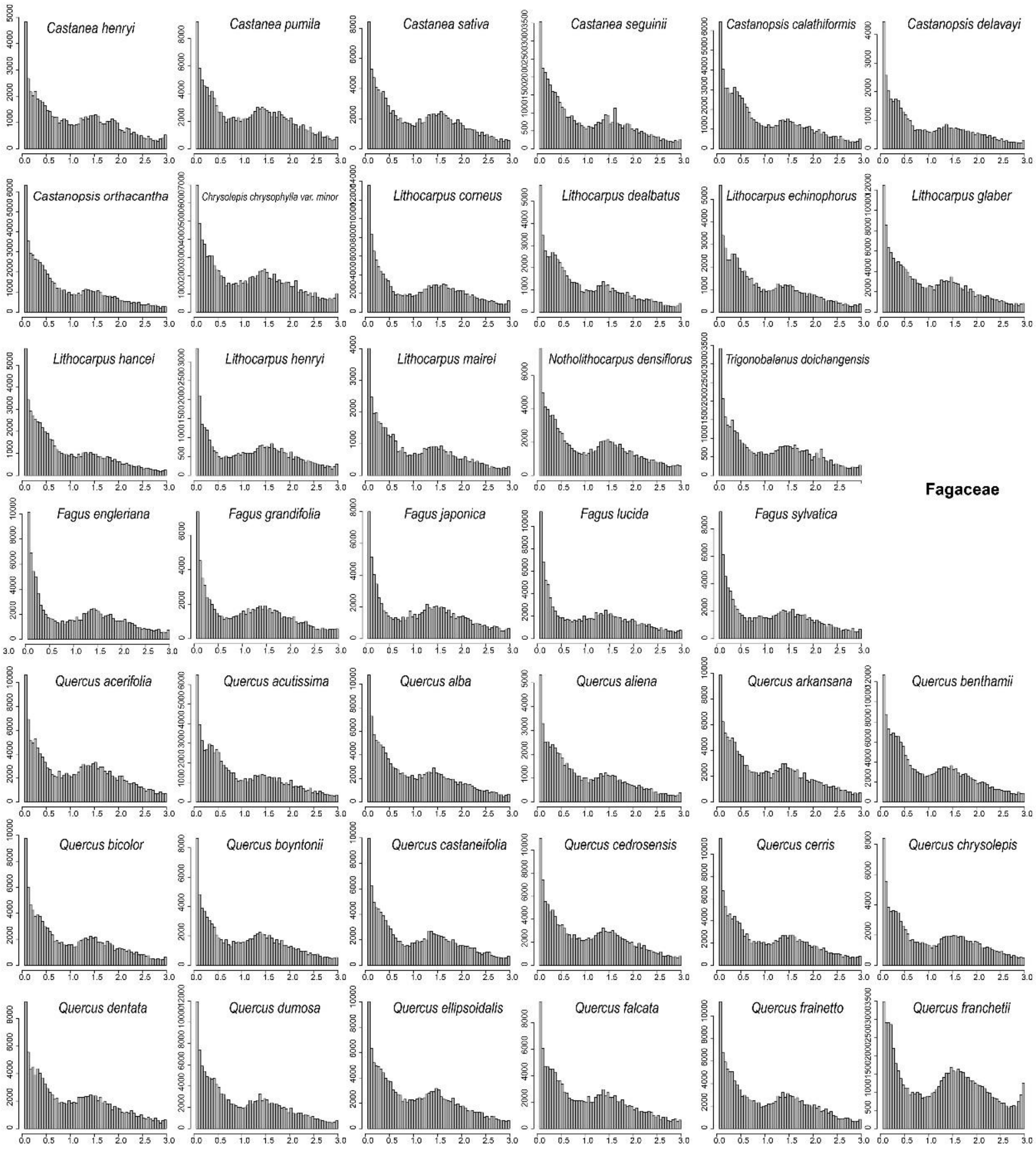

**Fig. S8-4 Distribution of synonymous substitutions (*Ks*) among paralogous gene pairs of Fagaceae and Nothofagaceae.** Peaks around *Ks* = 1.5 represents the early eudicot paleohexaploidy event. *Ks* plots were generated using Perl scripts available at https://github.com/nstenz/plot-ks.

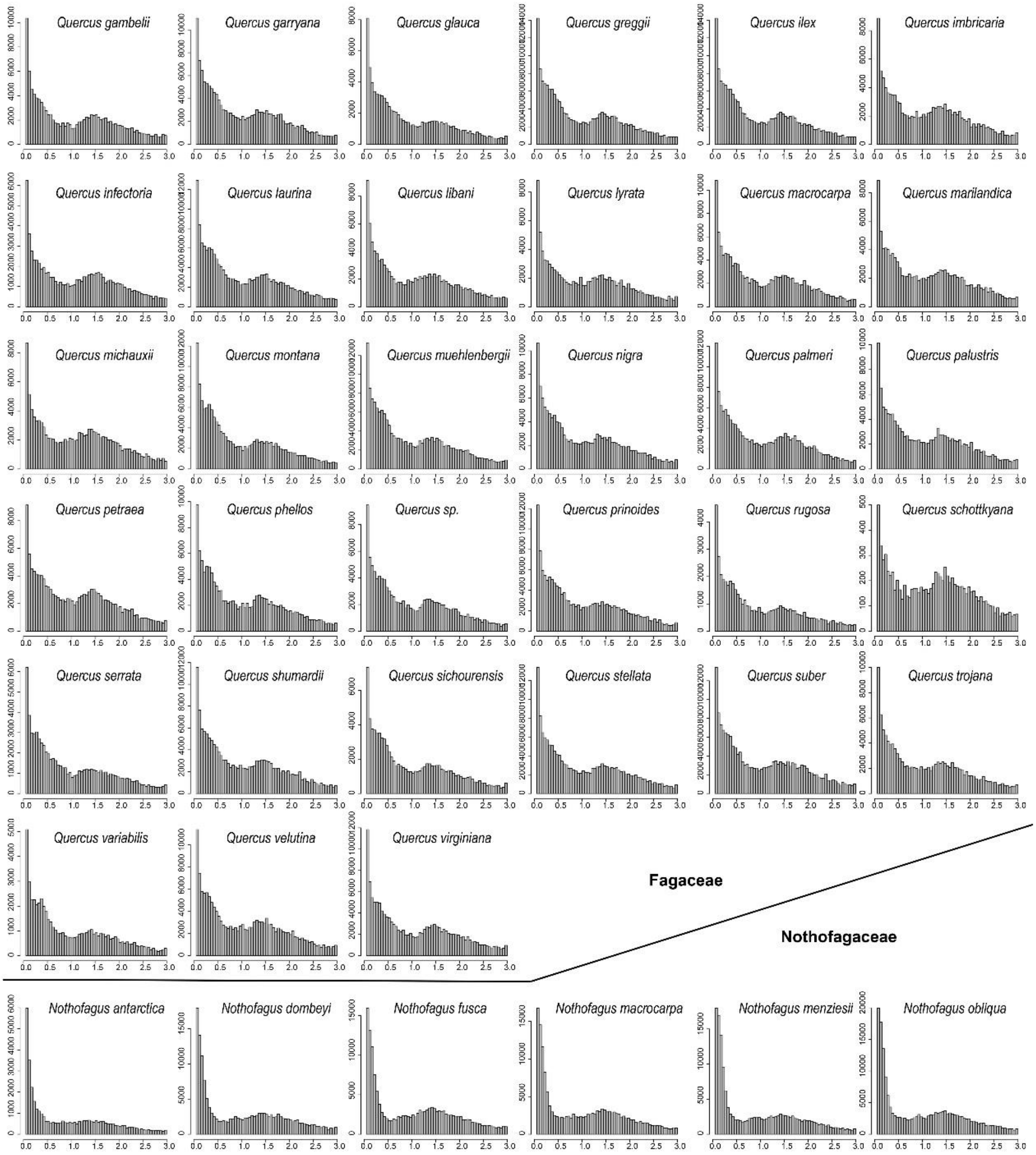

**Fig. S9 Distribution of synonymous substitutions (*Ks*) among paralogous gene pairs of species in Juglandaceae.** The inferred WGD is marked by blue arrows. *Ks* plots were generated using Python scripts available at https://github.com/tanghaibao/bio-pipeline/tree/master/synonymous_calculation.

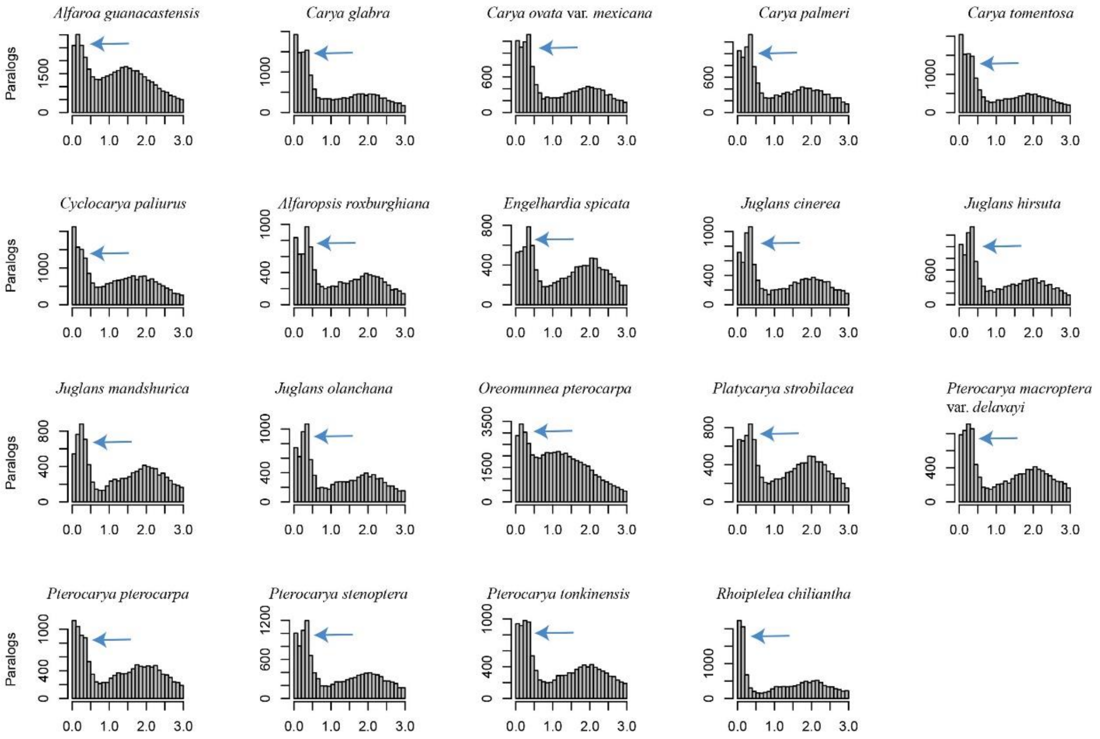

**Fig. S10 *Ks* plots of within-taxon and between-species comparisons**. These show that the divergence of *Rhoiptelea chiliantha* occurred after the WGD, indating that the event is shared across all living members Juglandaceae. *Ks* plots were generated using Python scripts available at https://github.com/tanghaibao/bio-pipeline/tree/master/synonymous_calculation.

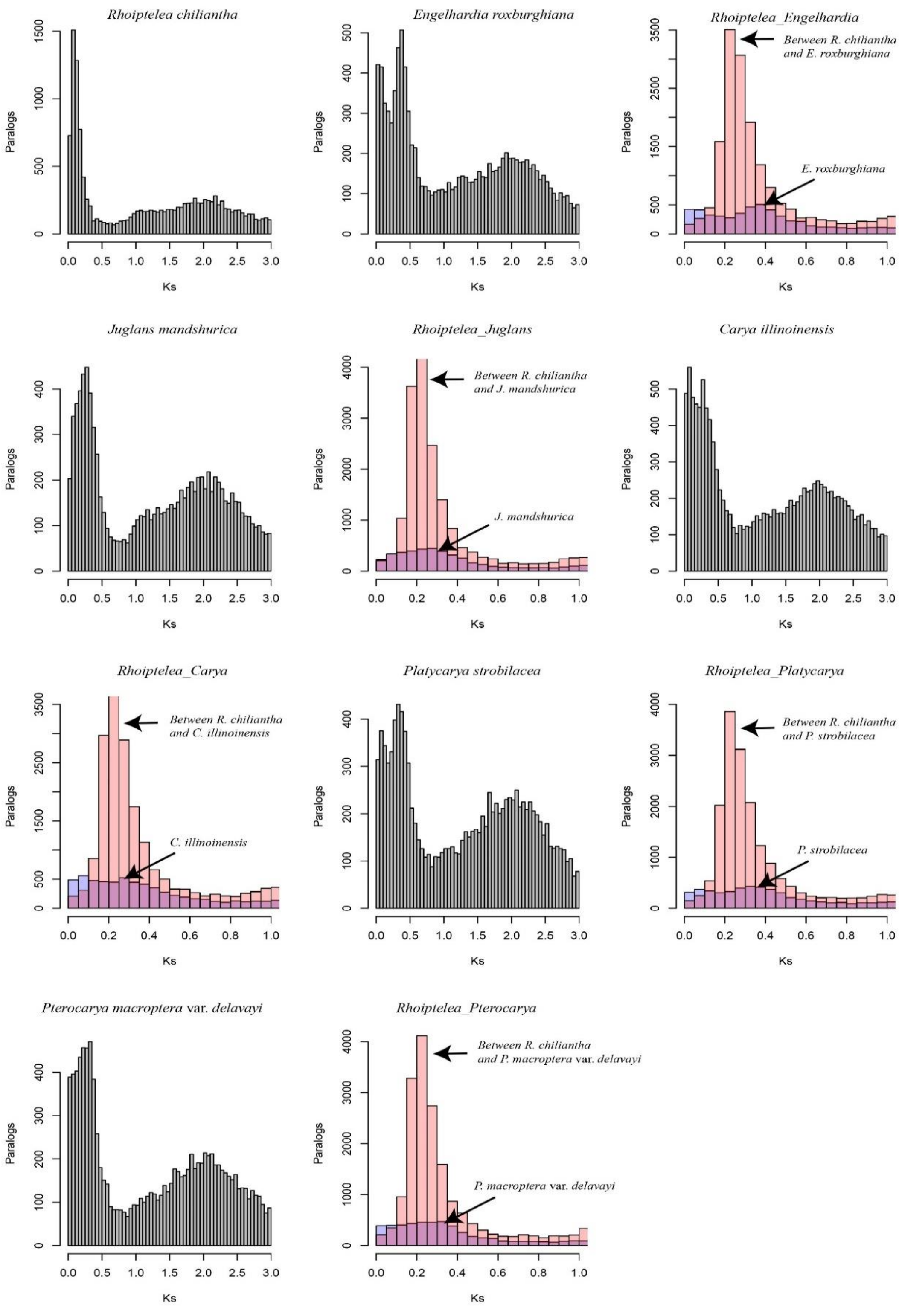

**Fig. S11 Ancestral state reconstruction of the haploid chromosome numbers across Fagales.**

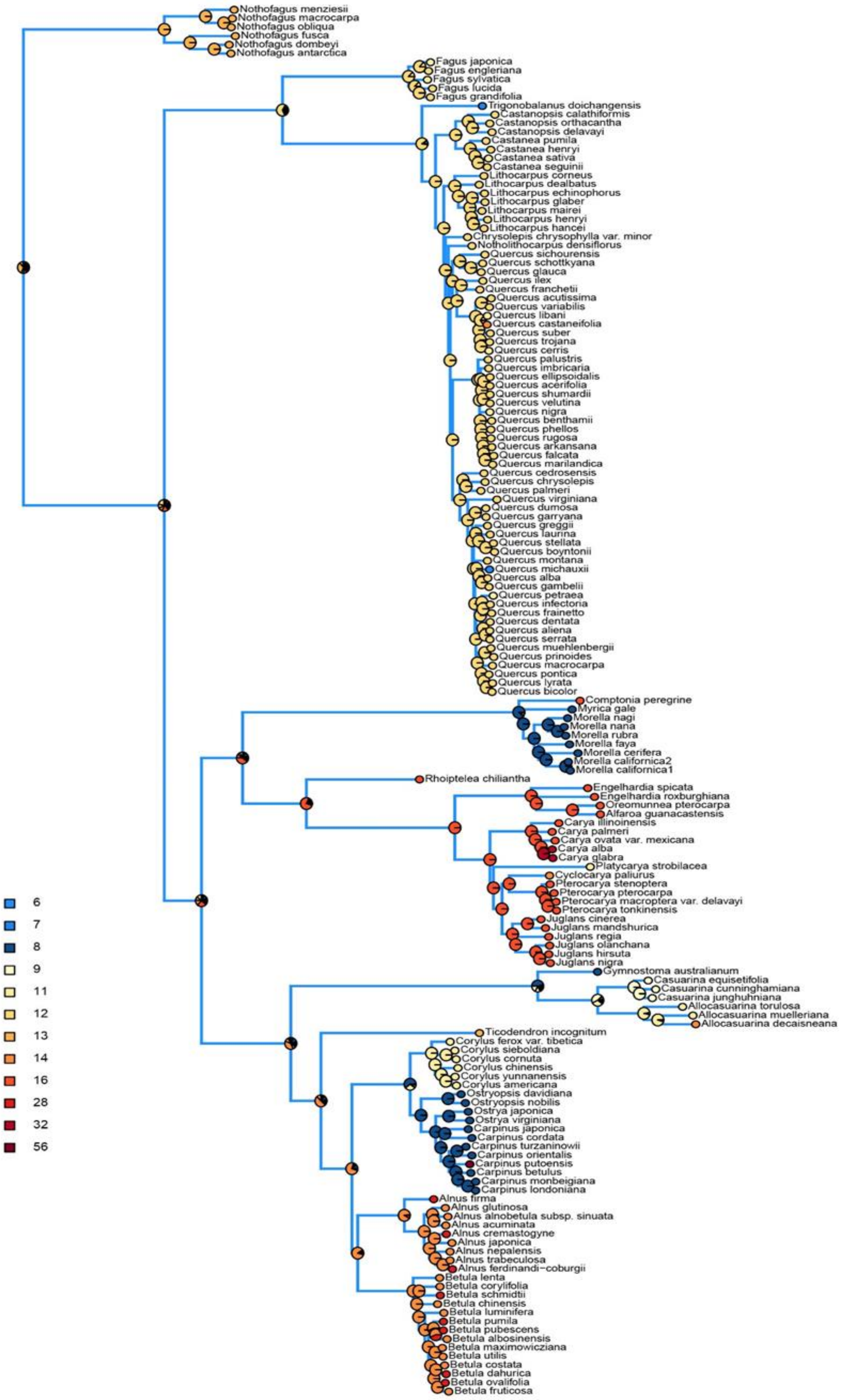

**Fig. S12 Diversification rates in Fagales, with the 11 inferred diversification rate shifts indicated by red dots.**

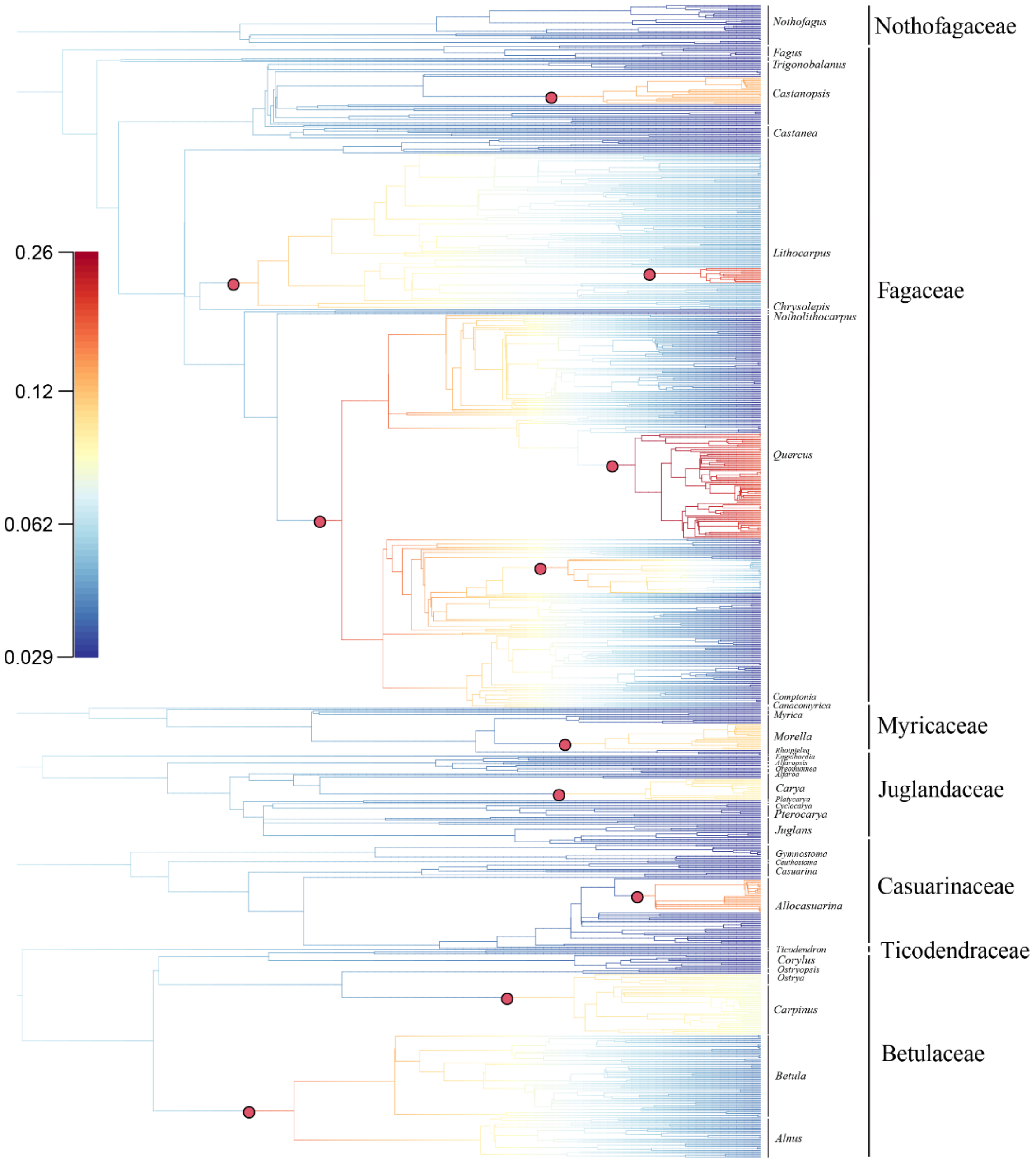

**Fig. S13 Chronophylomorphospace of extant Betulaceae.**

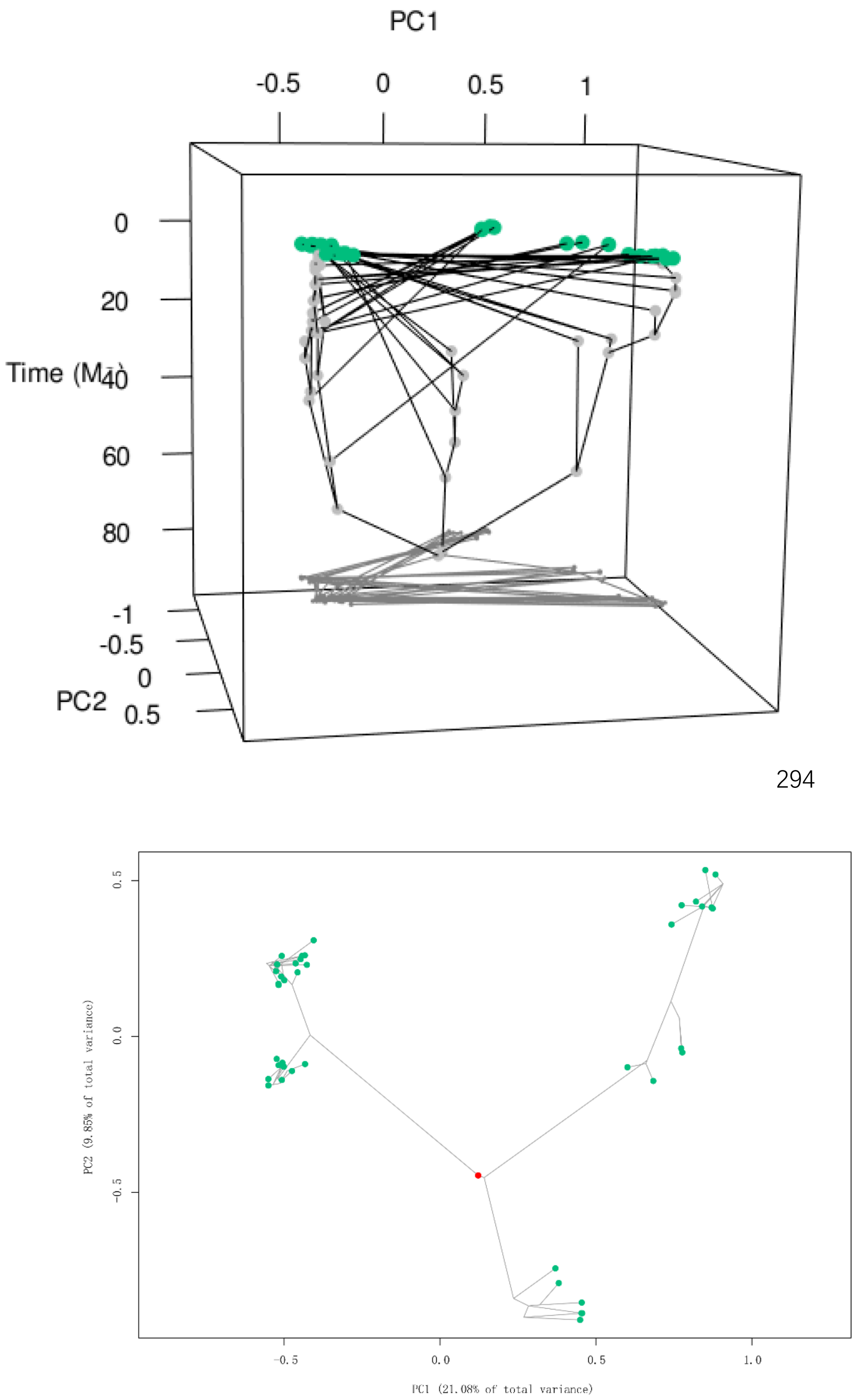

**Fig. S14 Chronophylomorphospace of extant Fagaceae.**

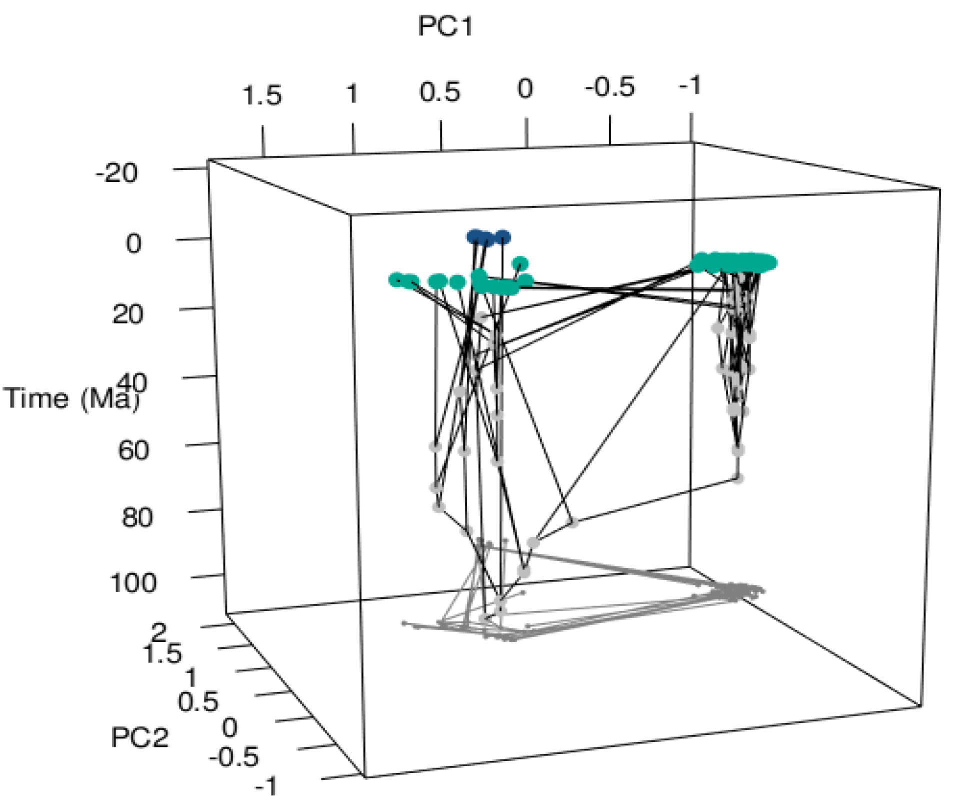


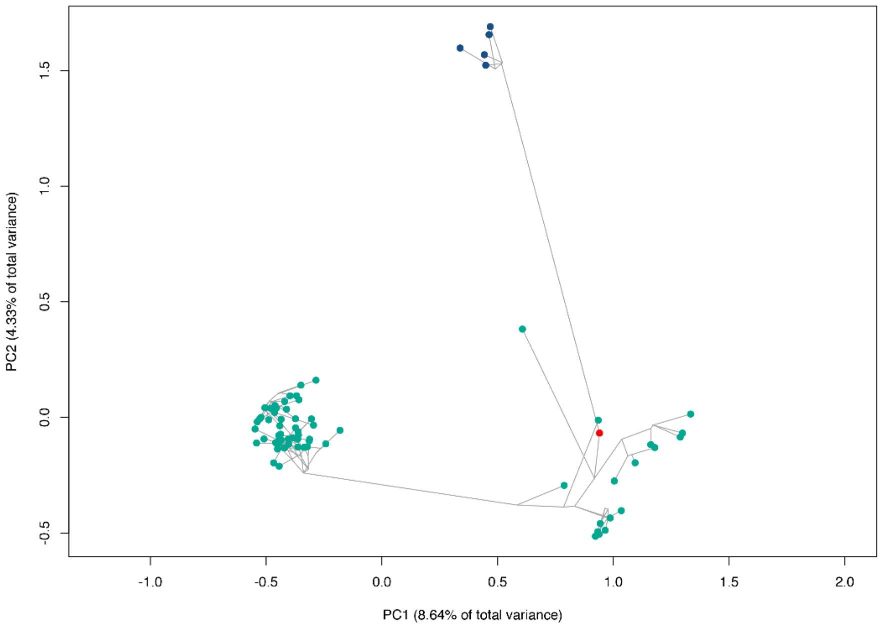

**Fig. S15 Chronophylomorphospace of extant Juglandaceae.**

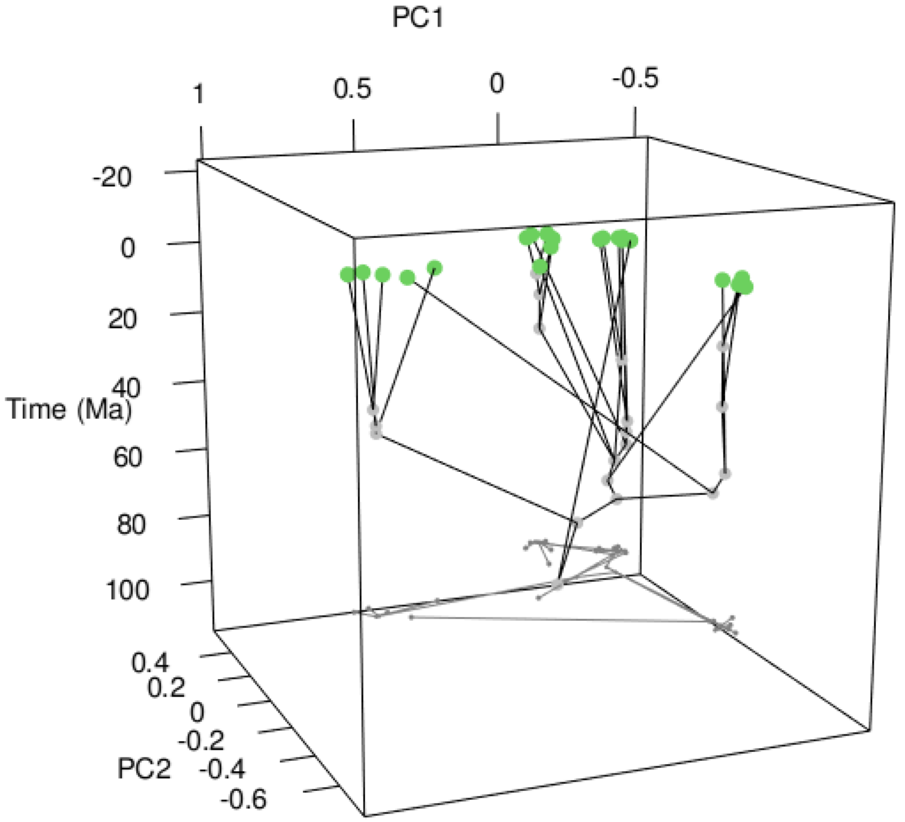


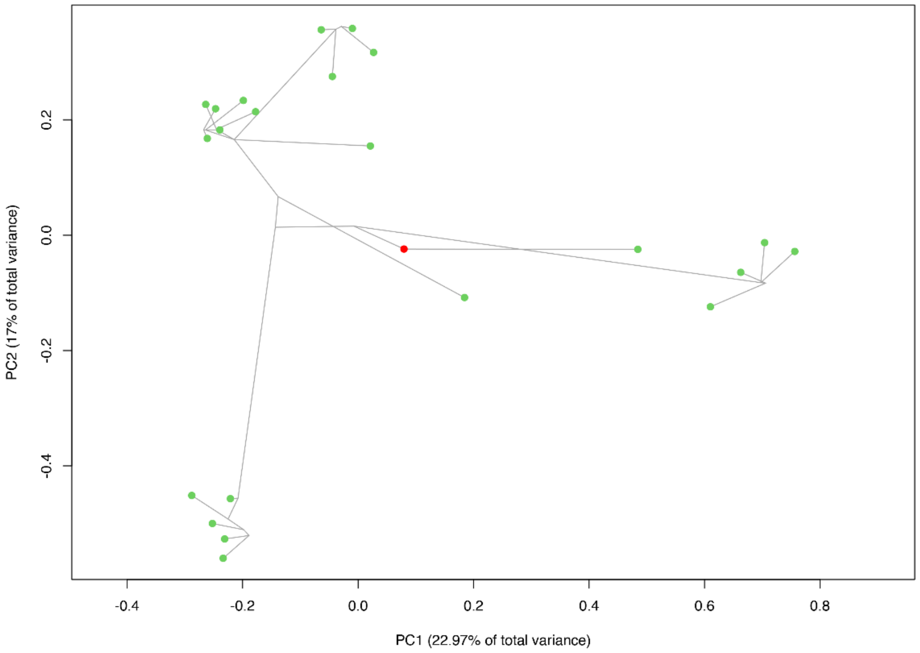

**Fig. S16 Rates of phenotypic evolution across Fagales phylogeny.** Branch lengths correspond to morphological rates inferred using the (A) Mk model or (B) maximum parsimony, following the approach of Parins-Fukuchi et al. (2021). These trees indicate that branches along the backbone of Fagales (subtending modern families and/or major clades) show the greatest rates of phenotypic evolution. On tree (A), stars indicate locations of major evolutionary 'jumps', i.e., significant changes in phenotypic from parent to child nodes, based on ancestral state reconstructions of the first two axes of a principal coordinates analysis.

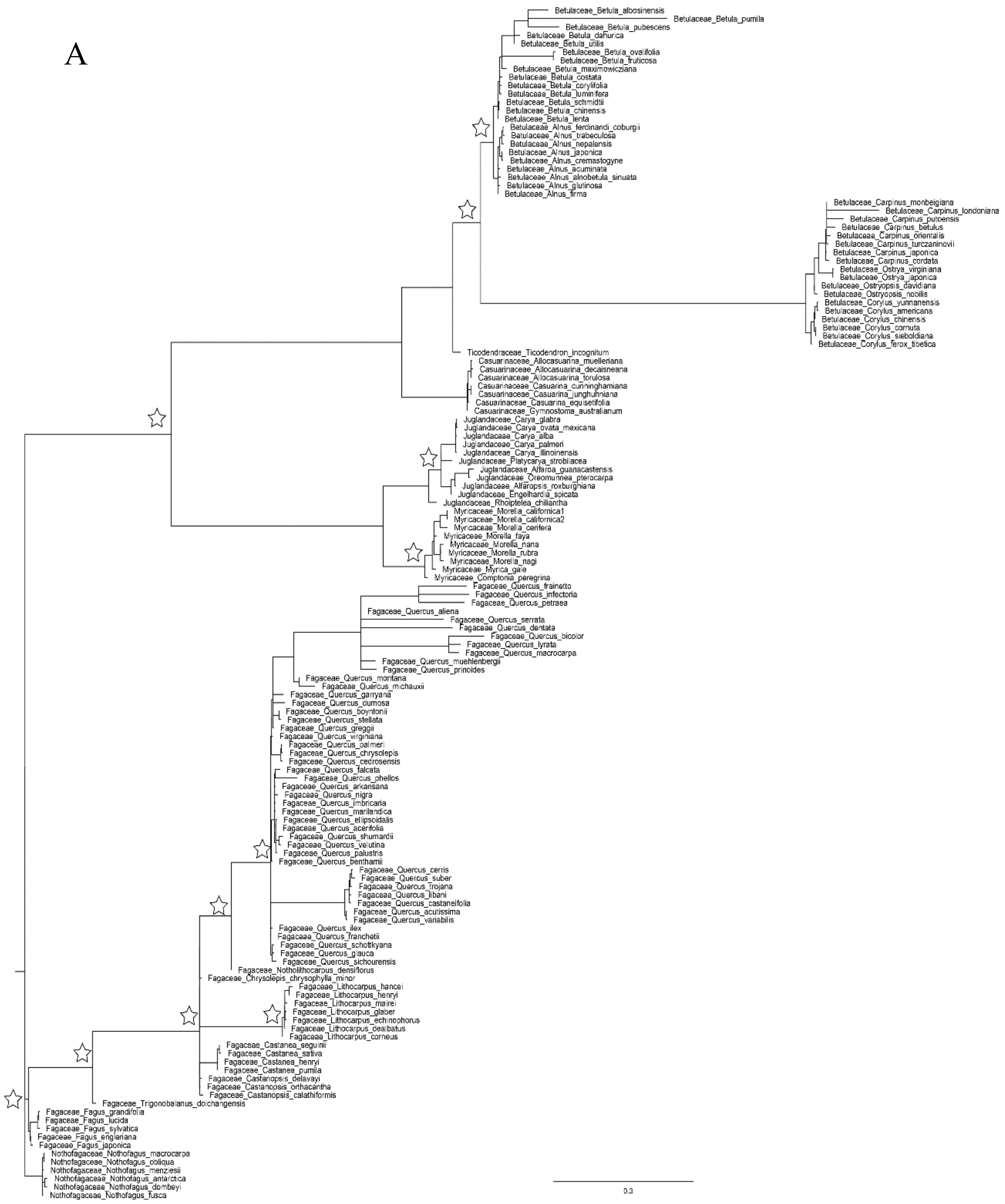

B

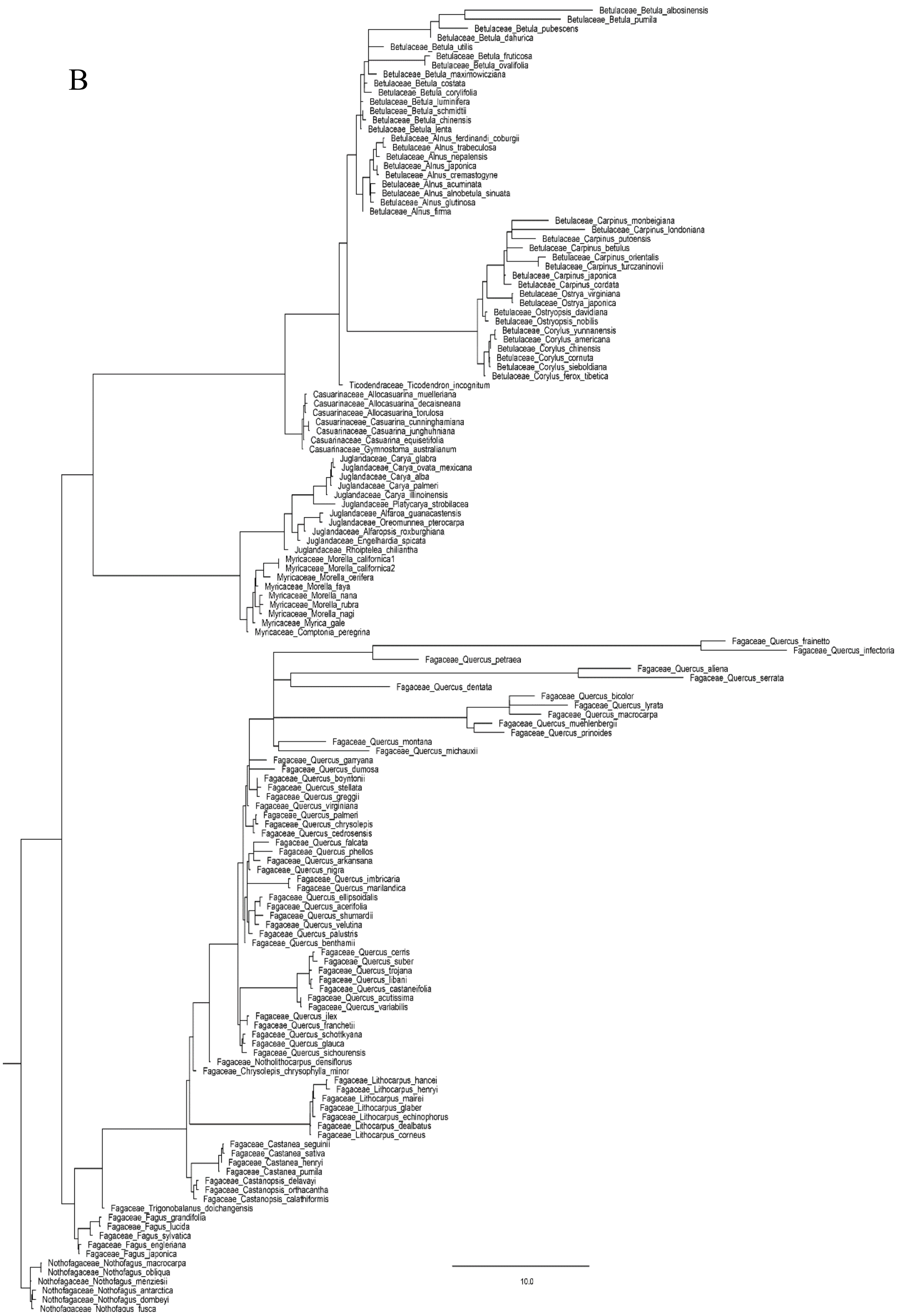

Betulaceae_Betula_albosinensis
Betulaceae_Betula_pumila
Betulaceae_Betula_pubescens
Betulaceae_Betula_dahurica
Betulaceae_Betula_utilis
Betulaceae_Betula_fruticosa
Betulaceae_Betula_ovalifolia
Betulaceae_Betula_maximowicziana
Betulaceae_Betula_costata
Betulaceae_Betula_corylifolia
Betulaceae_Betula_luminifera
Betulaceae_Betula_schmidtii
Betulaceae_Betula_chinensis
Betulaceae_Betula_lenta
Betulaceae_Alnus_ferdinandi_coburgii
Betulaceae_Alnus_trabeculosa
Betulaceae_Alnus_nepalensis
Betulaceae_Alnus_japonica
Betulaceae_Alnus_cremastogyne
Betulaceae_Alnus_acuminata
Betulaceae_Alnus_alnobetula_sinuata
Betulaceae_Alnus_glutinosa
Betulaceae_Alnus_firma
Betulaceae_Carpinus_monbeigiana
Betulaceae_Carpinus_londoniana
Betulaceae_Carpinus_putoensis
Betulaceae_Carpinus_betulus
Betulaceae_Carpinus_orientalis
Betulaceae_Carpinus_turczaninovii
Betulaceae_Carpinus_japonica
Betulaceae_Carpinus_cordata
Betulaceae_Ostrya_virginiana
Betulaceae_Ostrya_japonica
Betulaceae_Ostryopsis_davidiana
Betulaceae_Ostryopsis_nobilis
Betulaceae_Corylus_yunnanensis
Betulaceae_Corylus_americana
Betulaceae_Corylus_chinensis
Betulaceae_Corylus_cornuta
Betulaceae_Corylus_sieboldiana
Betulaceae_Corylus_ferox_tibetica
Ticodendraceae_Ticodendron_incognitum
Casuarinaceae_Allocasuarina_muelleriana
Casuarinaceae_Allocasuarina_decaisneana
Casuarinaceae_Allocasuarina_torulosa
Casuarinaceae_Casuarina_cunninghamiana
Casuarinaceae_Casuarina_junghuhniana
Casuarinaceae_Casuarina_equisetifolia
Casuarinaceae_Gymnostoma_australianum
Juglandaceae_Carya_glabra
Juglandaceae_Carya_ovata_mexicana
Juglandaceae_Carya_alba
Juglandaceae_Carya_palmeri
Juglandaceae_Carya_illinoinensis
Juglandaceae_Platycarya_strobilacea
Juglandaceae_Alfaroa_guanacastensis
Juglandaceae_Oreomunnea_pterocarpa
Juglandaceae_Alfaropsis_roxburghiana
Juglandaceae_Engelhardia_spicata
Juglandaceae_Rhoiptelea_chiliantha
Myricaceae_Morella_californica1
Myricaceae_Morella_californica2
Myricaceae_Morella_cerifera
Myricaceae_Morella_faya
Myricaceae_Morella_nana
Myricaceae_Morella_rubra
Myricaceae_Morella_nagi
Myricaceae_Myrica_gale
Myricaceae_Comptonia_peregrina
Fagaceae_Quercus_frainetto
Fagaceae_Quercus_infectoria
Fagaceae_Quercus_petraea
Fagaceae_Quercus_aliena
Fagaceae_Quercus_serrata
Fagaceae_Quercus_dentata
Fagaceae_Quercus_bicolor
Fagaceae_Quercus_lyrata
Fagaceae_Quercus_macrocarpa
Fagaceae_Quercus_muehlenbergii
Fagaceae_Quercus_prinoides
Fagaceae_Quercus_montana
Fagaceae_Quercus_michauxii
Fagaceae_Quercus_garryana
Fagaceae_Quercus_dumosa
Fagaceae_Quercus_boyntonii
Fagaceae_Quercus_stellata
Fagaceae_Quercus_greggii
Fagaceae_Quercus_virginiana
Fagaceae_Quercus_palmeri
Fagaceae_Quercus_chrysolepis
Fagaceae_Quercus_cedrosensis
Fagaceae_Quercus_falcata
Fagaceae_Quercus_phellos
Fagaceae_Quercus_arkansana
Fagaceae_Quercus_nigra
Fagaceae_Quercus_imbricaria
Fagaceae_Quercus_marilandica
Fagaceae_Quercus_ellipsoidalis
Fagaceae_Quercus_acerifolia
Fagaceae_Quercus_shumardii
Fagaceae_Quercus_velutina
Fagaceae_Quercus_palustris
Fagaceae_Quercus_benthamii
Fagaceae_Quercus_cerris
Fagaceae_Quercus_suber
Fagaceae_Quercus_trojana
Fagaceae_Quercus_libani
Fagaceae_Quercus_castaneifolia
Fagaceae_Quercus_acutissima
Fagaceae_Quercus_variabilis
Fagaceae_Quercus_ilex
Fagaceae_Quercus_franchetii
Fagaceae_Quercus_schottkyana
Fagaceae_Quercus_glauca
Fagaceae_Quercus_sichourensis
Fagaceae_Notholithocarpus_densiflorus
Fagaceae_Chrysolepis_chrysophylla_minor
Fagaceae_Lithocarpus_hancei
Fagaceae_Lithocarpus_henryi
Fagaceae_Lithocarpus_mairei
Fagaceae_Lithocarpus_glaber
Fagaceae_Lithocarpus_echinophorus
Fagaceae_Lithocarpus_dealbatus
Fagaceae_Lithocarpus_corneus
Fagaceae_Castanea_seguinii
Fagaceae_Castanea_sativa
Fagaceae_Castanea_henryi
Fagaceae_Castanea_pumila
Fagaceae_Castanopsis_delavayi
Fagaceae_Castanopsis_orthacantha
Fagaceae_Castanopsis_calathiformis
Fagaceae_Trigonobalanus_doichangensis
Fagaceae_Fagus_grandifolia
Fagaceae_Fagus_lucida
Fagaceae_Fagus_sylvatica
Fagaceae_Fagus_engleriana
Fagaceae_Fagus_japonica
Nothofagaceae_Nothofagus_macrocarpa
Nothofagaceae_Nothofagus_obliqua
Nothofagaceae_Nothofagus_menziesii
Nothofagaceae_Nothofagus_antarctica
Nothofagaceae_Nothofagus_dombeyi
Nothofagaceae_Nothofagus_fusca
10.0

**Fig. S17 Rates of phenotypic evolution across Fagales phylogeny inferred by BAMM.** The first axis (A) and second axis of (B) from a principle coordinates analysis (PCoA) showed that phenotypic rates were generally fastest during the initial radiation of Fagales, with red circles showing rate shifts along different lineags.

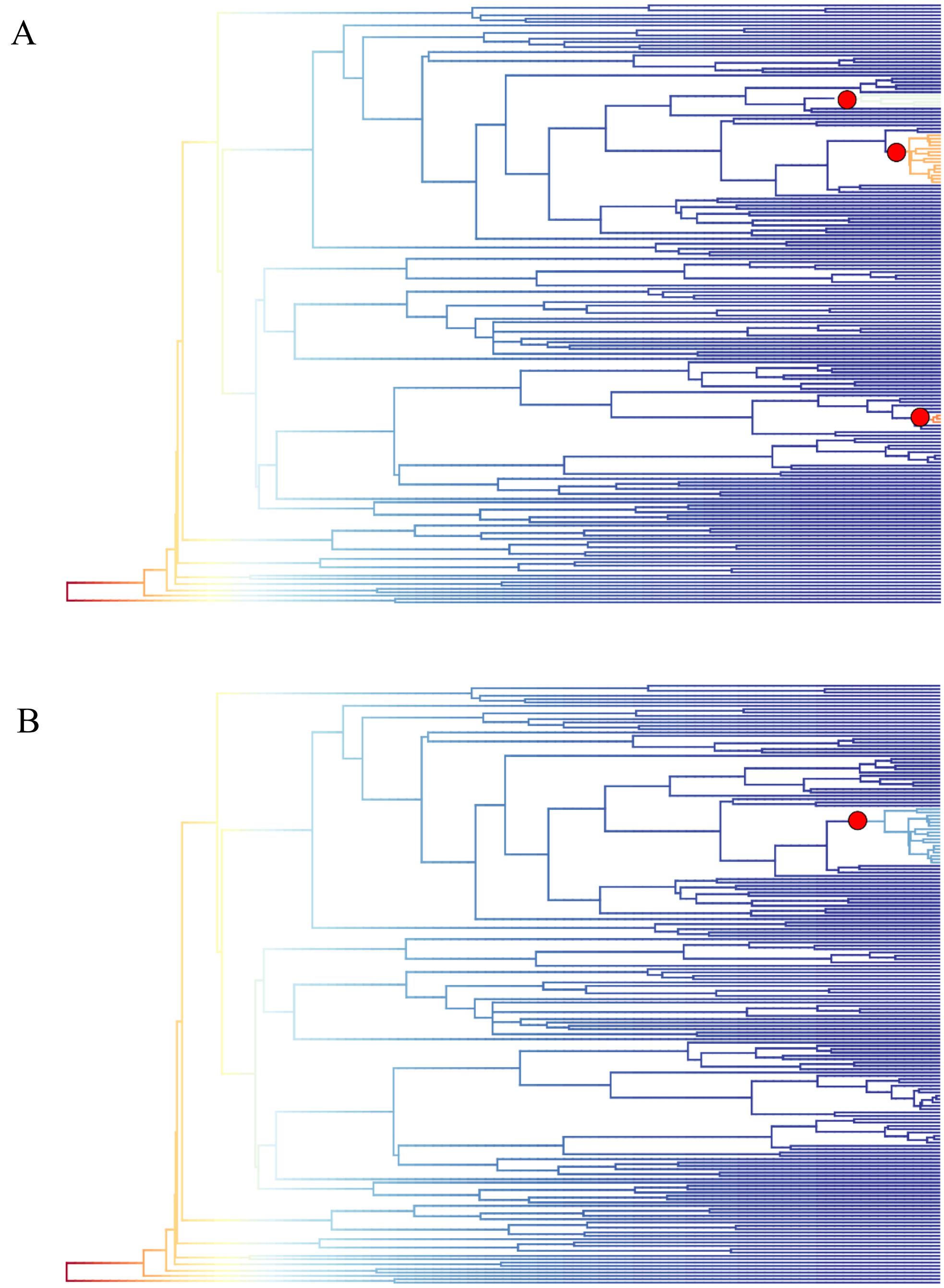

**Fig. S18 Generalized linear regression analysis of phenotypic rates vs. gene duplication levels.** All nodes showed duplications were included in this analysis (n = 88, $r^2 = 0.1786$, P ≈ 0.000).

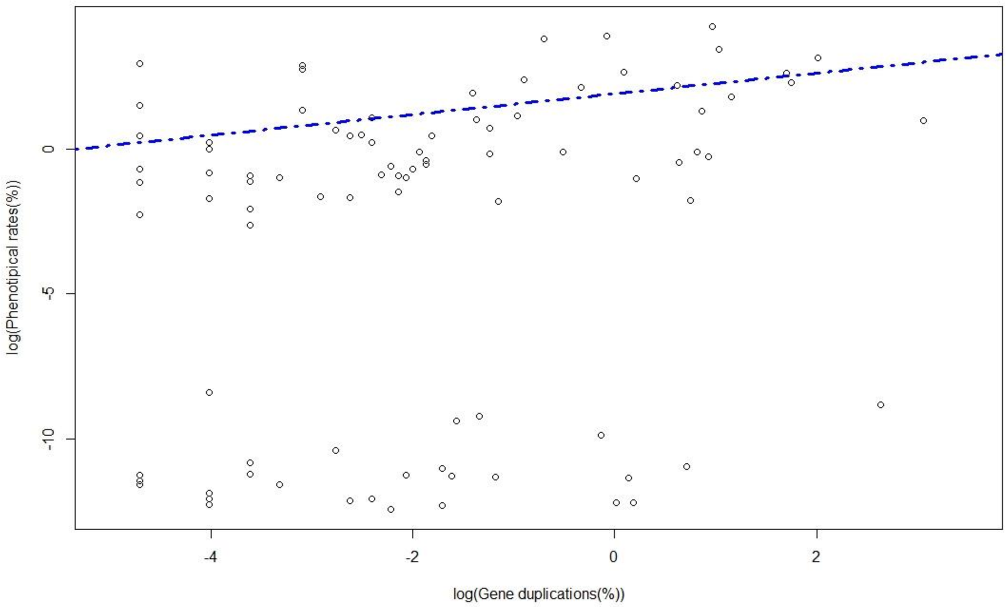


**Fig. S19 Phenotypic innovation versus gene conflicts.** Plots juxtaposing levels of phenotypic innovation (the number of state changes) and levels of gene tree conflict at corresponding nodes across Fagales phylogeny (n = 156). The x axis represents time in millions of years. Each circle represents a node in the phylogeny. Outlier branches are highlighted with colored circles. Betu, Betulaceae; Betu_COO, the clade of *Carpinus* + *Ostrya* + *Ostryopsis*; Casu, Casuarinaceae; Faga, Fagaceae; Faga_Litho, *Lithocarpus*; Faga_Q_Cerr, branch belong to *Quercus* section *Cerris*; Faga_Q_Quercus, *Quercus* section *Quercus* Faga-Fagus, the Fagaceae except *Fagus*; Faga_QNCL, the clade of *Quercus* + *Notholithocarpus* + *Chrysolepis* + *Lithocarpus*; Jugl, Juglandaceae; Jugl-Rhoi, Juglandaceae except *Rhoiptelea*; Jugl_PJPC, the clade of *Pterocarya* + *Juglans* + *Platycarya* + *Carya*; Noth, Nothofagaceae; BTC, he Betulaceae + Ticodendraceae + Casuarinaceae clade; JM, the Juglandaceae + Myricaceae clade; BTCJM, the core Fagales; FBTCJM, Fagales except Nothofagaceae; Myri_MM, the clade of *Myrica* + *Morella* in Myricaceae; crown, the crown node of the lineage.

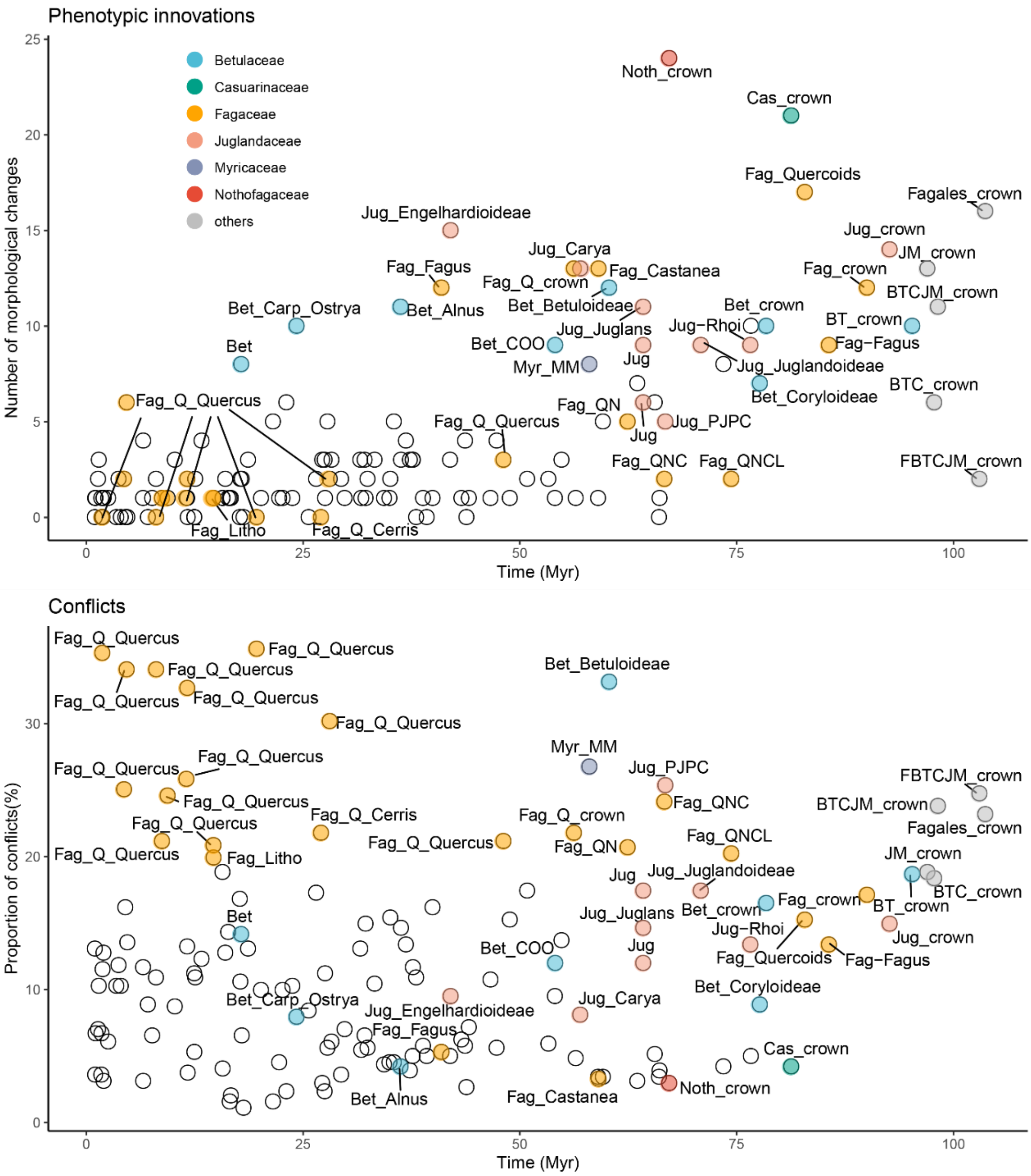
Phenotypic innovations
Betulaceae
Casuarinaceae
Fagaceae
Juglandaceae
Myricaceae
Nothofagaceae
others
Number of morphological changes
Time (Myr)
Noth_crown
Cas_crown
Fag_Quercoids
Fagales_crown
Jug_Engelhardioideae
Jug_crown
JM_crown
Jug_Carya
Fag_Fagus
Fag_Castanea
Fag_Q_crown
Fag_crown
BTCJM_crown
Bet_Carp_Ostrya
Bet_Alnus
Bet_Betuloideae
Bet_crown
Jug_Juglans
Jug−Rhoi
BT_crown
Bet_COO
Fag−Fagus
Bet
Myr_MM
Jug
Jug_Juglandoideae
BTC_crown
Bet_Coryloideae
Fag_Q_Quercus
Fag_QN
Jug_PJPC
Fag_QNC
Fag_QNCL
FBTCJM_crown
Fag_Litho
Fag_Q_Cerris
Conflicts
Proportion of conflicts(%)
Time (Myr)
Fag_Q_Quercus
Bet_Betuloideae
Fag_Q_Quercus
Fag_Q_Quercus
Fag_Q_Quercus
Myr_MM
Jug_PJPC
Fag_Q_Quercus
FBTCJM_crown
Fag_QNC
BTCJM_crown
Fag_Q_Quercus
Fag_Q_Cerris
Fag_Q_crown
Fagales_crown
Fag_Q_Quercus
Fag_Q_Quercus
Fag_QNCL
Fag_QN
JM_crown
Fag_Litho
Jug
Jug_Juglandoideae
Fag_crown
BTC_crown
BT_crown
Bet
Jug_Juglans
Bet_crown
Jug−Rhoi
Jug_crown
Bet_COO
Jug
Fag_Quercoids
Fag−Fagus
Bet_Coryloideae
Bet_Carp_Ostrya
Jug_Carya
Jug_Engelhardioideae
Fag_Fagus
Cas_crown
Noth_crown
Bet_Alnus
Fag_Castanea
0
5
10
15
20
25
30
50
75
100

**Fig. S20 Levels of gene tree conflict versus diversification rates of Fagales.** Each point represents a branch, with the color corresponding to family placement ( n = 143).

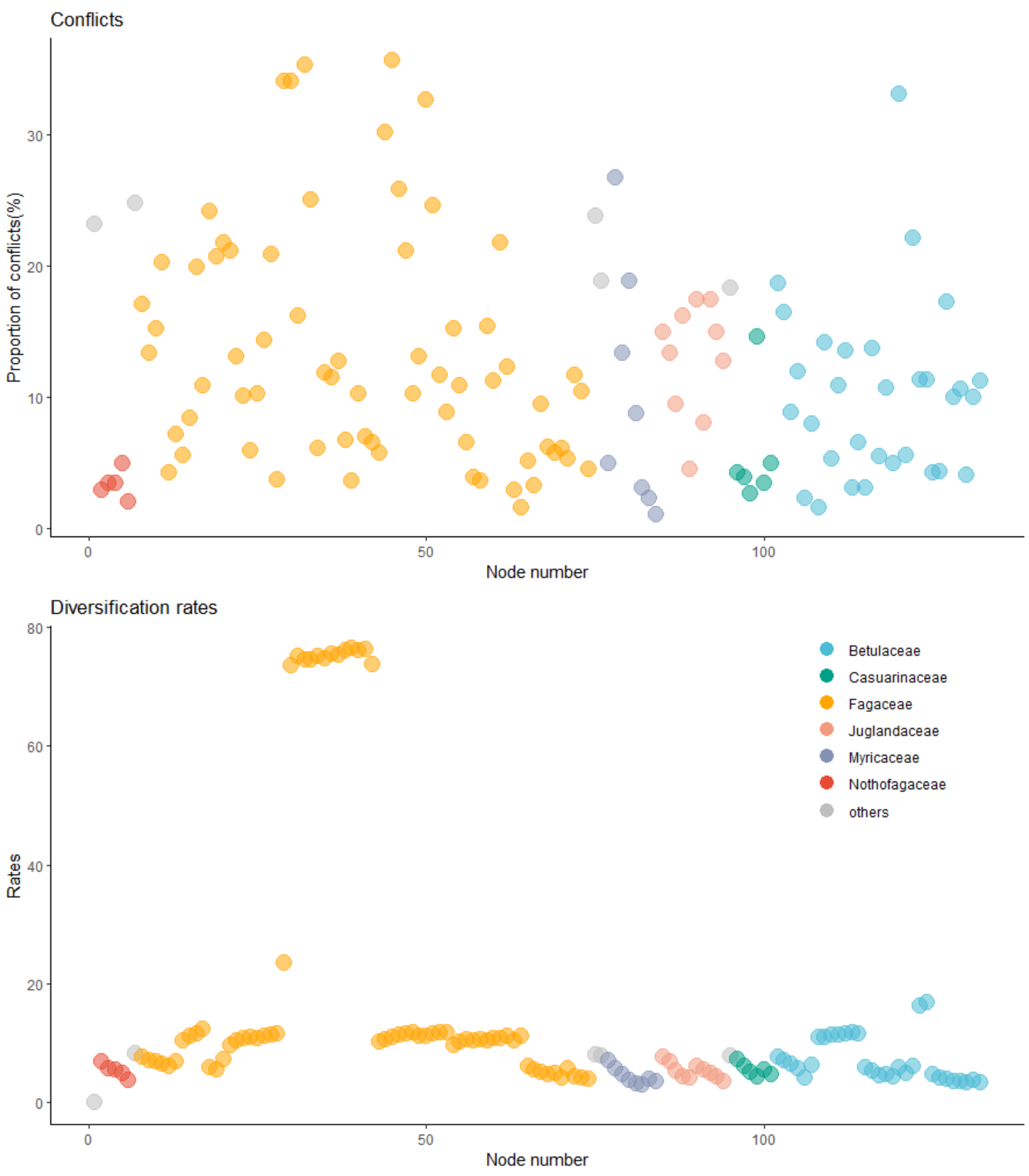

**Fig. S21 Phenotypic innovations versus genes duplications across Fagales.** Each point represents a branch, with the color corresponding to family placement (n = 156).

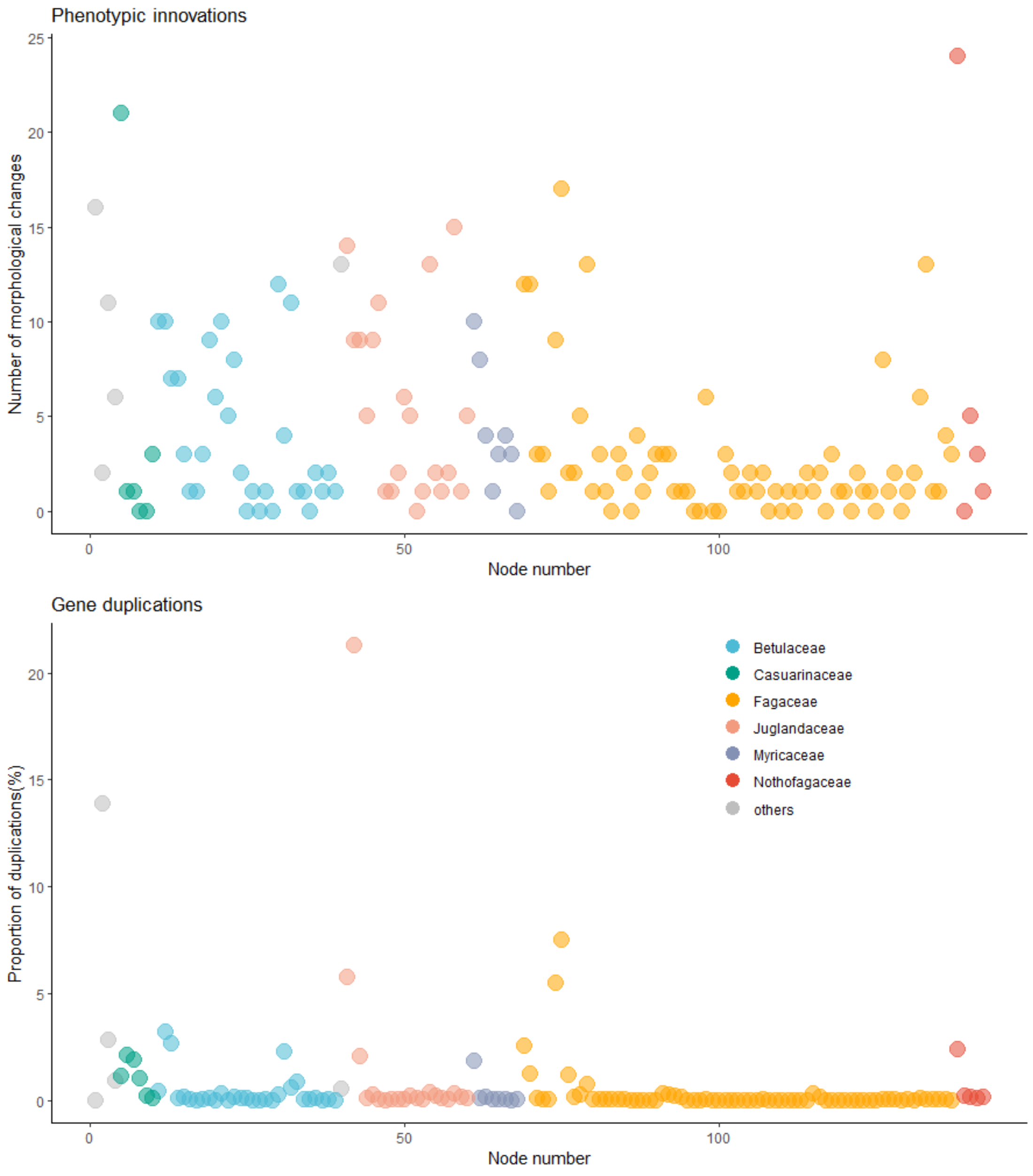

**Supplemental References:**


**Blanchard J, Wang H, Dilcher DL. 2016.** Fruits, seeds and flowers from the Bovay and Bolden clay pits (early Eocene Tallahatta Formation, Claiborne Group), northern Mississippi, USA. *Palaeontologia Electronica* 19:1–59.

**Crane PR, Manchester SR. 1982.** An extinct juglandaceous fruit from the Upper Palaeocene of southern England. *Botanical Journal of the Linnean Society* 85: 89–101.

**Crepet WL, Nixon KC. 1989.** Earliest Megafossil Evidence of Fagaceae: Phylogenetic and Biogeographic Implications. *American Journal of Botany* 76: 842–855.

**Elliott LL, Mindell Randal A, Stockey Ruth A. 2006.** *Beardia vancouverensis* gen. et sp. nov. (Juglandaceae): permineralized fruits from the Eocene of British Columbia. *American Journal of Botany* 93: 557–565.

**Friis EM. 1983.** Upper Cretaceous (Senonian) floral structures of juglandalean affinity containing Normapolles pollen. *Review of Palaeobotany and Palynology* 39: 161–188.

**Friis EM, Pedersen KR, Schönenberger J. 2003.** *Endressianthus*, a new Normapolles-Producing plant genus of Fagalean affinity from the Late Cretaceous of Portugal. *International Journal of Plant Sciences* 164: S201–S223.

**Friis EM, Pedersen KR, Schönenberger J. 2006.** Normapolles plants: a prominent component of the Cretaceous rosid diversification. *Plant Systematics and Evolution* 260: 107–140.

**Gandolfo MA, Nixon KC, Crepet WL, Grimaldi DA. 2018.** A late Cretaceous fagalean inflorescence preserved in amber from New Jersey. *American Journal of Botany* 105: 1424–1435.

**Grímsson F, Grimm GW, Zetter R, Denk T. 2016.** Cretaceous and Paleogene Fagaceae from North America and Greenland: evidence for a Late Cretaceous split between *Fagus* and the remaining Fagaceae. *Acta Palaeobotanica* 56: 247–305.

**Guerin G, Hill RS. 2003.** *Gymnostoma tasmanianum* sp. nov., a Fossil Casuarinaceae from the Early Oligocene of Little Rapid River, Tasmania, Australia. *International Journal of Plant Sciences* 164: 629–634.

**Herendeen PS, Crane PR, Drinnan AN. 1995.** Fagaceous flowers, fruits, and capsules from the Campanian (Late Cretaceous) of central Georgia, USA. *International Journal of Plant Sciences* 156: 93–116.

**Heřmanová Z, Kvaček J, Friis EM. 2011.** *Budvaricarpus serialis* Knobloch & Mai, An Unusual New Member of the Normapolles Complex from the Late Cretaceous of the Czech Republic. *International Journal of Plant Sciences* 172: 285–293.

**Hermsen EJ, Gandolfo MA. 2016.** Fruits of Juglandaceae from the Eocene of South America. *Systematic Botany* 41: 316–328.

**Herrera F, Manchester SR, Koll R, Jaramillo CA. 2014.** *Fruits of Oreomunnea (Juglandaceae) in the early Miocene of Panama*: Missouri Botanical Garden Press.

**Kodrul T, Krassilov V. 2005.** New juglandaceous fruit morphotype from the Palaeocene of Amur Province, Russian Far East. *Acta Palaeobotanica* 45: 139–144.

**Larson-Johnson K**. 2016. Phylogenetic investigation of the complex evolutionary history of dispersal mode and diversification rates across living and fossil Fagales. *New Phytologist* 209: 418–435.

**Liang XQ, Wilde V, Ferguson DK, Kvaček Z, Ablaev AG, Wang YF, Li CS. 2010.** *Comptonia naumannii* (Myricaceae) from the early Miocene of Weichang, China, and the palaeobiogeographical implication of the genus. *Review of Palaeobotany and Palynology* 163: 52–63.

**Manchester SR. 1987.** *The fossil history of the Juglandaceae.* Ph.D, Indiana University Bloomington.

**Manchester SR. 1991.** *Cruciptera*, a new Juglandaceous winged fruit from the Eocene and Oligocene of western North America. *Systematic Botany* 16: 715–725.

**Manchester SR. 2011.** Fruits of Ticodendraceae (Fagales) from the Eocene of Europe and North America. *International Journal of Plant Sciences* 172: 1179–1187.

**Manchester SR, Chen ZD. 1998.** A new genus of Coryloideae (Betulaceae) from the Paleocene of North America. *International Journal of Plant Sciences* **159**: 522–532.

**Manchester SR, Crane PR. 1983.** Attached leaves, inflorescences, and fruits of *Fagopsis*, an extinct genus of fagaceous affinity from the Oligocene Florissant Flora of Colorado, U.S.A. *American Journal of Botany* 70: 1147–1164.

**Manchester SR, Crane PR. 1987.** A new genus of Betulaceae from the Oligocene of western North America. *Botanical Gazette* 148: 263–273.

**Manchester SR, Dilcher DL. 1997.** Reproductive and vegetative morphology of Polyptera (Juglandaceae) from the Paleocene of Wyoming and Montana. *American Journal of Botany* 84: 649–663.

**Manchester Steven R, Pigg Kathleen B, Crane Peter R. 2004.** *Palaeocarpinus dakotensis* sp.n. (Betulaceae: Coryloideae) and Associated Staminate Catkins, Pollen, and Leaves from the Paleocene of North Dakota. *International Journal of Plant Sciences* **165**: 1135–1148.

**Schönenberge J, Pedersen KR, Friis EM. 2001.** Normapolles flowers of fagalean affinity from the Late Cretaceous of Portugal. *Plant Systematics and Evolution* 226: 205–230.

**Scriven LJ, Hill RS. 1995.** Macrofossil Casuarinaceae: their identification and the oldest macrofossil record, *Gymnostoma antiquum* sp. nov., from the late paleocene of New South Wales, Australia. *Australian Systematic Botany* 8: 1035–1053.

**Sims HJ, Herendeen PS, Crane PR. 1998.** New genus of fossil Fagaceae from the Santonian (Late Cretaceous) of central Georgia, U. S. A. *International Journal of Plant Sciences* 159: 391–404.

**Sims HJ, Herendeen PS, Lupia R, Christopher RA, Crane PR. 1999.** Fossil flowers with Normapolles pollen from the Upper Cretaceous of southeastern North America. *Review of Palaeobotany and Palynology* 106: 131–151.

**Smiley CJ, Huggins LM. 1981.** *Pseudofagus Idahoensis*, N. Gen. Et Sp. (Fagaceae) from the Miocene Clarkia Flora of Idaho. *American Journal of Botany* 68: 741–761.

**Takahashi M, Friis EM, Herendeen PS, Crane PR. 2008.** Fossil Flowers of Fagales from the Kamikitaba Locality (Early Coniacian; Late Cretaceous) of Northeastern Japan. *International Journal of Plant Sciences* 169: 899–907.

**Wang H, Blanchard J, Dilcher DL. 2013.** Fruits, seeds, and flowers from the Warman clay pit (middle Eocene Claiborne Group), western Tennessee, USA. *Palaeontologia Electronica* 16, 73p.

**Wheeler EA, Baas P, Manchester SR**. 2022. Wood anatomy of modern and fossil Fagales in relation to phylogenetic hypotheses, familial classification, and patterns of character evolution. *International Journal of Plant Sciences* 183: 61–86.

**Wilde V, Frankenhäuser H, Lenz OK. 2021.** A myricaceous male inflorescence with pollen in situ from the middle Eocene of Europe. *Palaeobiodiversity and Palaeoenvironments* 101: 873–883.

**Wilf P, Nixon KC, Gandolfo MA, Cuneo NR. 2019.** Eocene Fagaceae from Patagonia and Gondwanan legacy in Asian rainforests. *Science* 364: 972.

**Wing SL, Hickey LJ. 1984.** The *Platycarya* perplex and the evolution of the Juglandaceae. *American Journal of Botany* 71: 388–411.

**Zamaloa MC, Gandolfo MA, González CC, Romero EJ, Cúneo NR, Wilf P. 2006.** Casuarinaceae from the Eocene of Patagonia, Argentina. *International Journal of Plant Sciences* 167: 1279–1289.

**Zhang Q, Ree RH, Salamin N, Xing Y, Silvestro D**. 2022. Fossil-informed models reveal a boreotropical origin and divergent evolutionary trajectories in the walnut family (Juglandaceae). *Systematic Biology* 71: 242–258.

**Zhang YZ, Deng T, Sun L, Landis JB, Moore MJ, Wang HC, Wang YH, Hao XJ, Chen JJ, Li SH, et al. 2020.** Phylogenetic patterns suggest frequent multiple origins of secondary metabolites across the seed plant “tree of life”. *National Science Review* 8, nwaa105.
